# p53 protein abundance is a therapeutic window across TP53 mutant cancers and is targetable with proximity inducing small molecules

**DOI:** 10.1101/2024.07.27.605429

**Authors:** Ananthan Sadagopan, Nicholas Garaffo, Heng-Jui Chang, Stuart L. Schreiber, Matthew Meyerson, William J. Gibson

**Affiliations:** Broad Institute of MIT and Harvard, Cambridge, Massachusetts 02142, USA; Department of Medical Oncology, Dana-Farber Cancer Institute, Boston, Massachusetts 02215, USA; Division of Medical Sciences, Harvard Medical School, Boston, Massachusetts 02115, USA; Department of Chemistry and Chemical Biology, Harvard University, Cambridge, Massachusetts 02138, USA; Arena Bioworks, Cambridge, Massachusetts 02141, USA; Department of Medicine, Harvard Medical School, Boston, Massachusetts 02115, USA

## Abstract

*TP53* mutant cancers are associated with approximately half of cancer deaths. The most common mechanism of p53 inactivation involves missense mutations. Such mutations in *TP53* result in a robust upregulation of the p53 protein. Here, we demonstrate an induced proximity approach to selectively kill *TP53* mutant cells. This approach uses the increased abundance of p53 protein in *TP53* mutant cancer cells to concentrate toxic molecules in these cells. We demonstrate the first generalizable strategy using a small molecule to selectively kill *TP53* mutant cells. This molecule binds the Y220C mutant of p53 and concentrates a PLK1 inhibitor in cells harboring *TP53* Y220C mutations. Together, these data demonstrate that the abundance of p53 protein provides a therapeutic window for *TP53* missense mutant cancers that can be translated into a cell death signal using proximity-inducing small molecules.

## Introduction

*TP53* is the most mutated gene in human cancer (36% [PCAWG], 37% [TCGA], 39% [AACR GENIE])^1–3^. *TP53* mutations have been shown to induce resistance to cytotoxic chemotherapy and are associated with a worse prognosis across cancers^4–6^. In total, *TP53* mutant cancers are associated with ∼46% of cancer deaths (Supplementary Note, analysis across cBioPortal^7,8^). In the past, attempts have been made to therapeutically target *TP53* mutant cancers in several ways^9^: delivery of wild-type *TP53* genes to cells with gene therapy or mRNA^10,11^, immunotherapy^12–14^, or small molecule refolders of p53^15,16^. Attempts to enact these strategies have invariably failed, and the activity of purported mutant-agnostic p53 refolders has consistently been shown to be off-target^17,18^. Furthermore, small molecule refolding of p53 may not be possible for contact mutants of p53 that specifically mutate residues that interact with the negatively charged phosphate backbone of DNA. New generalizable strategies are needed to specifically kill cancer cells harboring these mutations.

In contrast to mutations of most tumor suppressor genes, such as *APC, RB1*, or *PTEN, TP53* mutations are typically missense mutations. These mutations act as dominant negative mutations that disallow DNA binding at consensus sequences, and therefore poison the tetramer^19^. Because they act as dominant negatives, they function as a shorter evolutionary path to inactivating most p53 function within the cells that harbor these mutations^20–22^. p53 typically acts as a central signal integrator for various cellular stress signals such as hypoxia, DNA damage, or excessive oncogenic signaling^23,24^. When these mutations occur, the mechanisms that normally sense the above cellular stressors are typically intact, and jointly work to activate p53 by increasing its abundance and promoting its nuclear translocation. Meanwhile the mutant p53 that accumulates is impotent to drive the transcription of its own destroyer, MDM2. The negative feedback loop that usually serves to tightly regulate p53 protein is broken and the half-life of p53 stretches from its typical ∼5-20 minutes to several hours. Thus, p53 protein accumulates in *TP53* missense mutant cells^25–29^.

Here, we develop small molecules that utilize this characteristic of *TP53* missense mutants, accumulation of p53 protein, to selectively kill cancer cells characterized by high levels of mutant p53.

## Results

### *TP53* mutant cells are not associated with synthetic lethality for any genes nor enriched for compound sensitivity

Large scale functional genomic studies have recently been undertaken to systematically determine the genetic vulnerabilities of nearly a thousand cancer cell lines^30,31^. We sought to test the hypothesis that there exist synthetic lethal interactions with *TP53* mutant cancer cells. We performed a genome-wide analysis of CRISPR dependencies in cells with *TP53* mutations vs. wild-type (WT) cell lines. Unfortunately, there is no genetic dependency that is enriched in cancer cells with *TP53* loss-of-function mutations (Figure 1A; no genes with (Mean DepScore)_Mut_< -0.5 and Δ(Mean DepScore)_Mut-WT_ < -0.2, consistent with a prior report^22^), nor any profiled small molecules selectively killing *TP53* mutant cells (Figure S1, no molecules >2-fold more toxic on average to p53^Mut^ vs. p53^WT^ cells in PRISM repurposing screen or with Δ(Mean AUC)_Mut-WT_ < -0.5 in CTD2 screen). One might instead hope that a subset of proteins are enriched in cells with *TP53* mutations, particularly cell surface proteins that might be amenable to targeting with a number of effector modalities such as CAR-T or antibody-drug conjugates. Unfortunately, no such cell-surface proteins are enriched in *TP53* mutant cancers (Figure 1B, Figure S1).

**Fig. 1.**
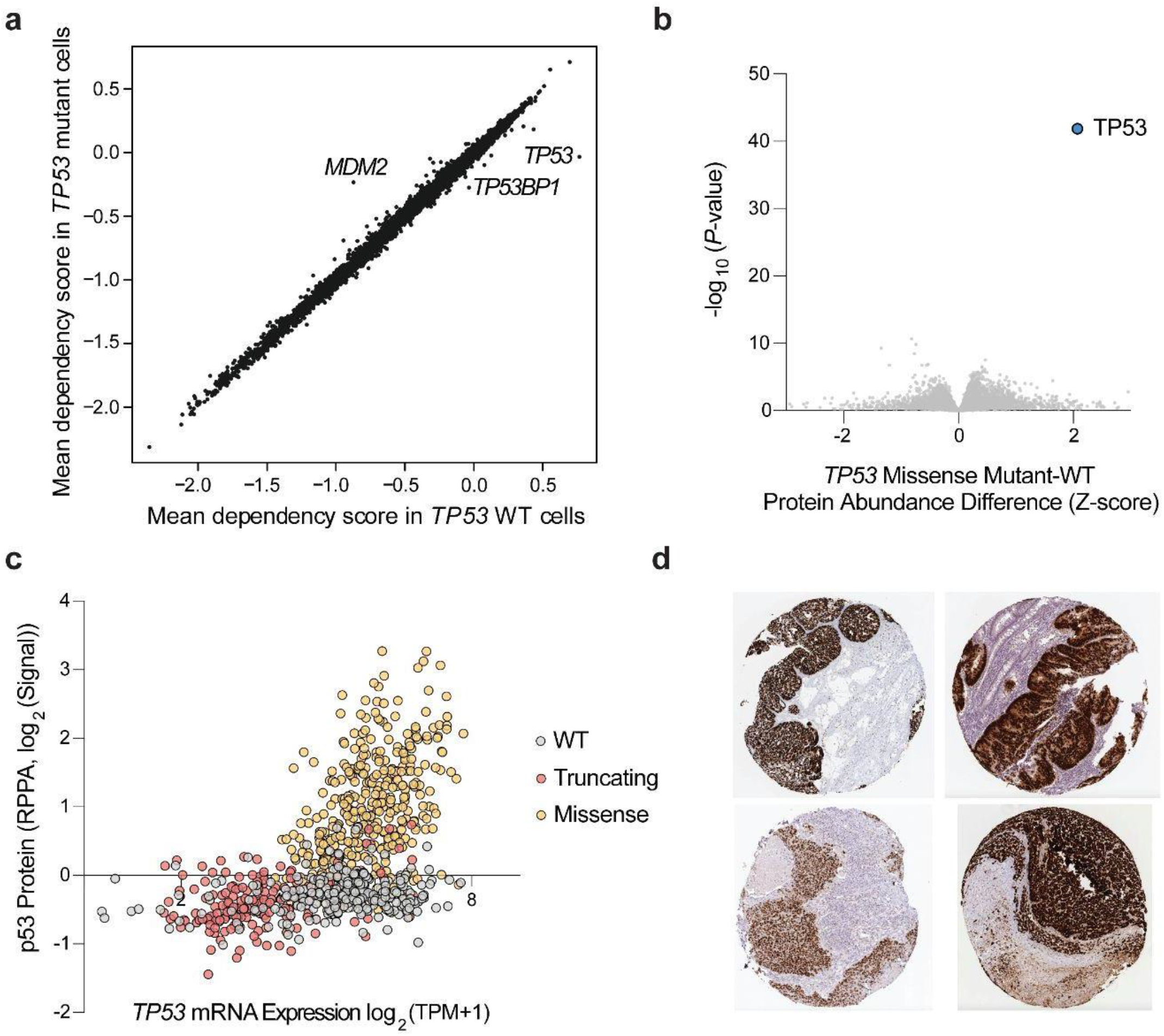
p53 protein abundance is the only genetic or proteomic distinction between *TP53* mutant and WT cancer cells. **a**, Mean CRISPR dependency scores for genes across DepMap in *TP53* wild-type (x-axis) vs. *TP53* mutant cells (y-axis). **b**, Proteome-wide volcano plot of proteins that are enriched or depleted in *TP53* mutant cells versus *TP53* wild-type cells across DepMap (difference in mean Z-scored quantitative proteomics; t-test). **c**, *TP53* mRNA expression (log_2_(TPM+1)) vs. p53 protein abundance (log_2_(Signal) from RPPA) across DepMap. Cell lines are colored by *TP53* mutation status (wild-type: gray, truncating: red, missense: yellow). **d**, representative immunohistochemistry of p53 in Human Protein Atlas^61^ from resected tumors.

### *TP53* mutant cells show significant upregulation of p53 protein

Quantitative proteomics of the Cancer Cell Line Encyclopedia (CCLE)^32^ shows that the only protein whose abundance is substantially and significantly increased in *TP53* mutant cancers is p53 itself (Figure 1B, only protein with Δ(mean Z-scored Protein Abundance)_Mut-WT_ > 2 and *q*<0.05). p53 protein is expressed at a barely detectable level in *TP53*^WT^ cancer cells but is elevated in *TP53* missense mutant cells (Figure 1C, >2-fold increase in mean reverse phase protein array signal). Correspondingly, immunohistochemistry of p53 in tumor tissues shows abundant staining in cancer cells, but not adjacent normal tissue (Figure 1D). Taken together, these data demonstrate that the only genetic dependency or proteomic difference between *TP53* mutant cells and WT cells, detected to date, is the overabundance of p53 protein in *TP53* mutant cells.

The increased abundance of the p53 protein in cancer cells with *TP53* missense mutations provides a unique therapeutic opportunity: a p53 concentration-dependent toxin could selectively kill *TP53* mutant cancer cells in a way that is generalizable to multiple p53 missense mutations.

### Bifunctional small molecules kill cells based on p53 protein abundance

We therefore hypothesized that if the abundance of p53 protein could be translated into a proportionate cell death signal, *TP53* missense mutant cells could be selectively killed. A recently developed method has shown that bifunctional small molecules that bind to an overexpressed intracellular protein can concentrate toxic molecules in cells overexpressing said protein^33^.

Unfortunately, no high affinity ligands for WT p53 exist. In the absence of such ligands, we used the Halo-tag as a surrogate for a small molecule binder of p53. Mutant p53^R273H^ was stably expressed in 293T cells fused to a Halo-tag and mCherry for visualization. The fusion protein showed nuclear expression similar to native p53 (Figure S2).

To leverage the abundance of p53 protein for therapeutic purposes, a bifunctional molecule would ideally be designed such that its binding partner is not only less abundant than over-expressed mutant p53 protein but also highly essential for cell survival. To determine which cytotoxin to append to a potential bifunctional molecule, we compared CRISPR gene essentiality scores across Dependency Map^34^ and absolute protein abundance profiled in OpenCell^35^ (Figure 2A). We identified five targets with high essentiality and low abundance: DNA2, WEE1, LRR1, PRELID1, and PLK1 (Figure 2A, see **Methods**). Of these, WEE1 and PLK1 have ligands previously used to synthesize bifunctional small molecules for targeted protein degradation^36–38^. We functionalized WEE1 inhibitor adavosertib^39^ (IC_50_ = 5.2 nM for cell-free WEE1 inhibition) and an analog of PLK1 inhibitor BI-2536^40^ (IC_50_ = 0.83 nM for cell-free PLK1 inhibition), both of which display potent antiproliferative effects across DepMap (adavosertib median IC_50_: 348 nM, BI-2536 median IC_50_: 11.8 nM; Figure S2), with a Halo-tag ligand and PEG2 linker (Figure 2B, Figure S3, yielding Halo-PEG2-adavosertib and Halo-PEG2-BI2536).

**Fig. 2.**
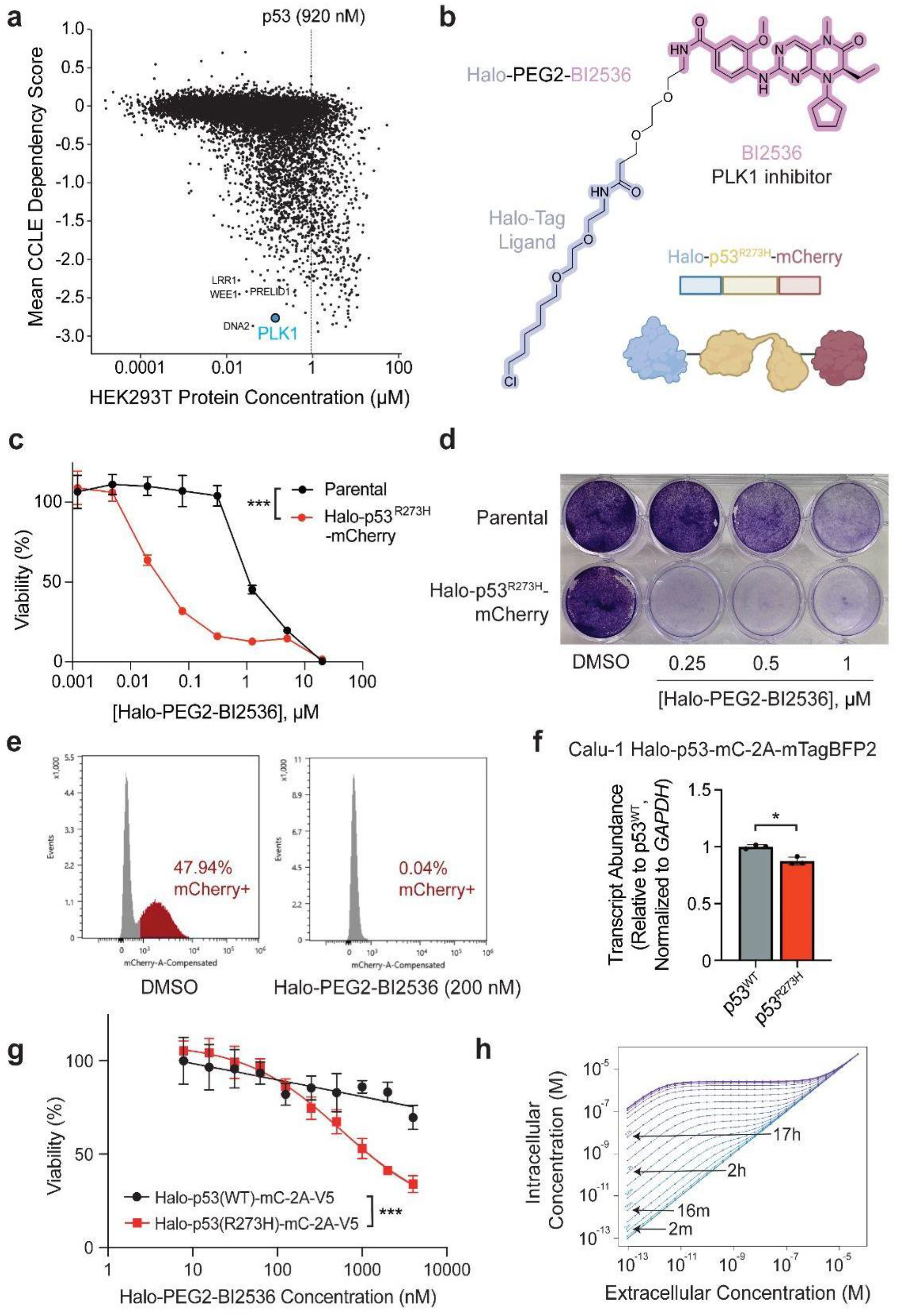
Bifunctional molecules liganding p53 fusion proteins selectively inhibit proliferation of p53 mutant cells. **a**, Protein concentration in HEK293T cells profiled in OpenCell vs. mean CRISPR dependency score across DepMap for all genes/protein products. **b**, Structure of small molecule (Halo-PEG2-BI2536) and fusion protein construct (Halo-p53^R273H^(FL)-mCherry) used in experiments. **c**, Cell viability of 293T cells stably expressing Halo-p53^R273H^(FL)-mCherry versus parental 293T cells treated with Halo-PEG2-BI2536, measured by cell-titer-glo after 5 days. \*\*\**P*<0.001 by two-way ANOVA. **d**, Crystal violet staining of 293T cells stably expressing Halo-p53^R273H^(FL)-mCherry versus parental 293T cells treated with Halo-PEG2-BI2536, analyzed after 4 days. **e**, Competition experiment in which a 1:1 mixture of Halo-p53^R273H^(FL)-mCherry:parental 293T cells were treated with Halo-PEG2-BI2536 (200 nM) for 7 days and analyzed for mCherry expression. **f**, Calu-1 cell lines stably expressing Halo-p53^WT^(FL)-mCherry-2A-mTagBFP2-V5 and Halo-p53^R273H^(FL)-mCherry-2A-mTagBFP2-V5 were established and analyzed by RT-qPCR for expression of the transgene. Transcript abundance is normalized to GAPDH expression using the 2^−ΔΔCt^ method and is plotted relative to Calu-1 cells stably expressing Halo-p53^WT^(FL)-mCherry-2A-mTagBFP2-V5. \**P*<0.05 by heteroscedastic t-test. **g**, Cell viability of Calu-1 cells stably expressing Halo-p53^WT^(FL)-mCherry-2A-mTagBFP2-V5 or Halo-p53^R273H^(FL)-mCherry-2A-mTagBFP2-V5 treated with Halo-PEG2-BI2536, measured by cell-titer-glo after 4 days. \*\*\**P*<0.001 by two-way ANOVA. **h**, Modeling of final intracellular concentration at steady-state for a freely diffusing bifunctional compound binding to p53 (see **Methods**), based on p53 protein half-life and initial extracellular concentration (held constant); purple indicates a longer half-life.

Halo-PEG2-BI2536 inhibited proliferation of 293T cells stably expressing Halo-p53^R273H^(FL)-mCherry at doses between 20 nM - 500 nM, with little impact on the proliferation of wild-type 293T cells (Figure 2C-D, Halo-p53^R273H^(FL)-mCherry 293T IC_50_: 23 nM, WT 293T IC_50_: 1143 nM, ratio = 49.7, *P*<0.001). 7-day competition of 293T cells stably expressing Halo-p53^R273H^(FL)-mCherry with wild-type 293T cells in the presence of Halo-PEG2-BI2536 resulted in a 1200-fold decline in the population of mCherry+ cells compared to DMSO control (47.94% vs. 0.04% at 200 nM; Figure 2E). These results were reproducible in a p53 null (*TP53* homozygous deletion) Calu-1 cell line stably expressing Halo-p53^R273H^(FL)-mCherry (Figure S2, EC_50_ = 13 nM from competition assay). Furthermore, treatment of Halo-p53^R273H^(FL)-mCherry 293T cells with low doses of Halo-PEG2-BI2536 resulted in selection for low expressors of the fusion protein (Figure S2). Halo-PEG2-adavosertib also selectively inhibited proliferation of Halo-p53^R273H^(FL)-mCherry 293T cells in the competition assay at doses between 100 nM - 1 μM (Figure S3, EC_50_ = 41 nM).

To assess the therapeutic window obtained by this strategy, we established Calu-1 cell lines stably expressing Halo-p53(FL)-mCherry-2A-mTagBFP2-V5 using either p53^WT^ or p53^R273H^. We confirmed that these cell lines displayed approximately equal expression of the transgene by RT-qPCR for the transcript (Figure 2F, in fact WT had slightly greater expression: Mean [*TP53*^WT^ mRNA] / [*TP53*^R273H^ mRNA] = 1.14, *P* < 0.05). Halo-PEG2-BI2536 was at least an order of magnitude more detrimental to the growth of p53^R273H^ cells (IC_50_ = 402 nM) compared to p53^WT^ cells (IC_50_ > 4 μM) after 4 days (Figure 2G, *P*<0.001). Prior reports have suggested that the enhanced efficacy of similar bifunctional compounds can occur due to accumulation of small molecules in cells expressing high concentrations of the protein target^33^. We therefore modeled reaction-diffusion kinetics of Halo-PEG2-BI2536 using ordinary differential equations (see **Methods**) to gain more insights into the enhanced toxicity observed in the presence of a ligandable p53 fusion protein. Previously reported increases in p53 protein half-life upon mutation greatly influence the expected accumulation of the bifunctional small molecule^28,29^. In particular, an increase in half-life from the WT p53 half-life of 16 minutes to a mutant p53 half-life of 17 hours is expected to increase final intracellular concentration of the small molecule at steady-state by 2-3 orders of magnitude depending on the initial concentration (Figure 2H). Together, these data suggest that if small molecule ligands of p53 were developed, the high protein abundance of its mutated forms would allow for differential cell killing.

### Bifunctional small molecules selectively kill p53^Y220C^ mutant cells

To assess the impact of bifunctional molecules containing p53-binding and cytotoxic moieties in an endogenous cellular context, we synthesized molecules that bind to p53^Y220C^. This mutant indirectly inhibits DNA binding activity through the loss of DNA-binding domain thermal stability^41^. The removal of a bulky tyrosine residue, creating a pocket, and its replacement with a ligandable cysteine makes it an attractive target for small molecule re-folders meant to restore transcriptional activity^42–47^, as well as to bifunctional cytotoxic strategies. We initially used acrylamide KG5 as the p53^Y220C^ binder in a PLK1-directed bifunctional; KG5 is a carbazole-based covalent fragment liganding >95% of p53^Y220C^ at 10 μM^45^. We synthesized KG5-PEG4-BI2536 and did not observe any difference in cell survival between 293T cells stably expressing Halo-p53^Y220C^ΔTAD-mCherry (transactivation domain deleted to remove confounding protein stabilization effects) and wild-type 293T cells in an 8-day competition assay (Figure 3D, Figure S3, ratio of mCherry+% for 1 μM KG5-PEG4-BI2536/DMSO = 0.998).

**Fig. 3.**
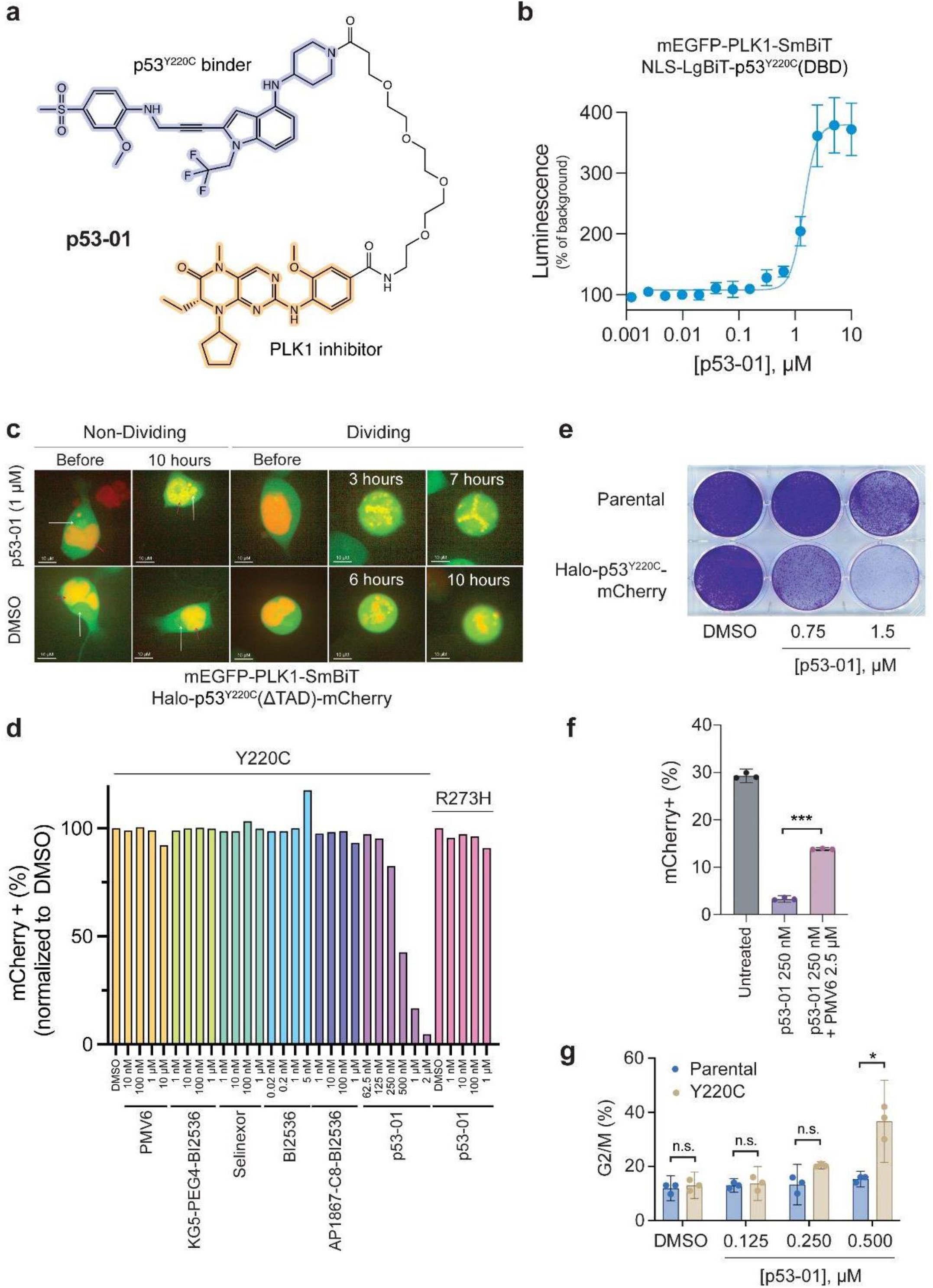
p53^Y220C^-PLK1 bifunctional molecules selectively inhibit proliferation of p53^Y220C^ mutant cells. **a**, Structure of p53^Y220C^-PLK1 bifunctional compound (PMV6-PEG4-BI2536, p53-01) used in experiments. **b**, Nanoluciferase signal one day after dose titration of p53-01 in 293T cells co-transfected with NLS-LgBiT-p53^Y220C^(DBD) and mEGFP-PLK1-SmBiT. **c**, Live cell imaging of 293T cells co-transfected with Halo-p53^Y220C^ΔTAD-mCherry and mEGFP-PLK1-SmBiT and treated with p53-01. White arrow indicates the extranuclear region containing PLK1; red arrow indicates the nucleus. Dividing and non-dividing cells shown. **d**, Competition of 293T cells stably expressing Halo-p53^Y220C^ΔTAD-mCherry vs. parental 293T cells in the presence of various compounds (PMV6, KG5-PEG4-BI2536, selinexor, BI2536, AP1867-C8-BI2536, p53-01). Competition of 293T cells stably expressing Halo-p53^R273H^(FL)-mCherry vs. parental 293T cells in the presence of p53-01 is also shown on the right. mCherry+ percentage on day 8 of competition normalized to that of DMSO-treated cells. **e**, Crystal violet staining of 293T cells stably expressing Halo-p53^Y220C^ΔTAD-mCherry versus parental 293T cells treated with p53-01, analyzed after 5 days. **f**, Competition of 293T cells stably expressing Halo-p53^Y220C^ΔTAD-mCherry vs. parental 293T cells in the presence of p53-01 (250 nM) alone or in combination with PMV6 (2.5 μM). \**P*<0.001 by heteroscedastic t-test. **g**, Percentage of cells in G2/M phase of cell cycle following treatment of Halo-p53^Y220C^ΔTAD-mCherry or parental 293T cells with p53-01 after 16 hours. \**P*<0.05; n.s. not significant by heteroscedastic t-test.

We surveyed the patent literature and identified a p53^Y220C^ binding pharmacophore consisting of an *o*-anisidine moiety linked to an indole and piperidine (US20230024905A1, US10138219B2, WO2021262483A1, WO2021061643A1, WO2022213975A1, WO2023016434A1). These binders are exemplified by PMV6, a compound identified in a structure-activity relationship series by PMV Pharma, with close similarity to the recently disclosed structure of the clinical compound, rezatapopt, which has a K_d_ ∼ 2.5 nM for p53^Y220C^ (PMV6, p. 96: US20230024905A1)^48^. PMV6 induces an 8°C thermal shift of p53^Y220C^ (p. 21: US20230024905A1). The co-crystal structure with p53^Y220C^ showed that the compound binds to a shallow groove created by the Y220C mutation, with the piperidine facing the solvent. To construct a bifunctional molecule containing PMV6 and BI-2536, we functionalized PMV6 at the solvent-exposed piperidine, which was functionalized in other analogs that continued to bind p53^Y220C^ via thermal shift assay (Compound 7 & Compound 10, p. 21&96: US20230024905A1).

We synthesized PMV6-PEG4-BI2536 (p53-01, Figure 3A) and confirmed that it induces ternary complex formation between p53^Y220C^ and PLK1 using a NanoBiT assay in 293T cells co-transfected with mEGFP-PLK1-SmBiT and NLS-LgBiT-p53^Y220C^(DBD) (Figure 3B, EC_50_ = 1.4 μM). Live cell imaging of 293T cells co-transfected with mEGFP-PLK1-SmBiT and Halo-p53^Y220C^ΔTAD-mCherry revealed diffuse PLK1 localization throughout the cell, with strong enrichment in an extranuclear region, presumably the centrosome, while p53^Y220C^ was constitutively diffusely nuclear. As an alternative method to demonstrate ternary complex formation, we assessed protein colocalization upon compound treatment^49–52^. In non-mitotic cells, PLK1 became enriched in the nucleus colocalizing with p53^Y220C^, while p53^Y220C^ also entered the centrosome colocalizing with PLK1. In mitotic cells, p53^Y220C^ is normally localized to chromatin while PLK1 is localized elsewhere in the cell. However, upon compound treatment, PLK1 becomes colocalized with p53^Y220C^ on chromatin (Figure 3C). We speculate this mislocalization could disrupt the function of PLK1 as a mitotic kinase beyond simple steric blockade of its active site.

In competition between Halo-p53^Y220C^ΔTAD-mCherry and parental 293T cells, we observed strong selection against p53^Y220C^+ cells when p53-01 was dosed between 250 nM - 2 μM by day 8 (21.5-fold decline in the mCherry+% at 2 μM, EC_50_ = 443 nM). We did not observe any activity from non-functionalized binders or control compounds, including PMV6, BI2536, AP1867-C8-BI2536, or selinexor (no mCherry+% declines >1.1-fold observed). We also did not observe any activity of p53-01 in either 293T or Calu-1 cells expressing Halo-p53^R273H^(FL)-mCherry (Figure 3D-E, Figure S2; no mCherry+% declines >1.1-fold observed). An analogous bifunctional molecule constructed with a WEE1 inhibitor (PMV6-PEG4-adavosertib) was unable to selectively inhibit proliferation of p53^Y220C^+ cells (Figure S3). To confirm that the observed selective proliferation effect was mediated by compound binding to p53^Y220C^, we repeated the Halo-p53^Y220C^ΔTAD-mCherry/WT competition experiment using PMV6 to compete off p53-01, and observed partial rescue (p53-01 [250 nM]: 8.9-fold decline in mCherry+%, p53-01 [250 nM] + PMV6 [2.5 μM]: 2.1-fold decline in mCherry+%, *P*<0.001) (Figure 3F). Of note, PMV6 at high concentrations (≥ 5 μM) was toxic even in wild-type 293T cells, making it challenging to perform experiments with higher doses likely required for complete rescue.

PLK1 activity is critical for mitotic progression, and its inhibitor, BI-2536, has been previously shown to induce G2/M arrest^53^. We monitored for induction of G2/M arrest upon p53-01 treatment (500 nM) and observed significant induction in Halo-p53^Y220C^ΔTAD-mCherry 293T cells but not WT cells (Figure 3G, G2/M% Halo-p53^Y220C^ΔTAD-mCherry 293T: 36.7%, WT 293T: 15.3%, *P*<0.05). Together, these data indicate that a small molecule binding to p53^Y220C^ and PLK1 can selectively inhibit proliferation of cells expressing *TP53*^Y220C^ due to enhanced PLK1 inhibition, leading to mitotic arrest.

### p53^Y220C^-PLK1 bifunctional small molecules do not function through p53 reactivation and are active in endogenous settings

We next analyzed whether the mechanism of action for p53-01 is distinct from that for PMV6, a reactivator of p53^Y220C^. We performed RNA sequencing of Huh7 hepatocellular carcinoma cells (homozygous p53^Y220C/Y220C^) treated with p53-01 (4 μM) and PMV6 (4 μM) for one day. We observed strong induction of a p53 signature by PMV6, including >10-fold upregulation of p53 targets such as *MDM2, CDKN1A*, and *GDF15* and downregulation of genes essential for proliferating cells such as *TOP2A*. Manual inspection of a set of high confidence p53 transcriptional targets, and gene set enrichment analysis (GSEA) of a larger group of consensus p53 targets confirmed strong upregulation of these genes by PMV6 (NES = 2.68) but not p53-01 (NES = 1.33) (Figure 4A-C, Figure S4).

**Fig. 4.**
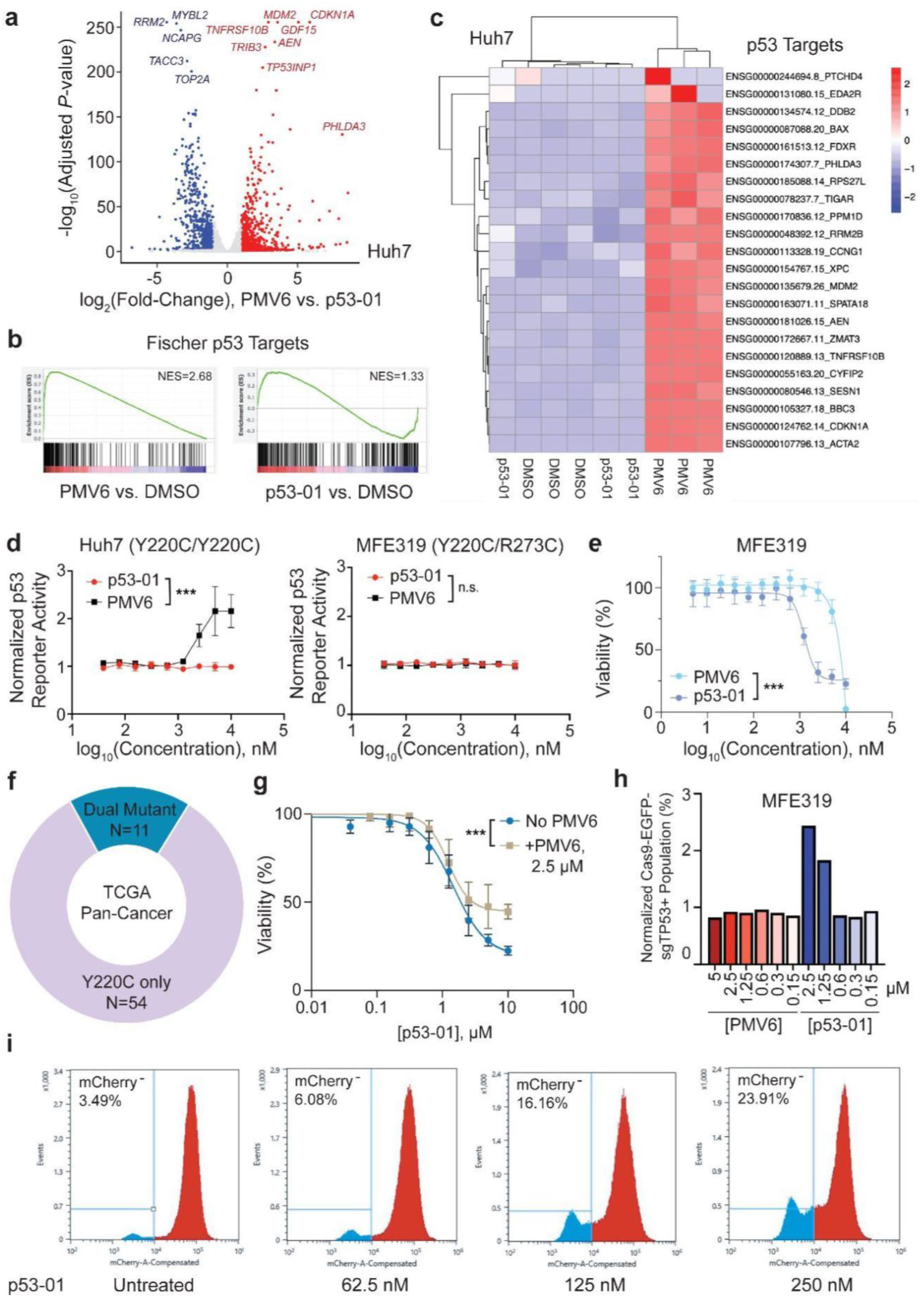
p53^Y220C^ -PLK1 bifunctionals are active in endogenous settings and do not reactivate p53. **a**, Huh7 cells homozygous for the p53^Y220C^ mutation were treated with PMV6 (4 μM) or p53-01 (4 μM) for 24 hours and subject to RNA sequencing. log_2_(fold-changes) in transcript abundance and log_10_(adjusted p-values) computed by DESeq2 are plotted. Top differentially expressed genes are labeled. **b**, Pre-ranked GSEA analysis using DESeq2 t-statistic as rank metric for PMV6 vs. DMSO (left) and p53-01 vs. DMSO (right) comparisons on MSigDB Fisher direct p53 targets gene set. Normalized enrichment scores are listed. **c**, Gene expression analysis of p53 targets from high confidence p53 target set. Samples are arranged in columns by hierarchical clustering (Euclidean distance), and rows scaled to z-scores (colors). **d**, p53 luciferase reporter activity following PMV6 or p53-01 treatment of Huh7 (left) or MFE319 (right) cell lines for 16 hours. Luciferase signal is normalized to untreated cells. \*\*\**P*<0.001; n.s. not significant by two-way ANOVA. **e**, Cell viability of MFE319 cells treated with PMV6 or p53-01, measured by cell-titer-glo after 4 days. \*\*\**P*<0.001 by two-way ANOVA. **f**, Donut plot of cancers in TCGA Pan-Cancer Atlas studies in cBioPortal with p53^Y220C^ mutations with/without additional p53 mutations. **g**, Cell viability of MFE319 cells treated with p53-01 alone or in combination with PMV6 (2.5 μM), measured by cell-titer-glo after 4 days. \*\*\**P*<0.001 by two-way ANOVA. **h**, Competition of MFE319 cells stably expressing sgTP53-Cas9-EGFP vs. parental MFE319 cells in the presence of p53-01 or PMV6. EGFP+ percentage on day 9 of competition normalized to that of DMSO-treated cells. **i**, Halo-p53^Y220C^ΔTAD-mCherry 293T cells were cultured in the presence of p53-01 for 3 weeks. The percentage of mCherry negative cells was analyzed by flow cytometry.

We subsequently established p53^Y220C^+ cell lines stably expressing a p53 luciferase reporter^54^. In Huh7 cells (p53^Y220C/Y220C^), PMV6 was able to induce activity of the reporter at doses of 1.25 - 10 μM (EC_50_ = 2.4 μM), while p53-01 had no such effect. We repeated the p53 reporter experiments in the p53^Y220C^+ endometrial cancer cell line MFE319, where no induction of luciferase activity for either compound was observed (Figure 4D). Note that MFE319 has both p53^Y220C^ and p53^R273C^ mutations reported in the CCLE dataset^31^. Analysis of paired RNA sequencing reads in this cell line revealed that the mutations were *in trans* (Figure S4). Notably, p53-01 was substantially more toxic to MFE319 cells than was PMV6 (p53-01 IC_50_: 1.26 μM, PMV6 IC_50_: 7.55 μM, *P*<0.001) (Figure 4E). These data suggest that the presence of an additional dominant negative p53 mutation imparts resistance to p53^Y220C^ re-folders, presumably due to poisoning of the tetramer even in the presence of correctly functioning p53^Y220C^ proteins. The strategy presented in this paper, using p53^Y220C^ to concentrate a toxin in cells, is not impacted by these additional p53 mutations, immediately resulting in a ∼20% greater scope compared to p53^Y220C^ reactivators (Figure 4F, as determined by analysis of p53^Y220C^ mutant cancers in TCGA^55^ with additional *TP53* mutations).

Lastly, we wanted to assess whether the effects of p53-01 in MFE319 were dependent on its interaction with p53^Y220C^. We were able to partially compete off the decline in cell viability induced by p53-01 in MFE319 using PMV6 (Figure 4G, *P*<0.001, max response for p53-01 alone: 22.5% viable, p53-01 + PMV6 [2.5 μM]: 44.6% viable). Additionally, we established MFE319 cells stably expressing sgTP53-Cas9-EGFP. In competition with wild-type MFE319 cells, p53-01 was able to enrich an EGFP+ population at doses between 1.25 μM - 2.5 μM (Figure 4H, 2.44-fold enrichment of EGFP+% at 2.5 μM). As further evidence of on-target activity, we cultured Halo-p53^Y220C^ΔTAD-mCherry 293T cells for 3 weeks in the presence of low doses of p53-01 (≤ 250 nM) and observed the expansion of an mCherry negative population (Figure 4I, 6.9-fold expansion of mCherry-% at 250 nM). Altogether, these data indicate that p53-01 has activity in cell lines harboring endogenous TP53^Y220C^ mutations through a non-transcriptional mechanism distinct from PMV6, and its activity is dependent on its interaction with p53^Y220C^.

### Linker optimization improves the efficacy of p53^Y220C^-PLK1 bifunctional small molecules

Lastly, we created a library of nine PMV6-BI2536 bifunctional compounds with varying linker lengths (4.33 - 22.22 Å). We assessed their efficacy in competition assays between wild-type and Halo-p53^Y220C^ΔTAD-mCherry 293T cells (Figure 5A). We observed no consistent effect of linker length on compound efficacy in general; although, shorter alkyl linkers outperformed longer alkyl linkers. PEG(2,4,6) linkers all performed similarly. The most potent compound: PMV6-C3-BI2536 (β-alanine linker), was over 1.5 orders of magnitude more potent than p53-01 in the competition assay (PMV6-C3-BI2536 EC_50_: 25 nM, p53-01 EC_50_: 850 nM) (Figure 5B-C). These data indicate that substantial improvements to the efficacy of p53^Y220C^-PLK1 bifunctional compounds are possible going forward.

**Fig. 5.**
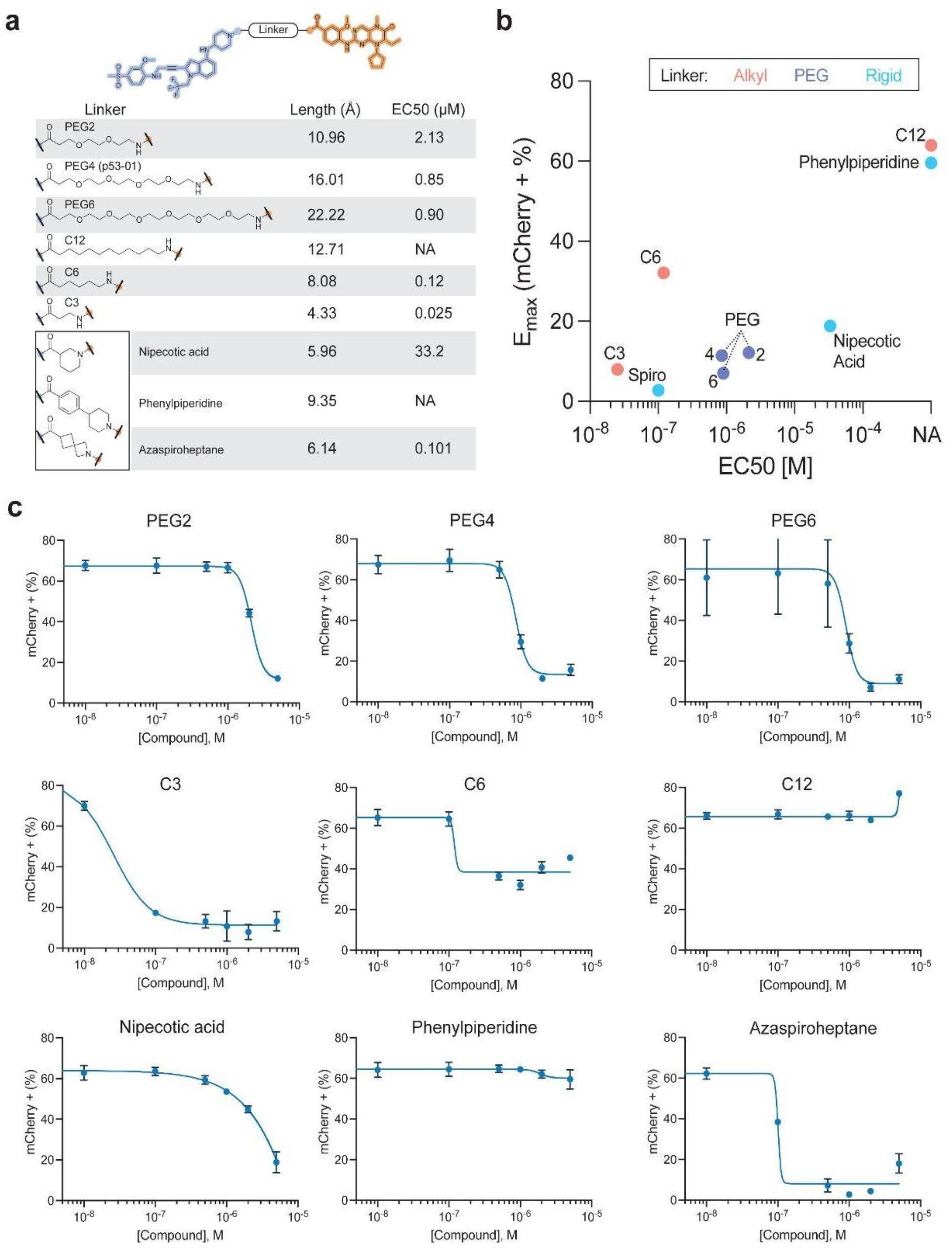
Linker optimization yields more potent p53^Y220C^ -PLK1 bifunctionals. **a**, Summary of linkers tested; linker length and EC_50_ from Halo-p53^Y220C^ΔTAD-mCherry vs. parental 293T competition is listed. Competition assays were analyzed on day 10. NA indicates the EC_50_ could not be computed due to lack of potency. **b**, EC_50_ from the competition experiment is plotted against E_max_ (calculated as the lowest percentage of mCherry+ cells in the competition assay observed at a tested dose). **c**, EC_50_ dose curves for individual compounds from the competition assay.

## Discussion

*TP53* mutations remain the dominant mutation associated with death from human cancers^5^. While there have been prior reports of mutation-agnostic small molecule re-folders of p53, these molecules have later been found to lack the desired activity^17,18^. To our knowledge, the molecules reported in this manuscript represent the first generalizable strategy using small molecules to specifically kill cancer cells bearing *TP53* mutations. Our strategy reveals one critical insight: that p53-targeted drugs need not restore native p53 function, but can instead use differential p53 protein abundance or mutant specific ligands to bring about a *TP53* mutant selective therapy. Here, we use the high intracellular concentration of the missense mutant p53 protein to induce cancer-selective cell death.

The molecules in this report are far from pharmacologically optimized. We anticipate that more potent and selective molecules may be achieved in the future. Because the bivalent compounds presented here include a non-specific small molecule toxin, they are able to induce toxicity in cells that do not bear *TP53* mutations when administered at sufficiently high doses. Future generations of molecules may have no toxicity in the absence of p53 proteins by working like other molecular glues, using mutant p53 as a “presenter protein” whose surface may form a complementary interaction with essential components of cellular machinery. The prior observation that members of the manumycin polyketide family of natural products are molecular glues between p53 and UBR7^56^ indicate that molecular glues involving p53 are indeed possible.

Future p53-selective therapies may bind to protein folds found on wild-type p53 proteins^57^ and use elevated p53 protein abundance to generate compounds that selectively target the majority of cancers carrying *TP53* missense mutations. Because most sources of cell stress can induce the accumulation of p53, the optimal pan-p53 selective compounds are likely to be covalent and dosed in pulses such that the therapeutic window between cancer and normal tissue is maximized. Other future compounds may bind selectively to mutant p53 proteins whose surface allows for selective small molecule binding. Furthermore, targeting certain essential genes may promote accumulation of mutant p53 protein, as was shown for WEE1 degraders recently^58^, possibly enabling larger therapeutic windows.

If clinical-grade compounds that can specifically kill cells with increased p53 abundance can be generated, they may enjoy roles outside of cancer therapy. In one example, p53 is highly expressed in senescent cells and deficiency of MDM2 causes a progeroid syndrome in humans^59,60^. Future p53-targeting compounds of this class could function as senolytics.

New targeted therapies are inevitably followed by the emergence of on-target and off-target resistance mechanisms. As we have demonstrated, cancer cells evolved resistance by decreasing mutant p53 protein expression upon chronic exposure to intermediate doses of these compounds. These results reflect the observation that antigen loss follows exposure to cytotoxic agents whose activity depends on antigen presence. Nonetheless, we hope that in the fullness of time, gain-of-function p53-selective small molecules will allow for sufficient specific killing of cancer cells such that in combination with other therapies, a greater fraction of patients will achieve meaningful remissions.

## Supporting information

SuppFigures_Note_and_ChemSI

## Acknowledgements

This work was supported by the Svenson Fellowship (W.J.G.), Lubin Scholar Award (WJ.G.), NCI’s Cancer Target Discovery and Development (CTD2) Network (grant number U01CA217848, S.L.S), NCI grant R35CA197568 (M.M.), and an American Cancer Society Research Professorship (M.M.). We thank Jonathan Ostrem, Claire Harmange-Magnani, Sameek Singh, Prashant Singh, and John Knapp for helpful discussions.

## Author Contributions

Chemical Synthesis: A.S., H-J.C., Conceptualization: W.J.G., Data Curation: W.J.G., A.S., N.G., H-J.C., Formal Analysis: W.J.G., A.S., Funding Acquisition: W.J.G, M.M., S.L.S., Investigation: W.J.G., A.S., Methodology: W.J.G., A.S., Resources: M.M., S.L.S., Software: W.J.G., A.S., Supervision and assistance in shaping the research approach: M.M., S.L.S., Validation: W.J.G., A.S., Visualization: W.J.G., A.S., Writing – original draft: W.J.G., A.S., Writing – review & editing: W.J.G., A.S., M.M., S.L.S. All authors approved of the final draft.

## Methods

### Cancer Genomics Analyses

CRISPR dependency scores, RPPA Z-scored protein expression, RNA-seq, *TP53* mutation calls, and compound sensitivity data were downloaded from the DepMap portal (https://depmap.org/portal/)^34^. Human cell surface proteins (N=1492) were previously reported^62^. Protein abundance in 293T cells was downloaded from OpenCell^35^. Immunohistochemistry images were downloaded from the Human Protein Atlas^61^. To identify genes that were both essential and had low protein abundance, we filtered to those with mean Chronos scores ≤ -2 and with protein abundance ≤ 920 nM (p53 protein abundance in 293T cells). We ranked the remaining genes based on (log_10_(protein concentration) + mean Chronos score) and identified those with the lowest score using this metric.

### MFE319 TP53 Mutation Analysis

MFE319 paired RNA-seq reads were downloaded from CCLE (SRR8615235)^31^. The reads were aligned directly to the p53 major isoform mRNA transcript fasta (NM_000546.6) using STAR^63^. The resulting .bam file was visualized in IGV^64^. Read pairs spanning both mutated residues (Y220C, R273C) were used to infer the mutations were *in trans*.

### Modeling of Compound Accumulation

We assume the cell membrane is permeable to the bifunctional small molecule, allowing it to freely diffuse. We also assume p53 is in greater abundance than the other target bound by the bifunctional small molecule, such that the binding of the other target does not contribute meaningfully to intracellular trapping of the molecule. We define the following variables: [M_A0_] is the initial concentration of the bifunctional small molecule outside of the cell (which is a parameter we vary). [M_B0_]=0 is the initial concentration of the bifunctional small molecule inside the cell. [P_0_]=1e-6 M is the initial intracellular concentration of the p53 (defined based on abundance from OpenCell^35^). [MP_0_]=0 is the initial intracellular concentration of the molecule-p53 complex. We also define constants for diffusion (k_diff_ = 1e3 s^-1^), binding of a pharmaceutically optimized bifunctional molecule to p53 (k_bind_ = 8.7e5 M^-1^ s^-1^), dissociation of a pharmaceutically optimized bifunctional molecule from p53 (k_unbind_ = 1e-6 s^-1^), and p53 protein half-life (t_1/2_, which is a parameter we vary).

We define the following rates:

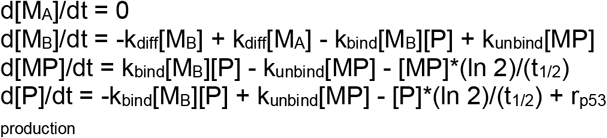

Note: The constant rate of p53 protein production, r_p53 production_, was estimated using the concentration of p53 missense mutant protein at steady state in a cell. d[p53]/dt = protein production rate - protein degradation rate = 0

d[p53_missense_]/dt = protein production rate - [p53_missense_]^*^ln(2) / t_1/2-p53missense_ = 0

Protein production rate =[p53_missense_]^*^ln(2) / t_1/2 -p53missense_, with t_1/2-p53missense_ = 24 hours

We ran an ODE solver in R (package: deSolve) using these parameters, and calculated final intracellular ([M_B_] + [MP]) and extracellular ([M_A_]) bifunctional small molecule concentrations at t=600 hours.

### Cell Culture

293T cells were obtained from ATCC, MFE319 from DSMZ, and Huh7 from the JCRB. All cell lines were cultured in DMEM supplemented with 10% FBS, and 100 IU/mL of penicillin, 100 μg/mL of streptomycin at 37°C in 5% CO_2_.

### Plasmids

Plasmids were ordered as codon-optimized entry vectors from TWIST (pTwist-ENTR). This includes the following constructs: Halo-p53^R273H^(FL)-mCherry, Halo-p53^Y220C^(ΔTAD)-mCherry, Halo-p53^Y220C^(FL)-mCherry, Halo-p53^WT^(FL)-mCherry-2A-mTagBFP2-V5, Halo-p53^R273H^(FL)-mCherry-2A-mTagBFP2-V5, NLS-LgBiT-p53^Y220C^(DBD), mEGFP-PLK1-SmBiT. p53^Y220C^ was ordered with stabilizing mutations (M133L/V203A/N239Y/N268D)^41^. Plasmids were Gateway cloned into lentiviral EF1a expression vector pLEX307 using LR clonase II (Invitrogen). The p53 reporter (#90363)^54^ and Cas9-EGFP plasmid were ordered from Addgene (#82416)^65^. DNA oligos encoding the *TP53* sgRNA (top strand: CACCGCAGAATGCAAGAAGCCCAGA, bottom strand: AAACTCTGGGCTTCTTGCATTCTGC) were ordered from GeneWIZ and restriction cloned into the Cas9-EGFP plasmid. Sequences were verified through Primordium whole-plasmid sequencing.

### Transient Transfection and Stable Cell Line Creation

Plasmids were transfected into 293T cells using TransIT-LT1 transfection reagent following the manufacturer’s protocol. Lentivirus was generated transfecting psPAX2, pMD2.G, and the cloned pLEX307 plasmid (3:3:2 ratio) into 293T cells. Lentivirus was collected 2 days after transfection and stable cell lines were established infecting with filtered lentivirus and polybrene (10 μg/mL). Cells were switched to selection media 2 days after infection.

### Cell Titer Glo

Cells were plated in 384-well plates (typically 1500 cells/well) with varying concentrations of compounds with a total volume of 50 μL/well. Cell titer glo (CTG) reagent was added (25 μL) to the solutions, mixed, and readout using an EnVision 2105 multimode plate reader.

### Crystal Violet

293T cells were plated in 6- or 12-well plates at low density (∼10% confluence) and treated with compound for the indicated duration. Cells were rinsed once with PBS then stained with crystal violet (0.5% m/v) in 20% methanol/water for 10 minutes. Cells were washed with water five times and air-dried overnight.

### Flow Cytometry-Based Competition Assays

Cells with and without a fluorescent marker (e.g. Halo-p53-mCherry) were mixed and the resulting solution was plated in 6-well plates or 10 cm dishes at low density and compound was added at varying concentrations. After reaching confluence, cells were passaged and retreated with compound if needed. Otherwise, cells were trypsinized, washed, and resuspended in complete growth media in 96-well plates or tubes for flow cytometry. Untreated cell mixtures were used to establish gates separating the mCherry+/- populations. Analysis was conducted on a CytoFLEX LX Flow Cytometer.

### NanoBiT Assay

293T cells were seeded in 6-well plates and co-transfected with NLS-LgBiT-p53^Y220C^(DBD) and mEGFP-PLK1-SmBiT (1:1). After one day, the cells were passaged into 96-well plates and treated with varying concentrations of compound. One day after treatment, nanoluciferase substrate was added to the cells and luminescence was monitored using an EnVision 2105 multimode plate reader.

### Colocalization Assay and Microscopy

293T cells were seeded in 6-well plates and co-transfected with Halo-p53^Y220C^ΔTAD-mCherry and mEGFP-PLK1-SmBiT (1:1). After one day, the cells were passaged into 96-well plates in FluoroBrite DMEM for microscopy. The cells were imaged with the Opera Phenix Plus High-Content Screening System (PerkinElmer) before compound treatment and as a time series following compound treatment.

### G2/M Arrest Assay

Halo-p53^Y220C^ΔTAD-mCherry or parental 293T cells were plated in 6-well dishes 24 hours prior to treatment. Cells were then exposed to p53-01 (0.125 μM, 0.250 μM, 0.5 μM) for 16 hours in triplicate, then harvested by trypsinization, washed twice with cold 1x PBS, fixed by dropwise addition of ice-cold 70% ethanol while vortexing, and incubated overnight (4ºC). Fixed samples were centrifuged at 1000 g x 5 minutes and 70% ethanol was removed. Cells were washed twice with 1X PBS + 1% BSA prior to RNA degradation using 1 mg/mL RNase A and DNA labeling with 50 μg/mL propidium iodide for 30 minutes at 37°C. DNA content was measured by flow cytometry (Beckman CytoFLEX LX) and data were analyzed using FlowJo.

### Huh7 RNA-sequencing and Analysis

Huh7 cells were passaged into 6-well plates and treated with PMV6 (4 μM), p53-01 (4 μM), or DMSO for 24 hours. Cells were washed once with cold PBS. Subsequently, TRIzol (Invitrogen) was added to cells, and following the manufacturer’s protocol, RNA was extracted. RNA concentration was monitored using a Qubit Fluorometer (ThermoFisher), and RNA integrity was analyzed using an Agilent Bioanalyzer. The NEBNext Ultra II RNA Library Prep Kit (Illumina) was used to prepare a RNA-seq library. A NovaSeq 6000 machine (Illumina) was used for paired-end 150 bp RNA-sequencing. STAR/RSEM^63,66^ was used to align RNA-seq reads to the GENCODE v38 transcript reference^67^ and generate a count matrix. Raw counts were rounded and DESeq2 was used for downstream analysis^68^. A pre-ranked GSEA^69^ was performed using the DESeq2 t-statistic (vs. DMSO-treated cells) as a rank metric using the Fisher p53 targets geneset^70^.

### P53 Reporter Assay

Huh7 and MFE319 cell lines stably expressing a p53 reporter (Addgene #90363)^54^ were generated. Cells were plated in 384-well plates (1500 cells/well) and treated with varying concentrations of compound. 16 hours after treatment, firefly luciferase substrate was added to the cells and luminescence was monitored using an EnVision 2105 multimode plate reader.

### RT-qPCR

Calu1 cells stably expressing Halo-p53(FL)-mCherry-2A-mTagBFP2-V5 were passaged into 6-well plates. RNA was extracted using a RNeasy Plus Mini Kit (Qiagen) following the manufacturer’s instructions. RNA (500 ng) was reverse transcribed to cDNA using SuperScript VILO Master Mix following the manufacturer’s instructions (total volume: 20 μL). The resulting solution was diluted 1:5 with water and 2 μL of cDNA was used in each qPCR reaction in 384-well PCR plates (AB1384W). 2X Power SYBR Green PCR Master Mix (Fisher) and primers (150 nM final concentration) were added to a total volume of 15 μL. Primers used – Halo-p53(FL)-mCherry-2A-mTagBFP2-V5 forward: CAGGACGGCTGCCTTATTTA, Halo-p53(FL)-mCherry-2A-mTagBFP2-V5 reverse: AGACGGCAGATCGCAATATC, GAPDH forward: GTCTCCTCTGACTTCAACAGCG, GAPDH reverse: ACCACCCTGTTGCTGTAGCCAA. Amplification was performed following the manufacturer’s protocol on a QuantStudio 7 Flex Real-Time PCR System.

