## Supplementary material for "p53 protein abundance is a therapeutic window across TP53 mutant cancers and is targetable with proximity inducing small molecules": SuppFigures_Note_and_ChemSI

**Supporting Information**

Supplementary Figures

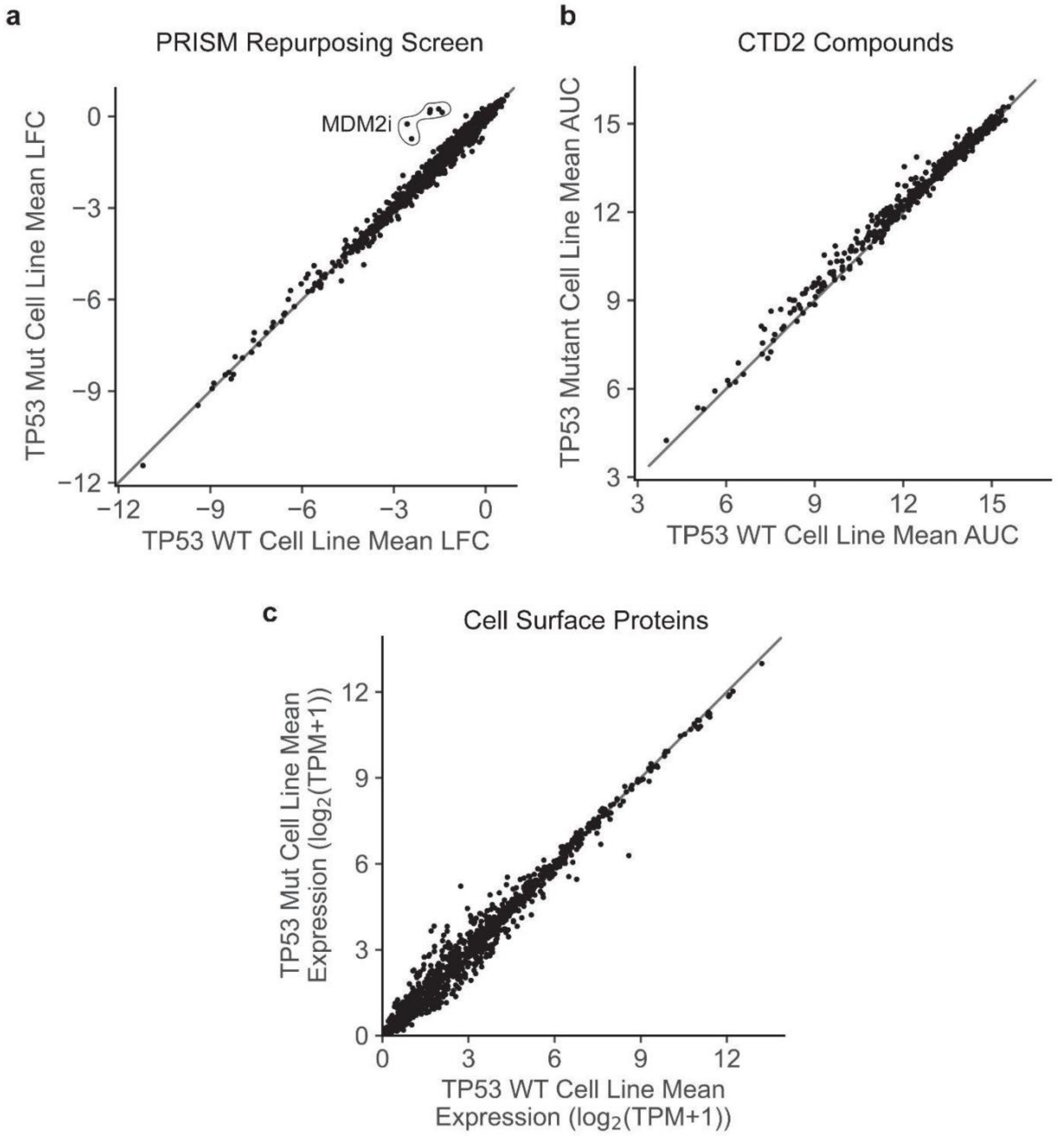


**Fig. S1** | No small molecules specifically kill p53 mutant cancer cells or cell surface receptors upregulated on TP53 mutant cells. **a**, Mean $log_2$(fold-change) for compounds in the PRISM repurposing library (23Q2) on the viability of *TP53* mutant (y-axis) and *TP53* wild-type cell lines (x-axis) across DepMap. MDM2 inhibitors are boxed. **b**, Mean area under the curve (AUC) for compounds in the CTD2 library on the viability of *TP53* mutant (y-axis) and *TP53* wild-type cell lines (x-axis) across DepMap. **c**, Mean mRNA expression ($log_2$(TPM+1)) for genes encoding cell surface proteins for *TP53* mutant (y-axis) and *TP53* wild-type cell lines across DepMap.

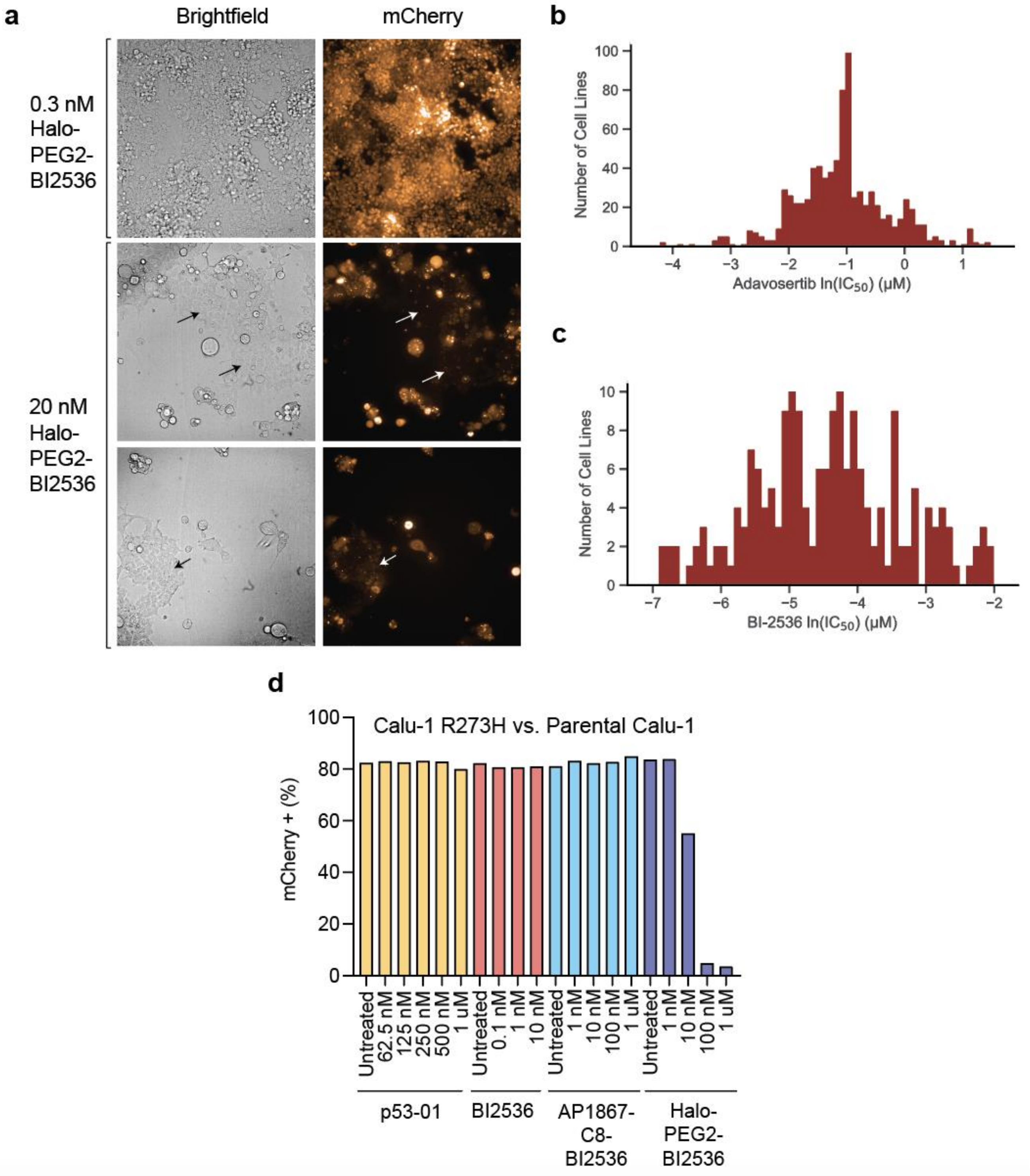


**Fig. S2** | Halo-PEG2-BI2536 selectively kills cells overexpressing p53. **a**, Brightfield and mCherry imaging of Halo-p53$^{R273H}$(FL)-mCherry 293T cells treated with Halo-PEG2-BI2536 (top row: 0.3 nM, bottom two rows: 20 nM) for 8 days. **b**, ln($IC_{50}$) for adavosertib in DepMap (PRISM OncRef), **c**, ln($IC_{50}$) for BI-2536 in DepMap (GDSC1). **d**, Competition of Halo-p53$^{R273H}$(FL)-mCherry vs. parental Calu-1 cells with p53-01, BI2536, AP1867-C8-BI2536, or Halo-PEG2-BI2536 for 7 days; analyzed for mCherry expression by flow cytometry.

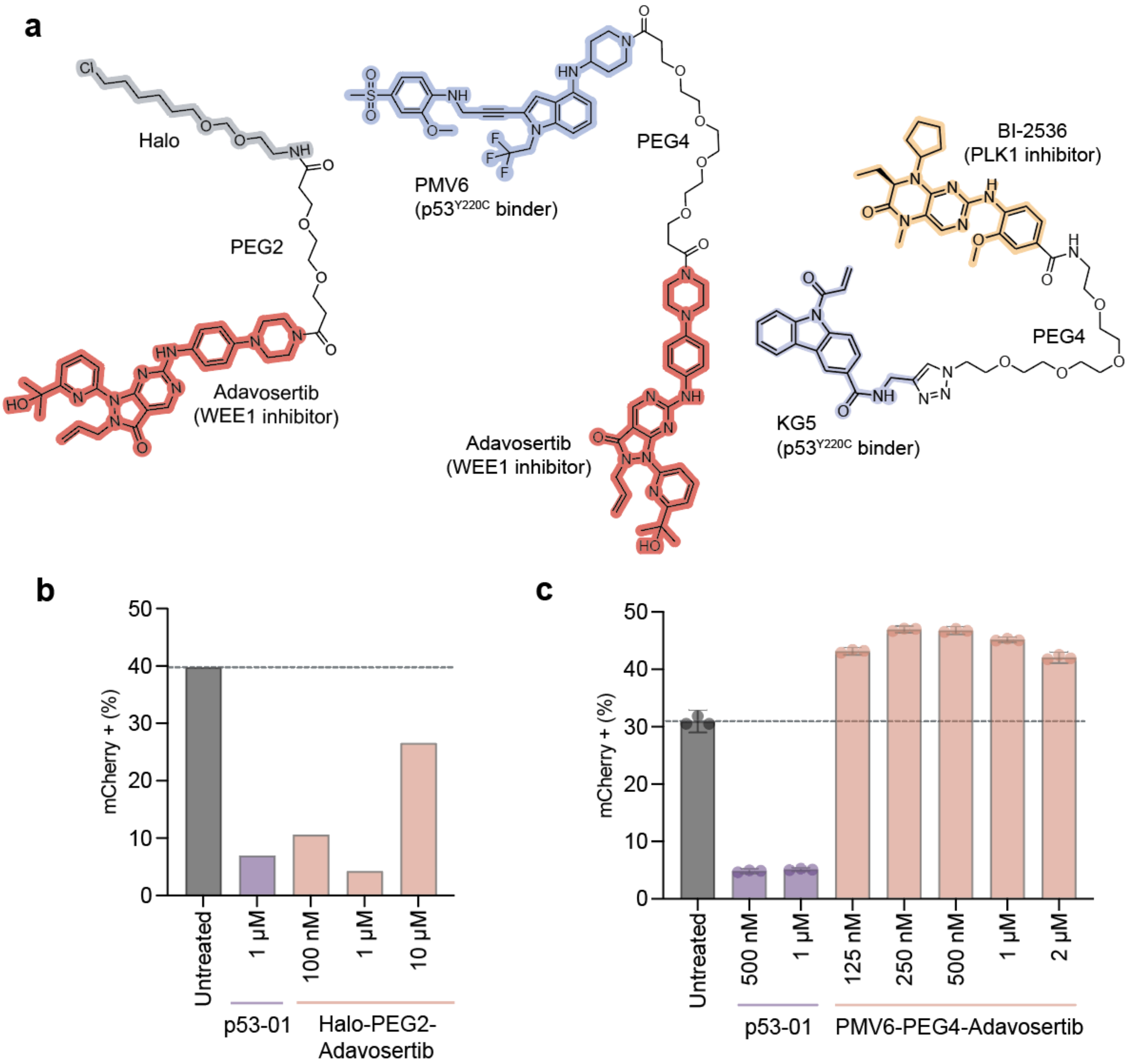


**Fig. S3** | Other classes of bifunctional small molecules have lower activity. **a**, Structure of Halo-PEG2-adavosertib (left), PMV6-PEG4-adavosertib (middle), and KG5-PEG4-BI2536 (right). **b**, Competition of Halo-p53$^{Y220C}$ΔTAD-mCherry vs. parental 293T cells with p53-01 or Halo-PEG2-adavosertib for 8 days; analyzed for mCherry expression by flow cytometry. **c**, Competition of Halo-p53$^{Y220C}$ΔTAD-mCherry vs. parental 293T cells with p53-01 or PMV6-PEG4-adavosertib for 10 days; analyzed for mCherry expression by flow cytometry.

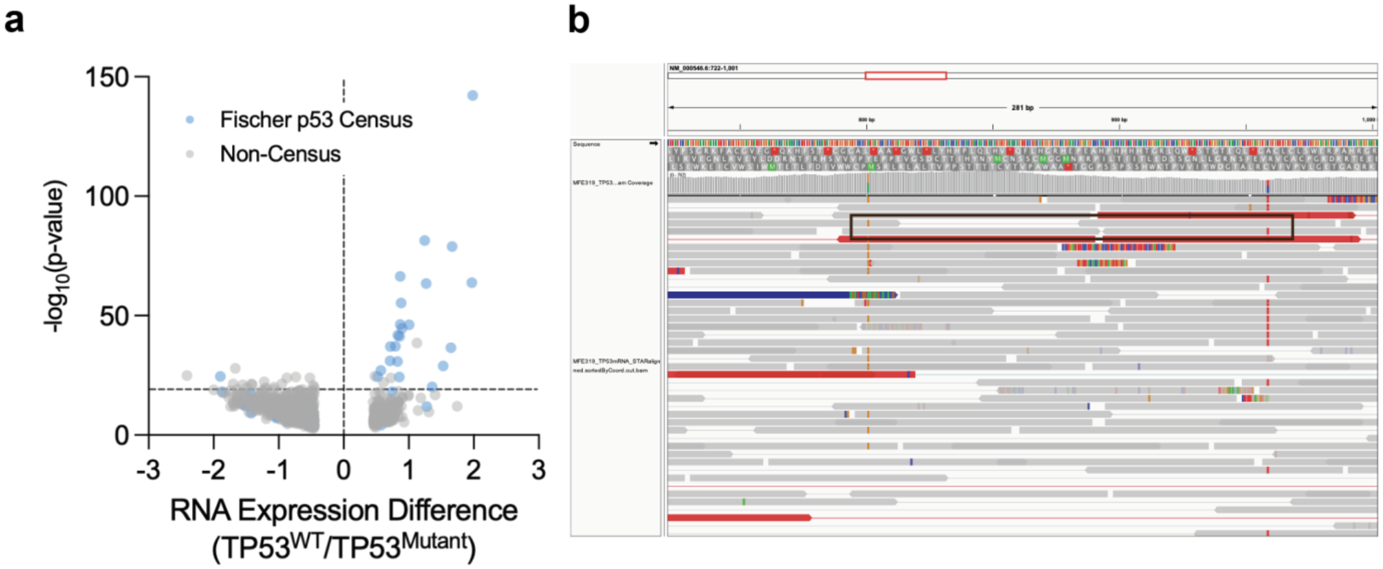


**Fig. S4** | p53-01 is active in endogenous settings. **a**, Difference in mRNA expression between *TP53* WT and mutant cell lines in DepMap for all genes. *P*-values computed by t-test. Fischer census p53 targets are colored in blue. **b**, Analysis of RNA-seq reads aligned to p53 transcript sequence in MFE319 and visualized in IGV. Paired reads boxed in red span residues 220-273 demonstrate the Y220C and R273C mutations are present in *trans*.

Supplementary Note. *TP53 mutant cancers are associated with approximately half of cancer deaths.*

To estimate the fraction of cancer deaths that occur in patients with detectable TP53 mutations, we queried the cBioPortal database[7,8], combining all non-redundant studies. This query retrieved 224 studies, 80,085 samples from 76,363 patients. 32.1% of patients harbored a TP53 mutation (28,102 total mutations detected). Survival data were only available for 33,176 patients, 10,350 of whom had a TP53 alteration (31.2%).

Disease specific survival data were only available for 10,021 patients, 3,709 of whom had a TP53 alteration (37.0%). The 10-year disease-specific survival rate was 61.10% in the TP53 WT group and 44.60% in the TP53 mutation group (HR: 1.62; CI-95%: 1.48-1.763; $p<10^{-10}$). The number of disease-specific deaths that occurred in the TP53 mutant group was 1,016 and the number of deaths in the WT group was 1,212. The fraction of cancer-specific deaths that occurred in TP53 mutant patients was 45.6%.

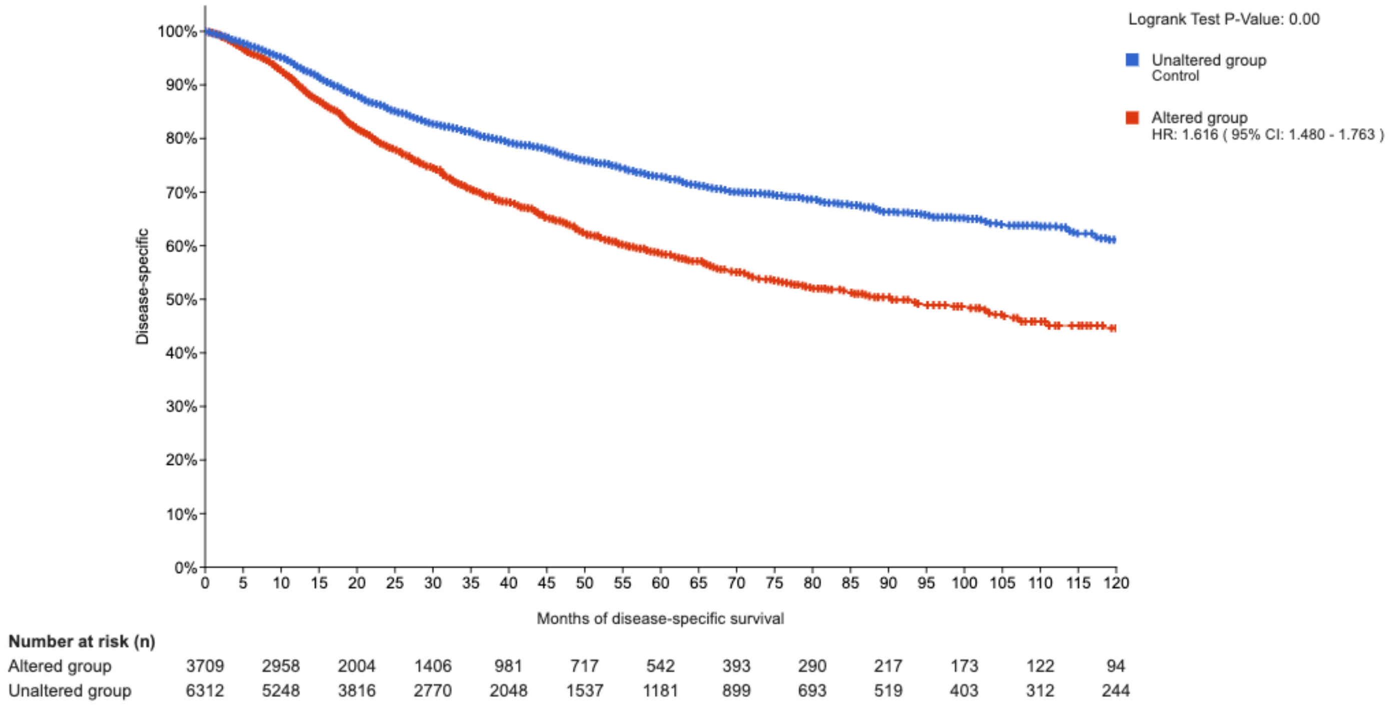

Chemistry Supplementary Information

General Procedures. Commercial reagents, without additional purification, were used for all syntheses. Round-bottom flasks were used for reactions with Teflon-coated magnetic stir bars. Reaction progress was monitored using ultra-performance liquid chromatography mass spectrometry on a Waters ACQUITY UPLC I-Class 15 PLUS System with an ACQUITY SQ Detector 2. Nuclear magnetic resonance (NMR) spectra were generated at the Broad Institute of MIT and Harvard, using a Bruker AVANCE III HD 400 MHz spectrometer at room temperature ($^{1}$H NMR, 400 MHz; $^{13}$C NMR 101 MHz). Chemical shifts for 1H NMR and 13C NMR are given in parts per million (ppm), referenced to residual solvent signals. DMSO-d6 and $CD_3CN$ were acquired from Cambridge Isotope Laboratories, Inc. and Oakwood Products, Inc., respectively. $^{1}$H NMR data is reported as follows: chemical shift value in ppm, multiplicity (s = singlet, d = doublet, t = triplet, dd = doublet of doublets, and m = multiplet), coupling constant value in Hz, and integration value. Electrospray ionization-high-resolution mass spectrometry (ESI-HRMS) was performed at the Harvard Center for Mass Spectrometry.

Organic Synthesis:

*PMV6.*

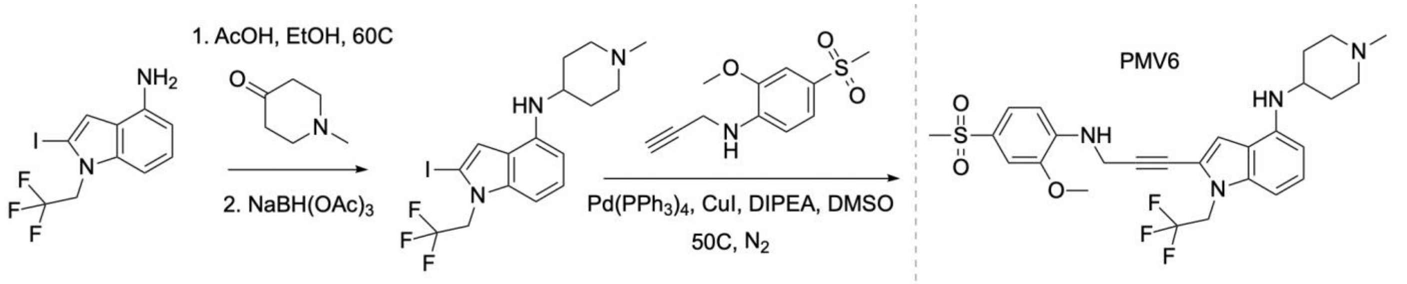


**2-(3-((2-methoxy-4-(methylsulfonyl)phenyl)amino)prop-1-yn-1-yl)-N-(1-methylpiperidin-4-yl)-1-(2,2,2-trifluoroethyl)-1H-indol-4-amine (PMV6, PMV6-NMe).** To a solution of 2-iodo-1-(2,2,2-trifluoroethyl)-1H-indol-4-amine (50 mg, 0.147 mmol, eNovation Chemicals) in ethanol (5 mL), 1-methyl-4-piperidone (100 µL, 0.867 mmol) and acetic acid (50 µL, 0.874 mmol) were added, and the mixture was heated to 60 °C with strong stirring. After 30 minutes, sodium triacetoxyborohydride (300 mg, 1.415 mmol) was added and allowed to react overnight. After completion, the reaction was purified by reverse-phase HPLC (acetonitrile:water gradient up to 90%) to yield 2-iodo-N-(1-methylpiperidin-4-yl)-1-(2,2,2-trifluoroethyl)-1H-indol-4-amine as an off-white solid (26 mg, 0.0595 mmol, 40.5% yield). Subsequently, 2-iodo-N-(1-methylpiperidin-4-yl)-1-(2,2,2-trifluoroethyl)-1H-indol-4-amine (10 mg, 0.0229 mmol), 2-methoxy-4-(methylsulfonyl)-N-(prop-2-yn-1-yl)aniline (10 mg, 0.0418 mmol, 1ClickChemistry), $Pd(PPh_3)_4$ (4 mg, 0.00346 mmol), and CuI (2 mg, 0.0105 mmol) were combined and dissolved in DMSO. The mixture was heated to 50 °C with strong stirring, as nitrogen was bubbled through the mixture. DIPEA (20 µL, 0.115 mmol) was added to the mixture and it was allowed to react overnight under a nitrogen atmosphere. After completion, the reaction was purified by reverse-phase HPLC (acetonitrile:water gradient up to 90%) to yield PMV6 as a light brown solid (3.1 mg, 0.00566 mmol, 24.7% yield). $^{1}$H NMR (400 MHz, DMSO-d6) δ 7.39 (dd, $J$ = 8.4, 2.0 Hz, 1H), 7.26 (d, $J$ = 2.1 Hz, 1H), 7.09 (s, 1H), 7.00 (t, $J$ = 8.0 Hz, 1H), 6.89 (d, $J$ = 8.4 Hz, 1H), 6.68 (d, $J$ = 8.2 Hz, 1H), 6.49 (t, $J$ = 6.2 Hz, 1H), 6.16 (d, $J$ = 7.8 Hz, 1H), 5.49 (d, $J$ = 7.9 Hz, 1H), 4.93 (q, $J$ = 9.0 Hz, 2H), 4.36 (d, $J$ = 6.2 Hz, 2H), 3.90 (s, 3H), 3.10 (s, 3H), 2.79 (d, $J$ = 11.2 Hz, 2H), 2.19 (s, 3H), 2.08-1.98 (m, 2H), 1.92 (d, $J$ = 12.6 Hz, 2H), 1.55-1.42 (m, 2H), 1.28-1.11 (m, 1H). HRMS (top: observed, bottom: theoretical isotope pattern; M+H, $C_{27}H_{31}F_3N_4O_3S$+H] m/z theoretical 549.2142, found 549.2138.

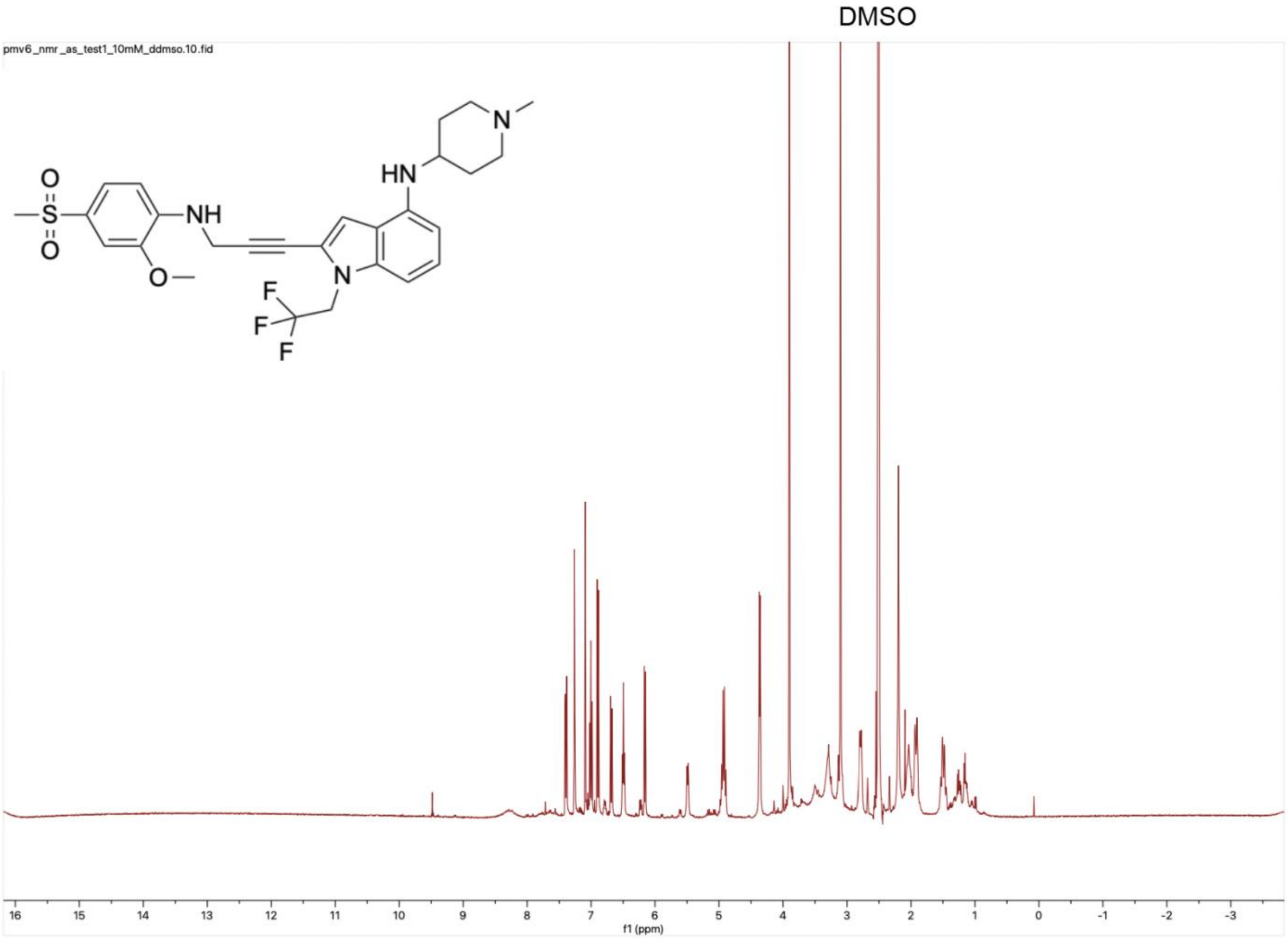
DMSO
pmv6_nmr_as_test1_10mM_ddmso.10.fid
O
S
O
NH
O
HN
N
N
F
F
F
16
15
14
13
12
11
10
9
8
7
6
5
4
3
2
1
0
-1
-2
-3
f1 (ppm)

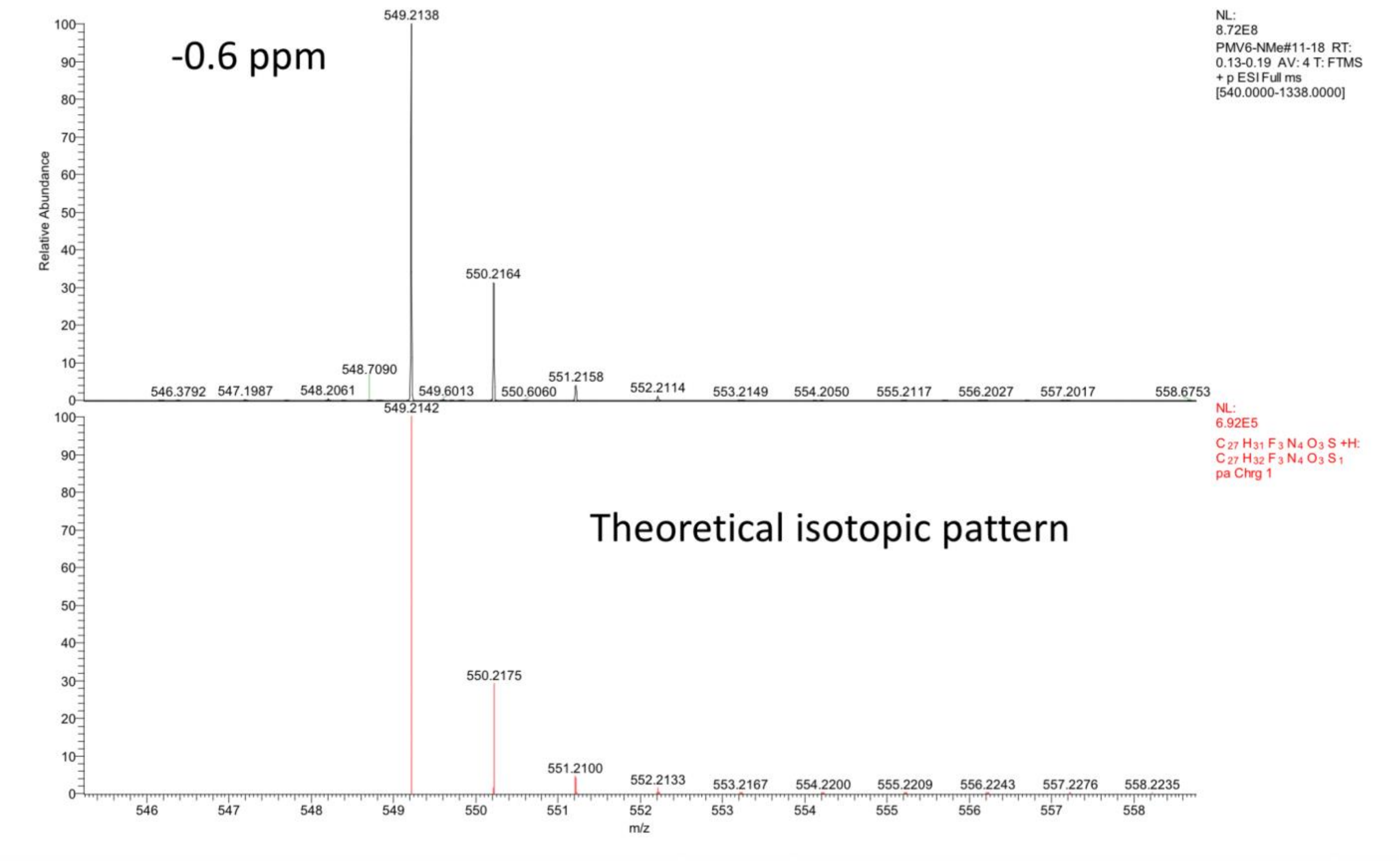


*PMV6-NBoc.*

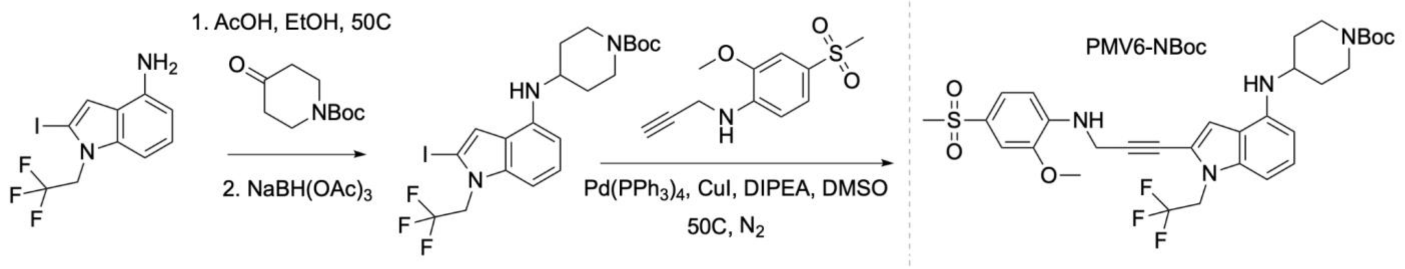


**tert-butyl-4-((2-(3-((2-methoxy-4-(methylsulfonyl)phenyl)amino)prop-1-yn-1-yl)-1-(2,2,2-trifluoroethyl)-1H-indol-4-yl)amino)piperidine-1-carboxylate (PMV6-NBoc).** To a solution of 2-iodo-1-(2,2,2-trifluoroethyl)-1H-indol-4-amine (45 mg, 0.133 mmol, eNovation Chemicals) in ethanol (5 mL), 1-Boc-4-piperidone (95 µL, 0.537 mmol) and acetic acid (50 µL, 0.874 mmol) were added, and the mixture was heated to 50 °C with strong stirring. After 30 minutes, sodium triacetoxyborohydride (108 mg, 0.510 mmol) was added and allowed to react overnight. After completion, the reaction was purified by reverse-phase HPLC (acetonitrile:water gradient up to 90%) to yield tert-butyl-4-((2-iodo-1-(2,2,2-trifluoroethyl)-1H-indol-4-yl)amino)piperidine-1-carboxylate as an off-white solid (18 mg, 0.0344 mmol, 25.9% yield). All of this product, 2-methoxy-4-(methylsulfonyl)-N-(prop-2-yn-1-yl)aniline (16.5 mg, 0.0690 mmol, 1ClickChemistry), $Pd(PPh_3)_4$ (4 mg, 0.00346 mmol), and CuI (2 mg, 0.0105 mmol) were combined and dissolved in DMSO. The mixture was heated to 50 °C with strong stirring, as nitrogen was bubbled through the mixture. DIPEA (36 µL, 0.207 mmol) was added to the mixture and it was allowed to react overnight under a nitrogen atmosphere. After completion, the reaction was purified by reverse-phase HPLC (acetonitrile:water gradient up to 90%) to yield PMV6-NBoc as a brown solid (9.8 mg, 0.0155 mmol, 45.0% yield). $^1$H NMR (400 MHz, DMSO-d6) δ 7.39 (dd, *J* = 8.4, 2.0 Hz, 1H), 7.26 (d, *J* = 2.0 Hz, 1H), 7.06 (s, 1H), 7.01 (t, *J* = 8.0 Hz, 1H), 6.89 (d, *J* = 8.4 Hz, 1H),

6.70 (d, *J* = 8.3 Hz, 1H), 6.49 (t, *J* = 6.2 Hz, 1H), 6.22 (d, *J* = 7.8 Hz, 1H), 5.52 (d, *J* = 7.9 Hz, 1H), 4.93 (q, *J* = 9.1 Hz, 2H), 4.36 (d, *J* = 6.2 Hz, 2H), 3.90 (s, 3H), 3.10 (s, 3H), 2.96-2.80 (m, 2H), 1.93 (d, *J* = 11.6 Hz, 2H), 1.41 (s, 9H), 1.39-1.22 (m, 4H), 0.91-0.79 (m, 1H). HRMS (top: observed, bottom: theoretical isotope pattern; M+H, $C_{31}H_{37}F_3N_4O_5S$+H] m/z theoretical 635.2510, found 635.2499.

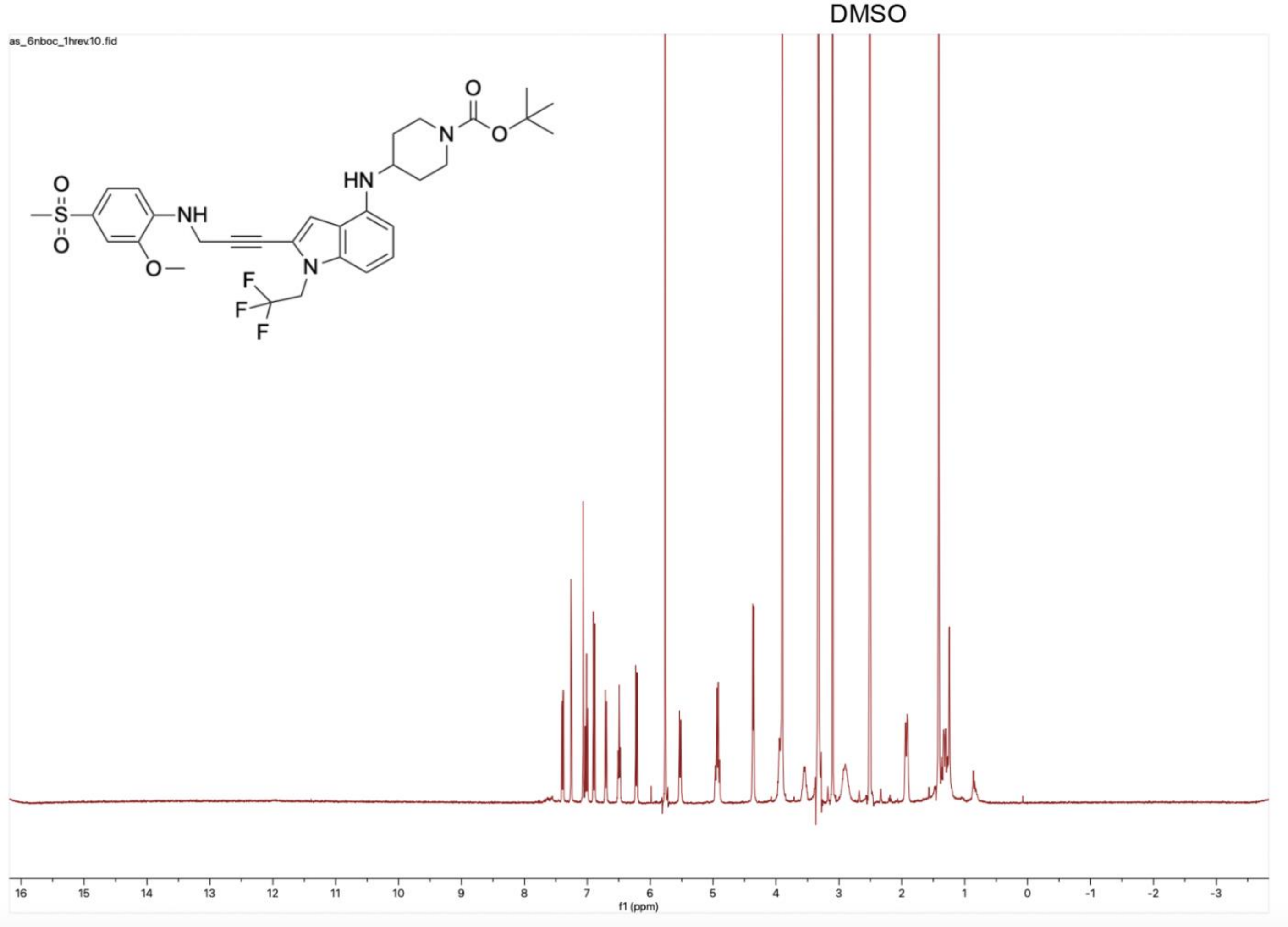

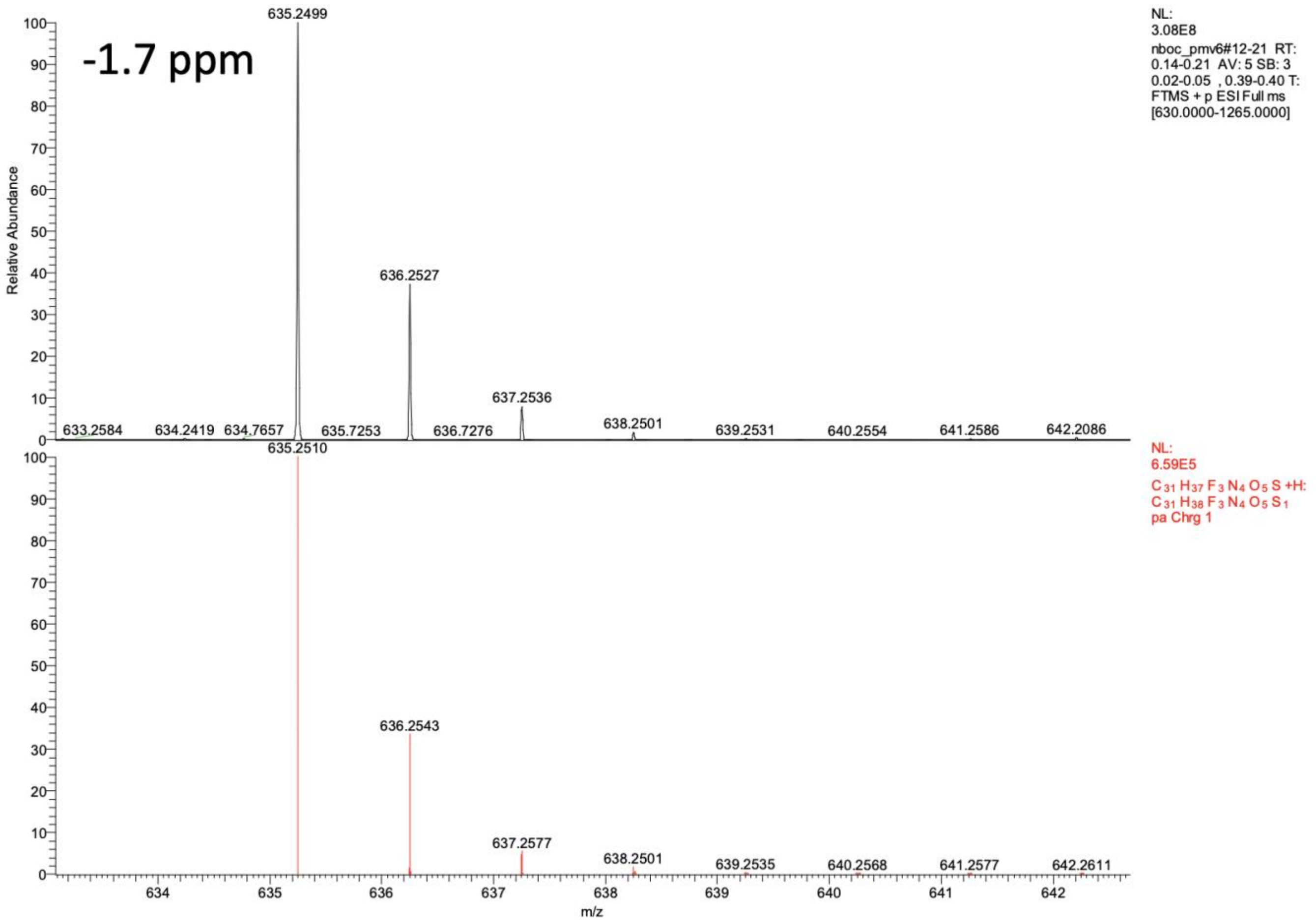


*PMV6-PEG4-BI2536 (p53-01).*

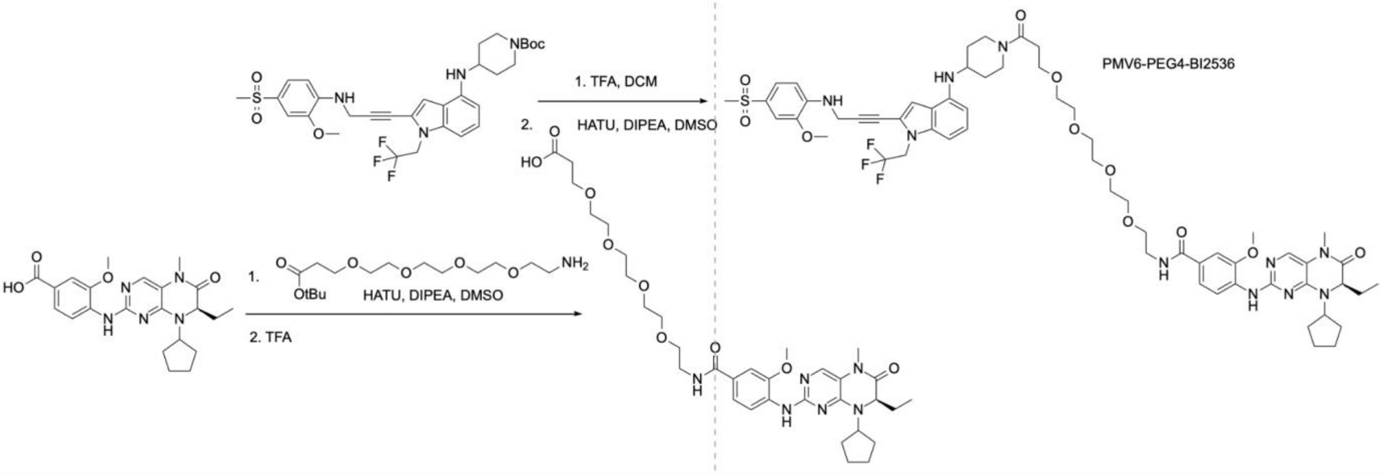


**(R)-4-((8-cyclopentyl-7-ethyl-5-methyl-6-oxo-5,6,7,8-tetrahydropteridin-2-yl)amino)-3-methoxy-N-(15-(4-((2-(3-((2-methoxy-4-(methylsulfonyl)phenyl)amino)prop-1-yn-1-yl)-1-(2,2,2-trifluoroethyl)-1H-indol-4-yl)amino)piperidin-1-yl)-15-oxo-3,6,9,12-tetraoxapentadecyl)benzamide (PMV6-PEG4-BI2536, PMV6-PEG4-BI, p53-01).** To a solution of (R)-4-(8-cyclopentyl-7-ethyl-5-methyl-6-oxo-5,6,7,8-tetrahydropteridin-2-ylamino)-3-methoxybenzoic acid (10 mg, 0.0235 mmol, 1ClickChemistry) in DMSO (1 mL), $H_2N$-PEG4-$CH_2CH_2CO_2$tBu (20 μL, 0.0755 mmol), HATU (12 mg, 0.0315 mmol), and DIPEA (20 μL, 0.115 mmol) were added. After stirring for 15 minutes, trifluoroacetic acid (2 mL) was added and the reaction was stirred for 1 hour. After completion, the reaction was purified by reverse-phase HPLC (acetonitrile:water gradient up to 90%) to yield (R)-1-(4-((8-cyclopentyl-7-ethyl-5-methyl-6-oxo-5,6,7,8-tetrahydropteridin-2-yl)amino)-3-methoxyphenyl)-1-oxo-5,8,11,14-tetraoxa-2-azaheptadecan-17-oic acid (BI2536-PEG4-acid) as a white solid (10 mg, 0.0149 mmol, 63.1% yield). tert-butyl-4-((2-(3-((2-methoxy-4-(methylsulfonyl)phenyl)amino)prop-1-yn-1-yl)-1-(2,2,2-trifluoroethyl)-1H-indol-4-yl)amino)piperidine-1-carboxylate (PMV6-NBoc, 8.2 mg, 0.0129 mmol) was dissolved in DCM (1 mL), and trifluoroacetic acid (2 mL) was added to the solution. After stirring for 1 hour, solvents were removed under reduced pressure. The residue was redissolved in DMSO (1 mL) and BI2536-PEG4-acid (10 mg, 0.0149 mmol), HATU (6 mg, 0.0158 mmol), and DIPEA (20 µL, 0.115 mmol) were added to the solution. After completion (30 minutes), the reaction was purified by reverse-phase HPLC (acetonitrile:water gradient up to 90%) to yield PMV6-PEG4-BI2536 as a brown solid (5.6 mg, 0.00471 mmol, 36.5% yield). $^{1}H$ NMR (400 MHz, DMSO-d6) δ 8.47 – 8.37 (m, 2H), 7.85 (s, 1H), 7.61 (s, 1H), 7.50 (d, *J* = 7.3 Hz, 2H), 7.39 (dd, *J* = 8.3, 2.0 Hz, 1H), 7.26 (d, *J* = 2.0 Hz, 1H), 7.07 (s, 1H), 7.01 (t, *J* = 8.0 Hz, 1H), 6.89 (d, *J* = 8.4 Hz, 1H), 6.71 (d, *J* = 8.2 Hz, 1H), 6.49 (t, *J* = 6.3 Hz, 1H), 6.22 (d, *J* = 7.9 Hz, 1H), 5.53 (d, *J* = 8.0 Hz, 1H), 4.93 (q, *J* = 9.1 Hz, 2H), 4.39 – 4.27 (m, 4H), 4.24 (dd, *J* = 7.6, 3.6 Hz, 1H), 3.91 (d, *J* = 14.4 Hz, 6H), 3.87 (s, 1H), 3.61 (t, *J* = 6.7 Hz, 2H), 3.56 – 3.50 (m, 6H), 3.49 (d, *J* = 3.1 Hz, 7H), 3.44 (dd, *J* = 12.6, 7.0 Hz, 2H), 3.25 (s, 3H), 3.18 (s, 3H), 3.14 (s, 1H), 3.10 (s, 3H), 2.76 (t, *J* = 12.0 Hz, 1H), 2.57 (t, *J* = 6.7 Hz, 2H), 2.07 – 1.84 (m, 5H), 1.83 – 1.71 (m, 3H), 1.70 – 1.55 (m, 3H), 1.42 – 1.22 (m, 2H), 0.76 (t, *J* = 7.4 Hz, 3H). $^{13}C$ NMR (101 MHz, DMSO-d6) δ 168.96, 166.29, 163.40, 154.79, 151.96, 147.15, 146.58, 141.83, 140.99, 138.80, 138.30, 132.68, 129.04 – 124.35 (m), 123.28 – 114.61 (m), 109.63, 109.10, 108.23, 107.61, 100.02, 99.10, 93.63, 73.76, 71.79 – 68.74 (m), 69.59, 67.35, 60.28, 58.85, 56.36 (d, *J* = 20.4 Hz), 49.23 (d, *J* = 32.7 Hz), 44.63 (d, *J* = 33.4 Hz), 33.29, 32.94, 32.68, 29.19, 28.87, 28.25, 26.95, 23.64, 23.39, 9.32. HRMS (top: observed, bottom: theoretical isotope pattern; M+H, $C_{59}H_{75}F_{3}N_{10}O_{11}S$+H] m/z theoretical 1189.5362, found 1189.5350.

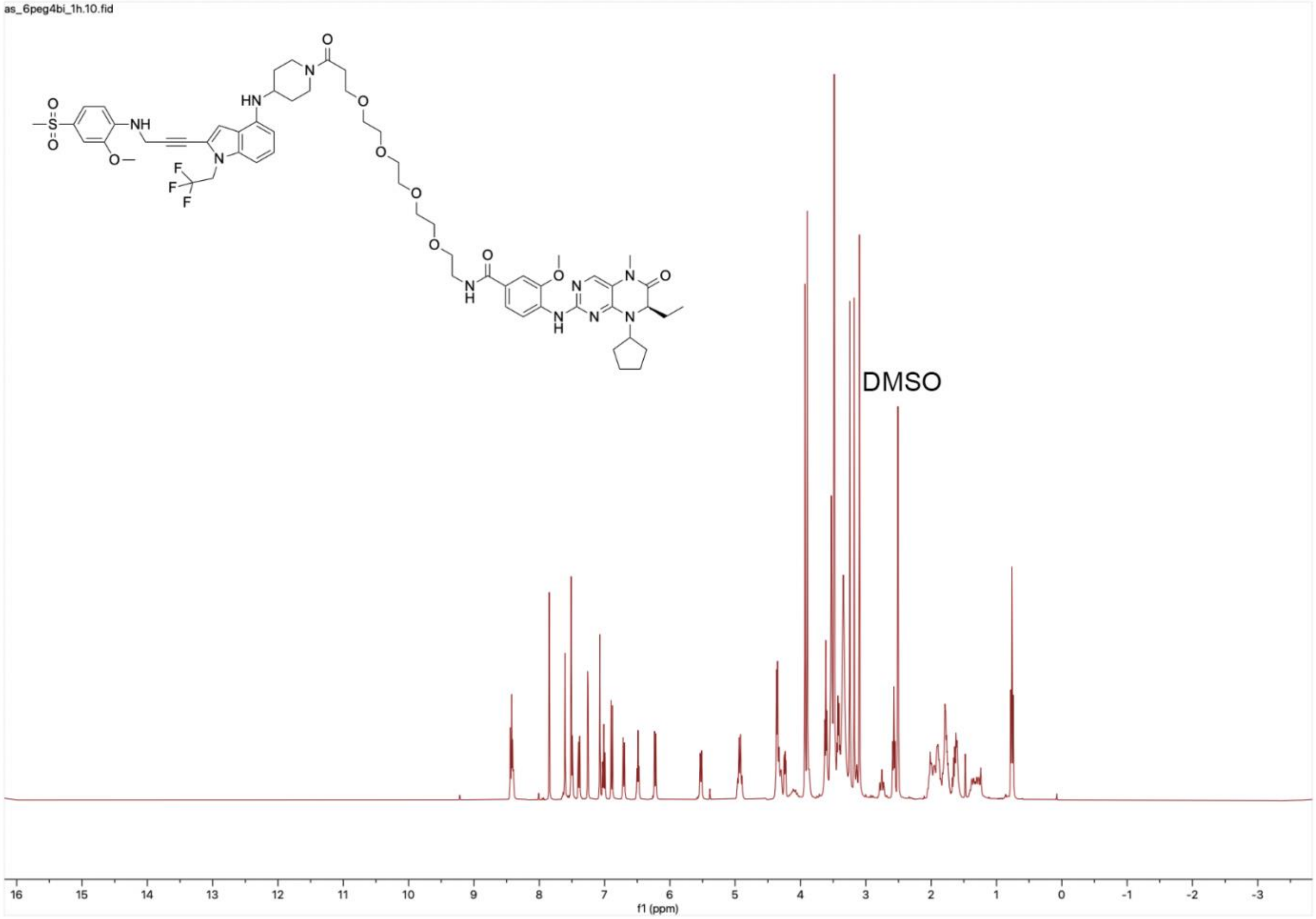
as_6peg4bi_1h.10.fid
DMSO
f1 (ppm)

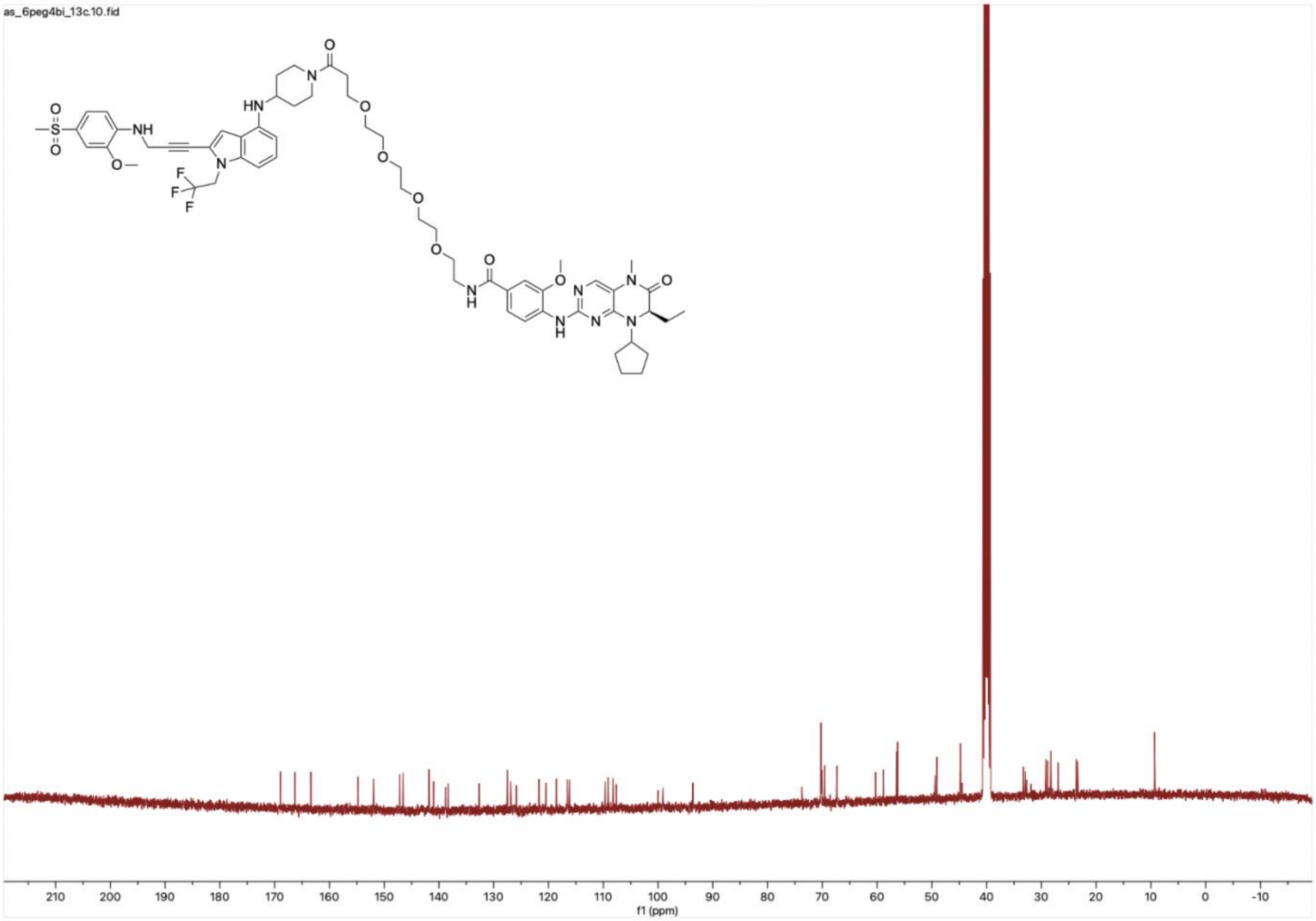
as_6peg4bi_13c.10.fid
210
200
190
180
170
160
150
140
130
120
110
100
90
80
70
60
50
40
30
20
10
0
-10
f1 (ppm)

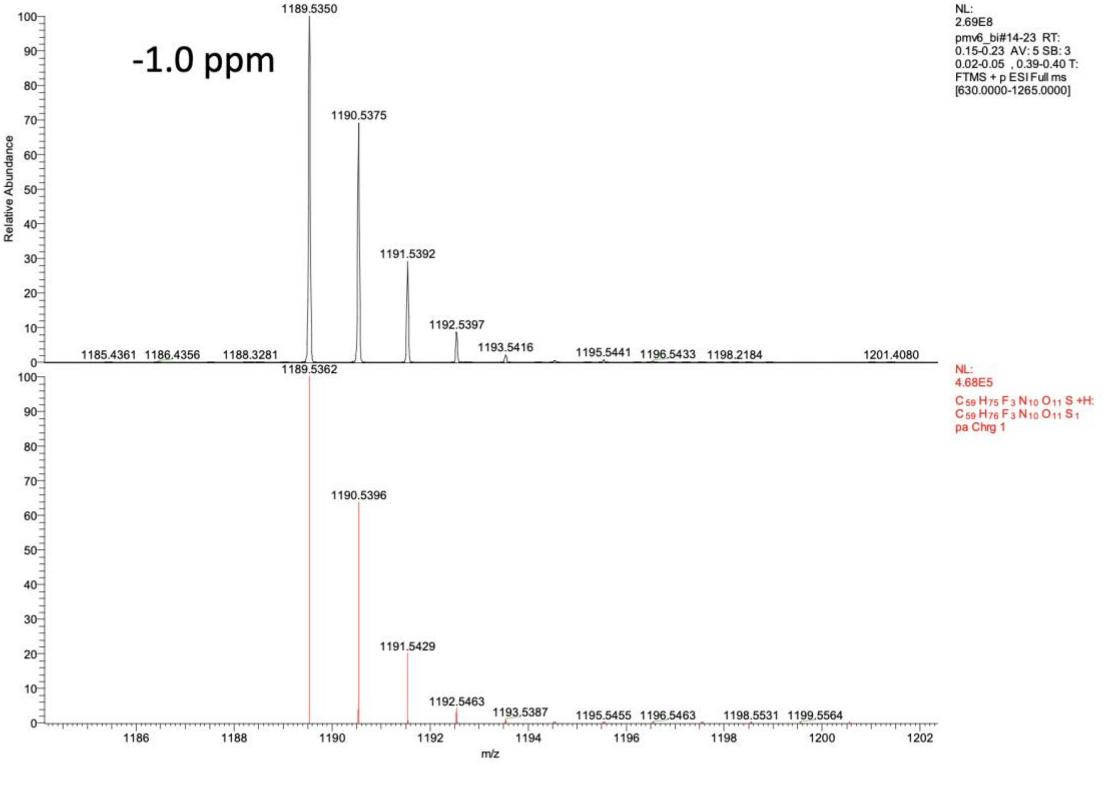
-1.0 ppm
1189.5350
1190.5375
1191.5392
1192.5397
1193.5416
1185.4361
1186.4356
1188.3281
1195.5441
1196.5433
1198.2184
1201.4080
NL:
2.69E8
pmv6_bi#14-23 RT:
0.15-0.23 AV: 5 SB: 3
0.02-0.05 , 0.39-0.40 T:
FTMS + p ESI Full ms
[630.0000-1265.0000]
Relative Abundance
1189.5362
1190.5396
1191.5429
1192.5463
1193.5387
1195.5455
1196.5463
1198.5531
1199.5564
NL:
4.68E5
C59 H75 F3 N10 O11 S +H:
C59 H76 F3 N10 O11 S1
pa Chrg 1
m/z

*Other PMV6-BI2536 bifunctionals.*

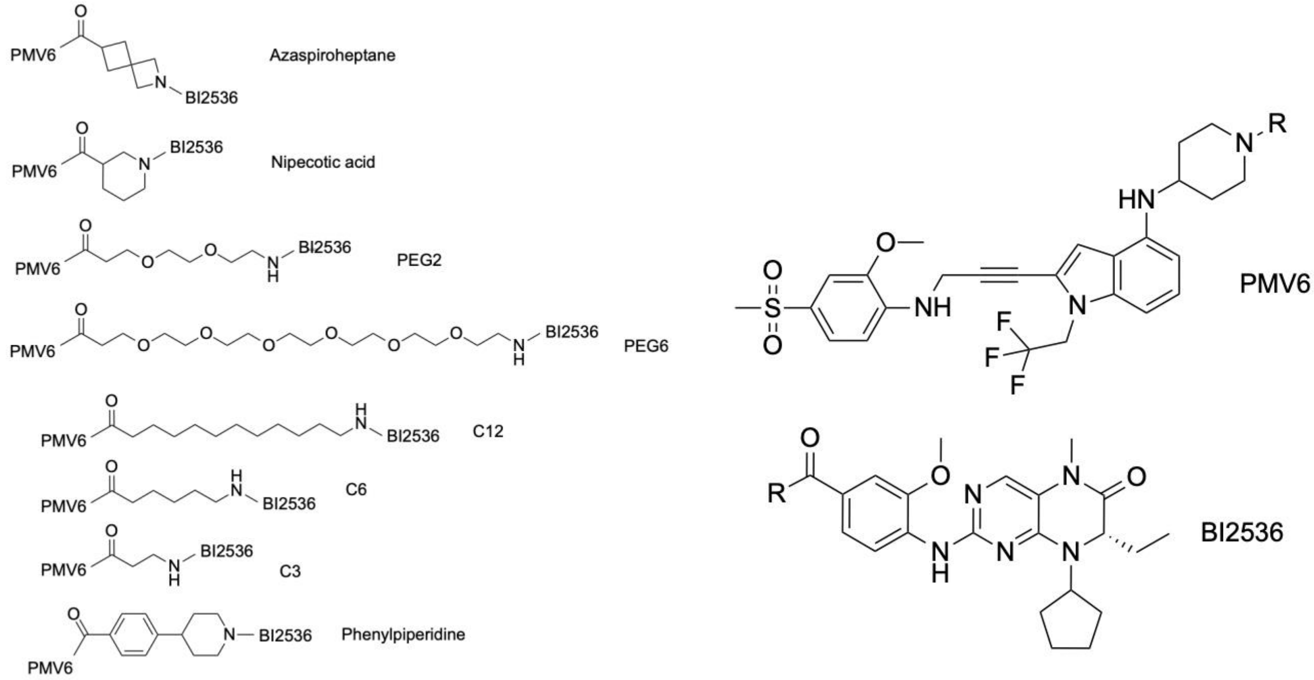


The general procedure for the synthesis of PEG2, C6, and phenylpiperidine compounds was based on the procedure for the synthesis of PMV6-PEG4-BI2536. In brief, the linker (PEG2: amino-PEG2-t-butyl ester, C6: t-butyl-6-aminohexanoate, phenylpiperidine: methyl 4-(piperidin-4-yl)benzoate) was coupled with a stoichiometric amount of (R)-4-(8-cyclopentyl-7-ethyl-5-methyl-6-oxo-5,6,7,8-tetrahydropteridin-2-ylamino)-3-methoxybenzoic acid in the presence of excess HATU (~2 equivalents) and triethylamine (~5 equivalents) in DMSO (1 mL). Subsequently, TFA (3 mL; for PEG2 and C6 linker) or 4M LiOH/$H_2O$ solution (3 mL, phenylpiperidine linker) were added and the solutions were stirred for 4 hours at room temperature. The reactions were purified by reverse-phase HPLC (acetonitrile:water gradient up to 90%) and the resulting compounds were coupled with stoichiometric hydrolyzed PMV6-NBoc (75% TFA in DCM for 1 hour, solvents evaporated), in the presence of excess HATU (~2 equivalents) and triethylamine (~20 equivalents). The reactions were purified by reverse-phase HPLC (acetonitrile:water gradient up to 90%) to yield the products as yellow solids. For the synthesis of azaspiroheptane compound 2-(tert-Butoxycarbonyl)-2-azaspiro[3.3]heptane-6-carboxylic acid was used as the linker. The first coupling occurred with hydrolyzed PMV6-NBoc and the second coupling occurred with (R)-4-(8-cyclopentyl-7-ethyl-5-methyl-6-oxo-5,6,7,8-tetrahydropteridin-2-ylamino)-3-methoxybenzoic acid to yield the product as a yellow solid. For the synthesis of the remaining compounds (PEG6: amino-PEG6-acid, C12: 12-aminododecanoic acid, C3: beta-alanine, nipecotic acid: nipecotic acid), the reaction occurred in one-pot; with stoichiometric hydrolyzed PMV6-NBoc (1 equivalent), linker (1 equivalent), (R)-4-(8-cyclopentyl-7-ethyl-5-methyl-6-oxo-5,6,7,8-tetrahydropteridin-2-ylamino)-3-methoxybenzoic acid (1 equivalent), HATU (2 equivalents), and excess triethylamine (~20 equivalents). Purification by reverse-phase HPLC (acetonitrile:water gradient up to 90%) yielded the products as yellow solids. All reactions were performed at milligram scale, with 3 mg of PMV6-NBoc used per coupling. Overall yields across all steps were as follows: C12: 0.5 mg (9%), PEG2: 0.7 mg (13%), C6: 0.7 mg (14%), Spiro: 0.7 mg (14%), C3: 0.8 mg (17%), PEG6: 1.3 mg (22%), PhPip: 2.0 mg (38%), Nip: 2.4 mg (48%). Purified compounds were dissolved in $CD_3CN$ and $^1H$ NMR was performed (400 MHz, $CD_3CN$). After NMRs were taken, solvent was evaporated under reduced pressure and the residue was redissolved in DMSO to create stock solutions. $^1H$ NMR (400 MHz, $CD_3CN$) and HRMS analysis (top: observed, bottom: theoretical isotope pattern) is provided for all compounds.

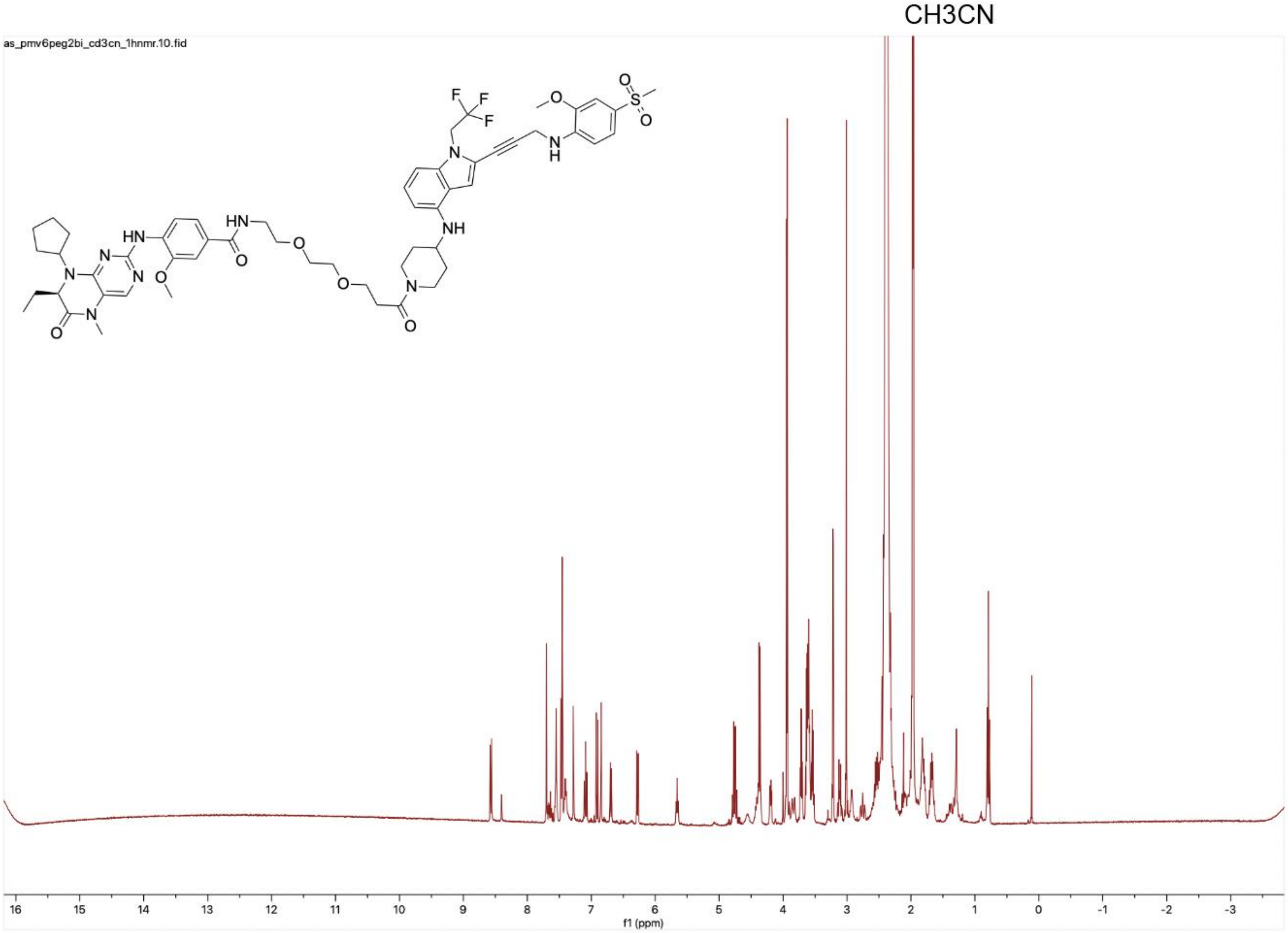

CH3CN
as_pmv6peg2bi_cd3cn_1hnmr.10.fid
f1 (ppm)

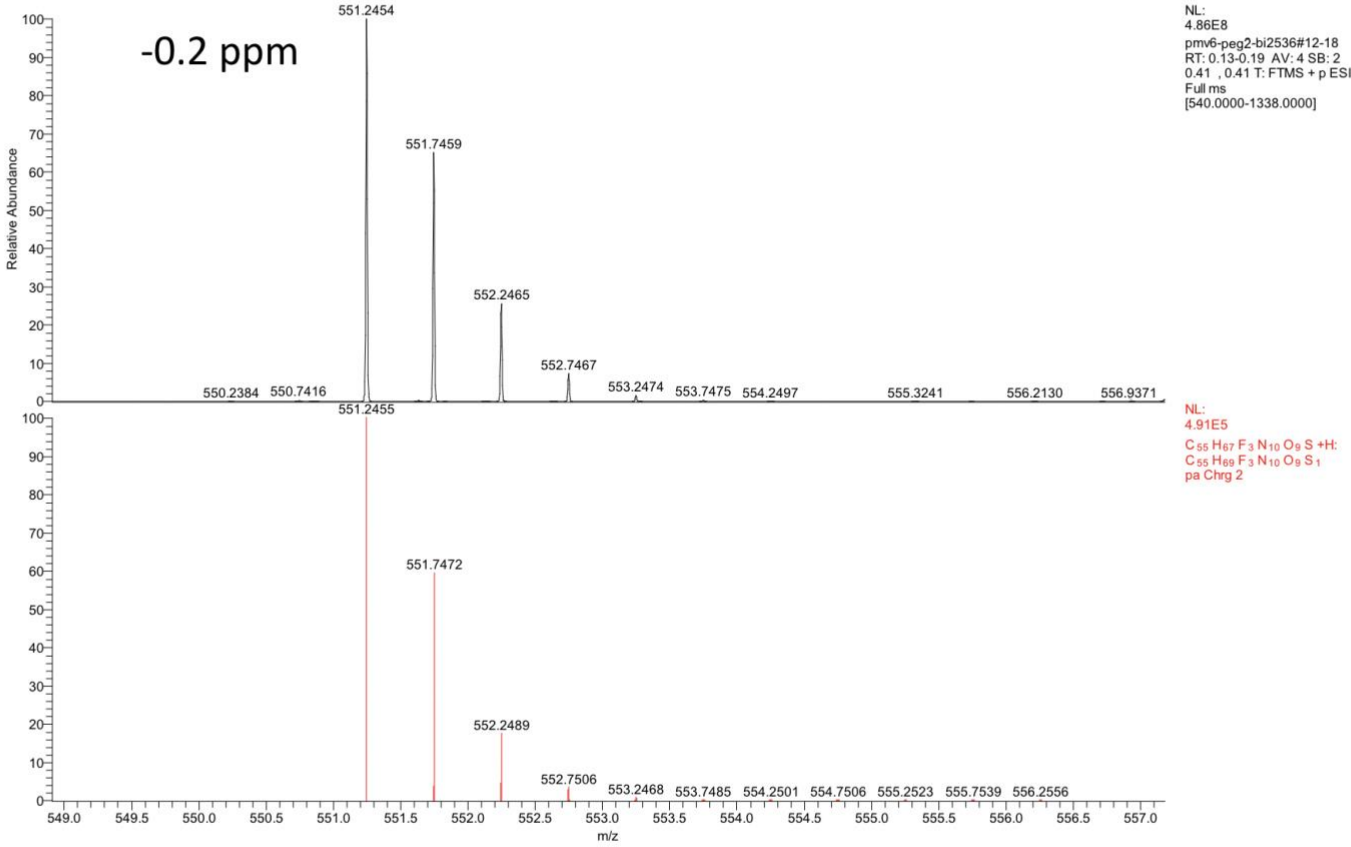

-0.2 ppm
551.2454
551.7459
552.2465
552.7467
553.2474
553.7475
554.2497
550.2384
550.7416
555.3241
556.2130
556.9371
NL:
4.86E8
pmv6-peg2-bi2536#12-18
RT: 0.13-0.19 AV: 4 SB: 2
0.41 , 0.41 T: FTMS + p ESI
Full ms
[540.0000-1338.0000]
551.2455
551.7472
552.2489
552.7506
553.2468
553.7485
554.2501
554.7506
555.2523
555.7539
556.2556
NL:
4.91E5
C 55 H 67 F 3 N 10 O 9 S +H:
C 55 H 69 F 3 N 10 O 9 S 1
pa Chrg 2
Relative Abundance
m/z

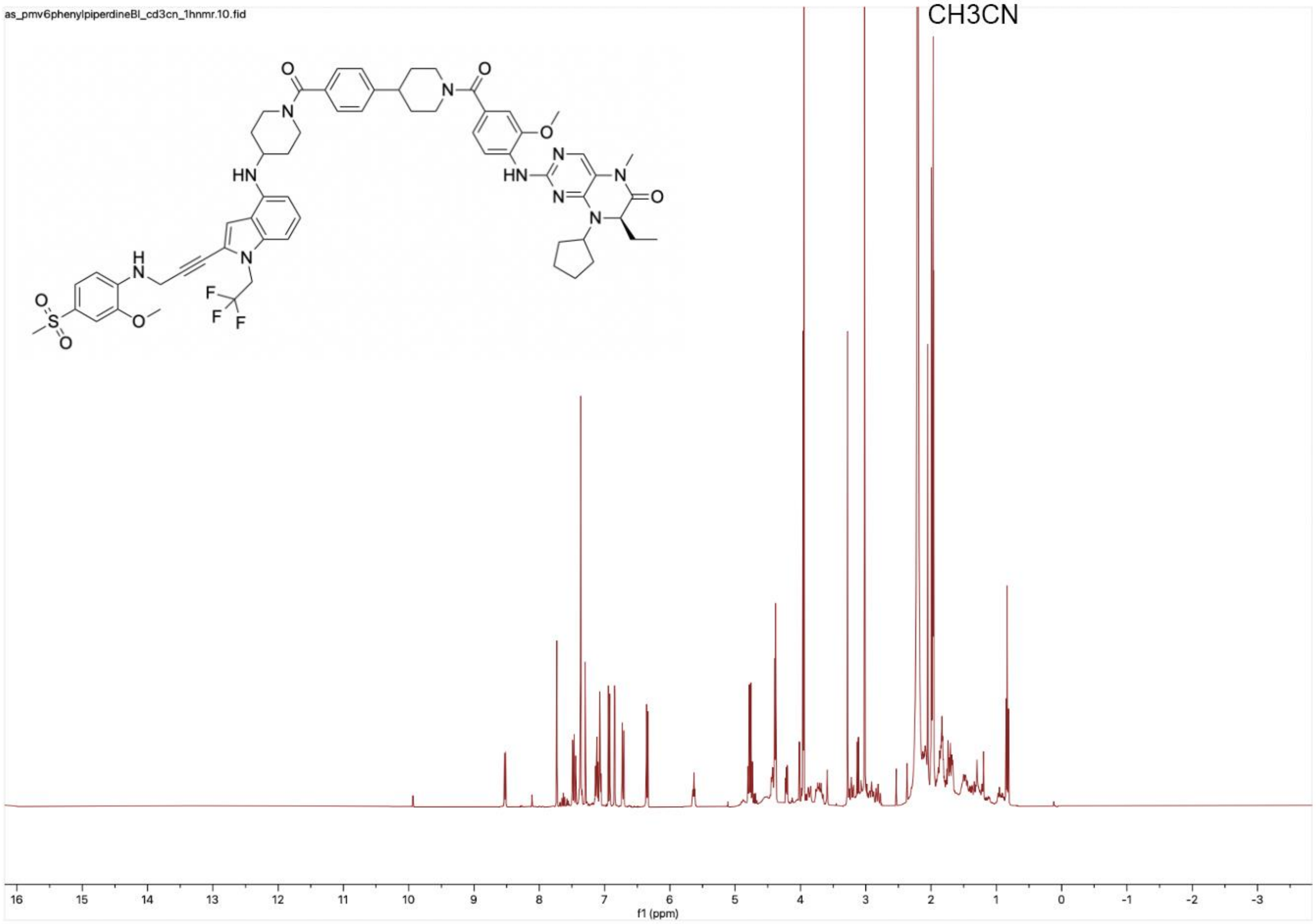
as_pmv6phenylpiperdineBI_cd3cn_1hnmr.10.fid
CH3CN
f1 (ppm)

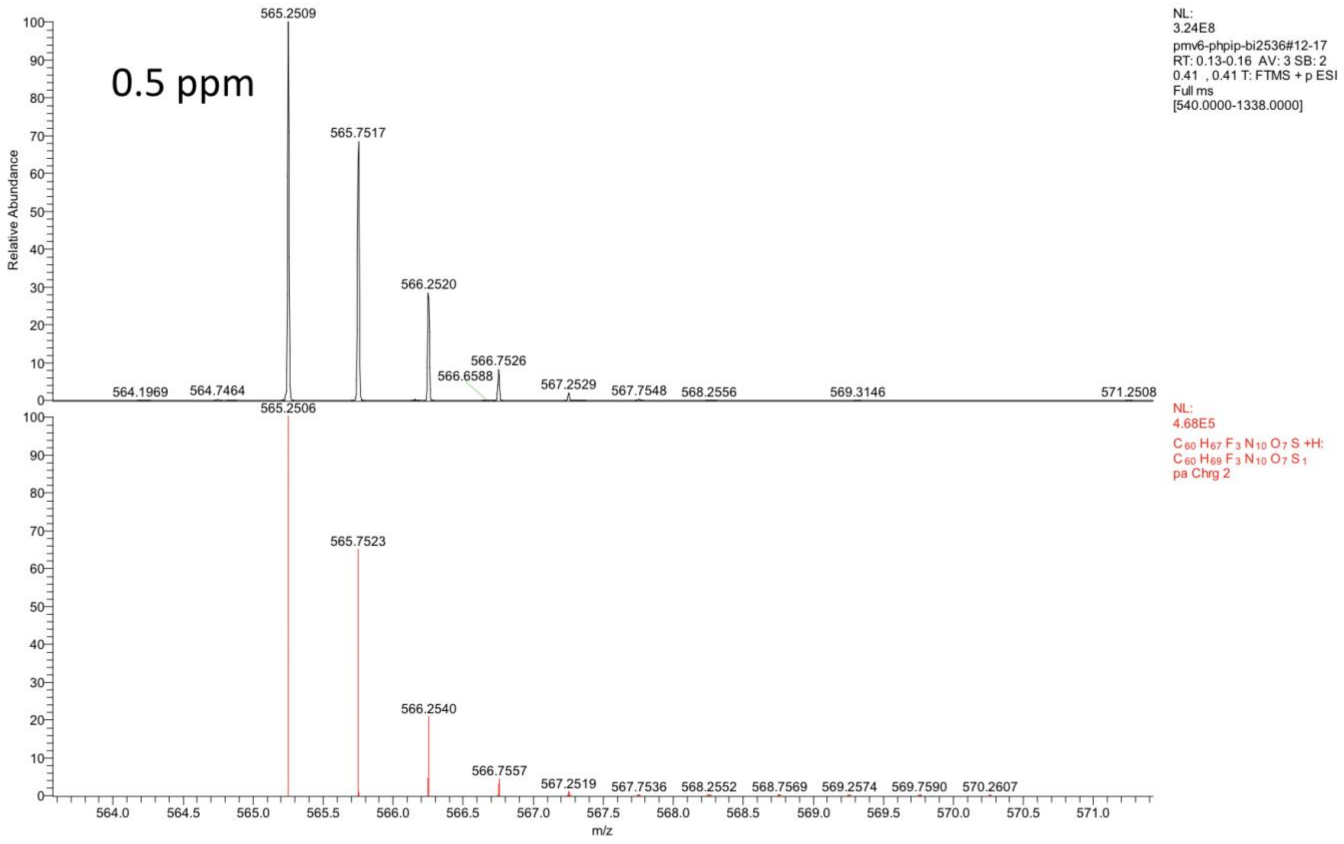

0.5 ppm
565.2509
565.7517
566.2520
566.6588
566.7526
567.2529
567.7548
568.2556
569.3146
571.2508
564.1969
564.7464
NL:
3.24E8
pmv6-phpip-bi2536#12-17
RT: 0.13-0.16 AV: 3 SB: 2
0.41 , 0.41 T: FTMS + p ESI
Full ms
[540.0000-1338.0000]
565.2506
565.7523
566.2540
566.7557
567.2519
567.7536
568.2552
568.7569
569.2574
569.7590
570.2607
NL:
4.68E5
C 60 H 67 F 3 N 10 O 7 S +H:
C 60 H 69 F 3 N 10 O 7 S 1
pa Chrg 2
Relative Abundance
m/z

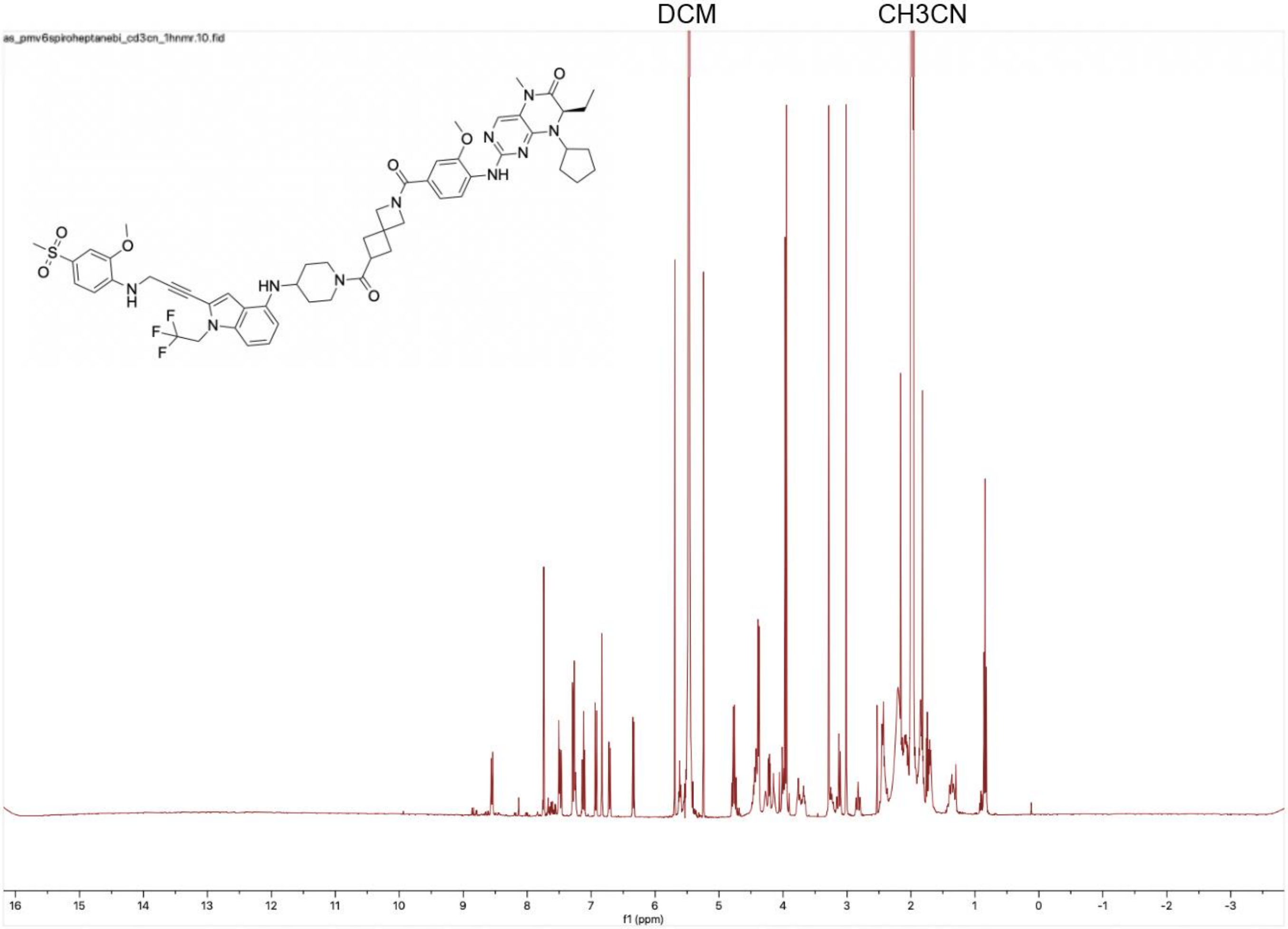
DCM
CH3CN
as_pmv6spiroheptanebi_cd3cn_1hnmr.10.fid
f1 (ppm)

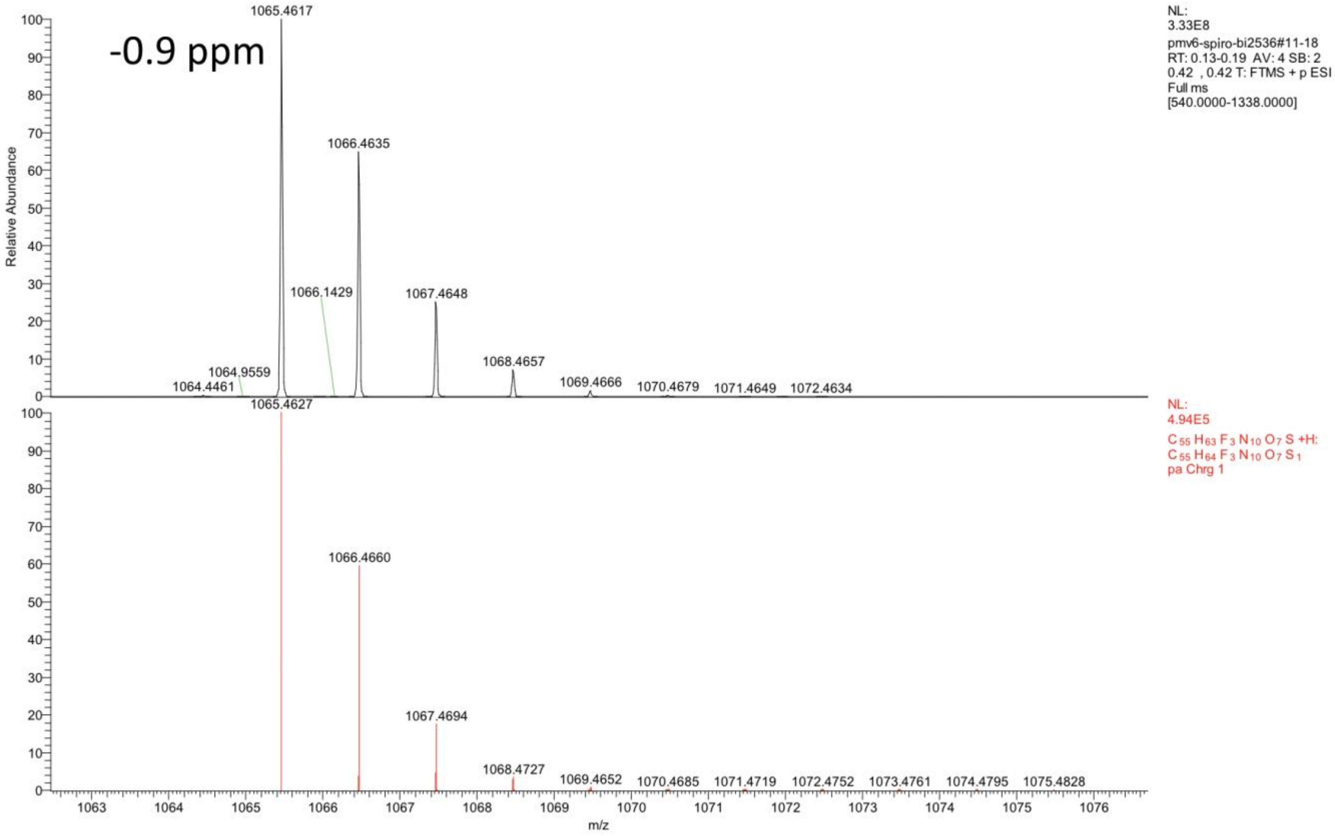

-0.9 ppm
1065.4617
1066.4635
1066.1429
1067.4648
1068.4657
1064.9559
1064.4461
1069.4666
1070.4679
1071.4649
1072.4634
1065.4627
1066.4660
1067.4694
1068.4727
1069.4652
1070.4685
1071.4719
1072.4752
1073.4761
1074.4795
1075.4828
Relative Abundance
m/z
NL:
3.33E8
pmv6-spiro-bi2536#11-18
RT: 0.13-0.19 AV: 4 SB: 2
0.42 , 0.42 T: FTMS + p ESI
Full ms
[540.0000-1338.0000]
NL:
4.94E5
C 55 H 63 F 3 N 10 O 7 S +H:
C 55 H 64 F 3 N 10 O 7 S 1
pa Chrg 1

as_pmv6c5bi_cd3cn_1hnmr.10.fid
Note: C5 is a typo in the name of this file, it is actually C6

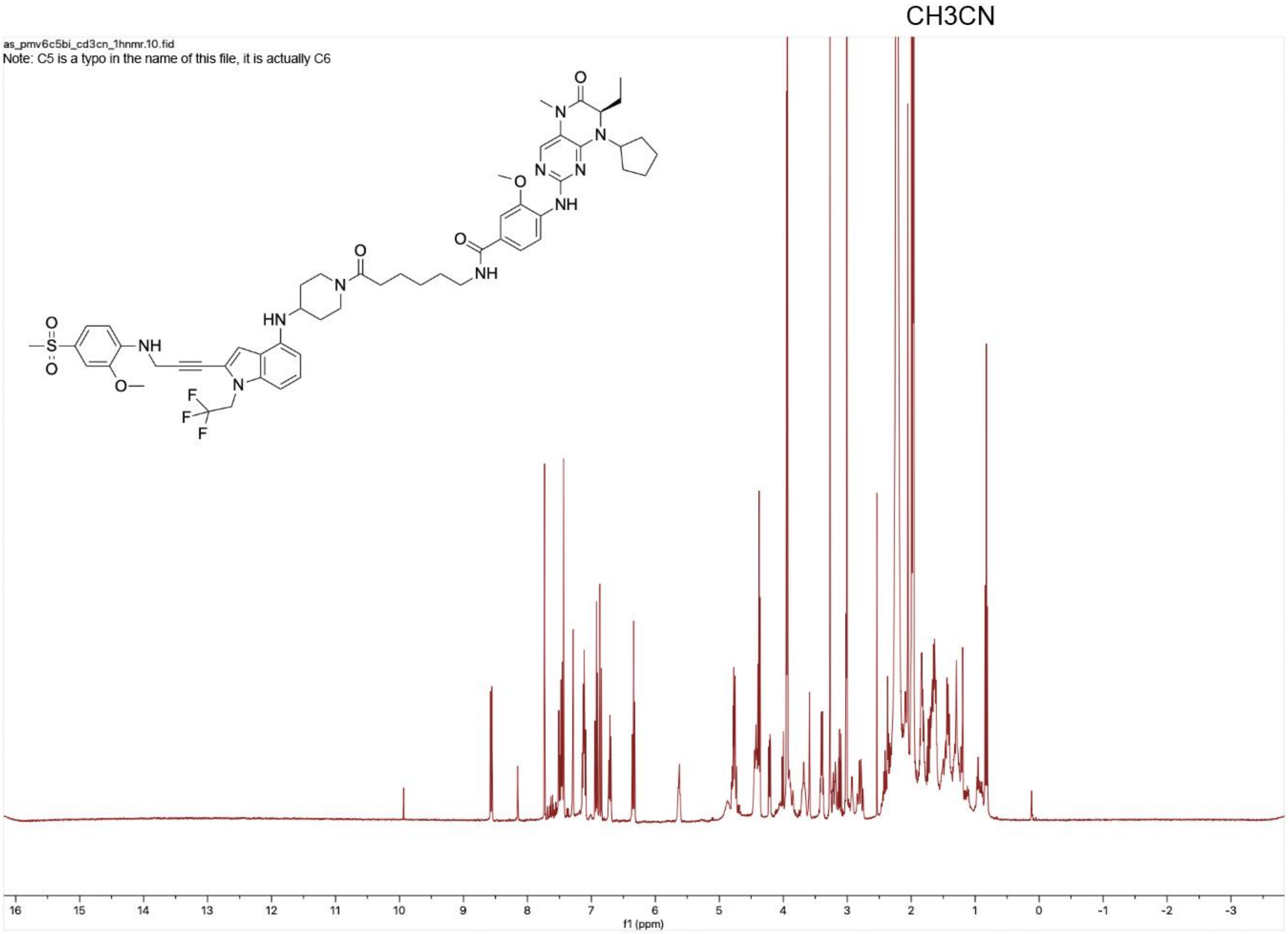

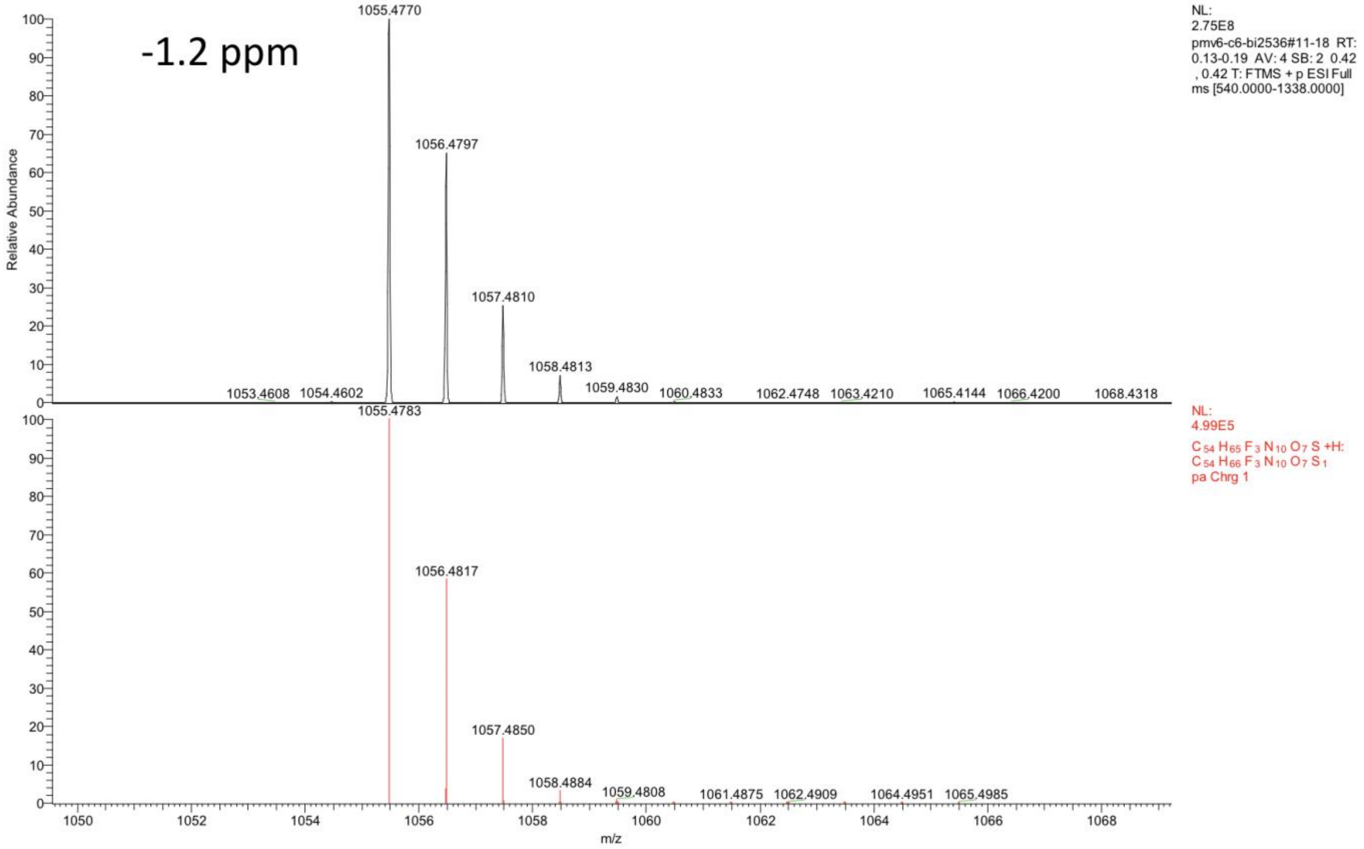
-1.2 ppm
1055.4770
1056.4797
1057.4810
1058.4813
1059.4830
1053.4608
1054.4602
1060.4833
1062.4748
1063.4210
1065.4144
1066.4200
1068.4318
NL:
2.75E8
pmv6-c6-bi2536#11-18 RT: 0.13-0.19 AV: 4 SB: 2 0.42 , 0.42 T: FTMS + p ESI Full ms [540.0000-1338.0000]
Relative Abundance
1055.4783
1056.4817
1057.4850
1058.4884
1059.4808
1061.4875
1062.4909
1064.4951
1065.4985
NL:
4.99E5
$C_{54}H_{65}F_3N_{10}O_7S$ +H:
$C_{54}H_{66}F_3N_{10}O_7S_1$
pa Chrg 1
m/z

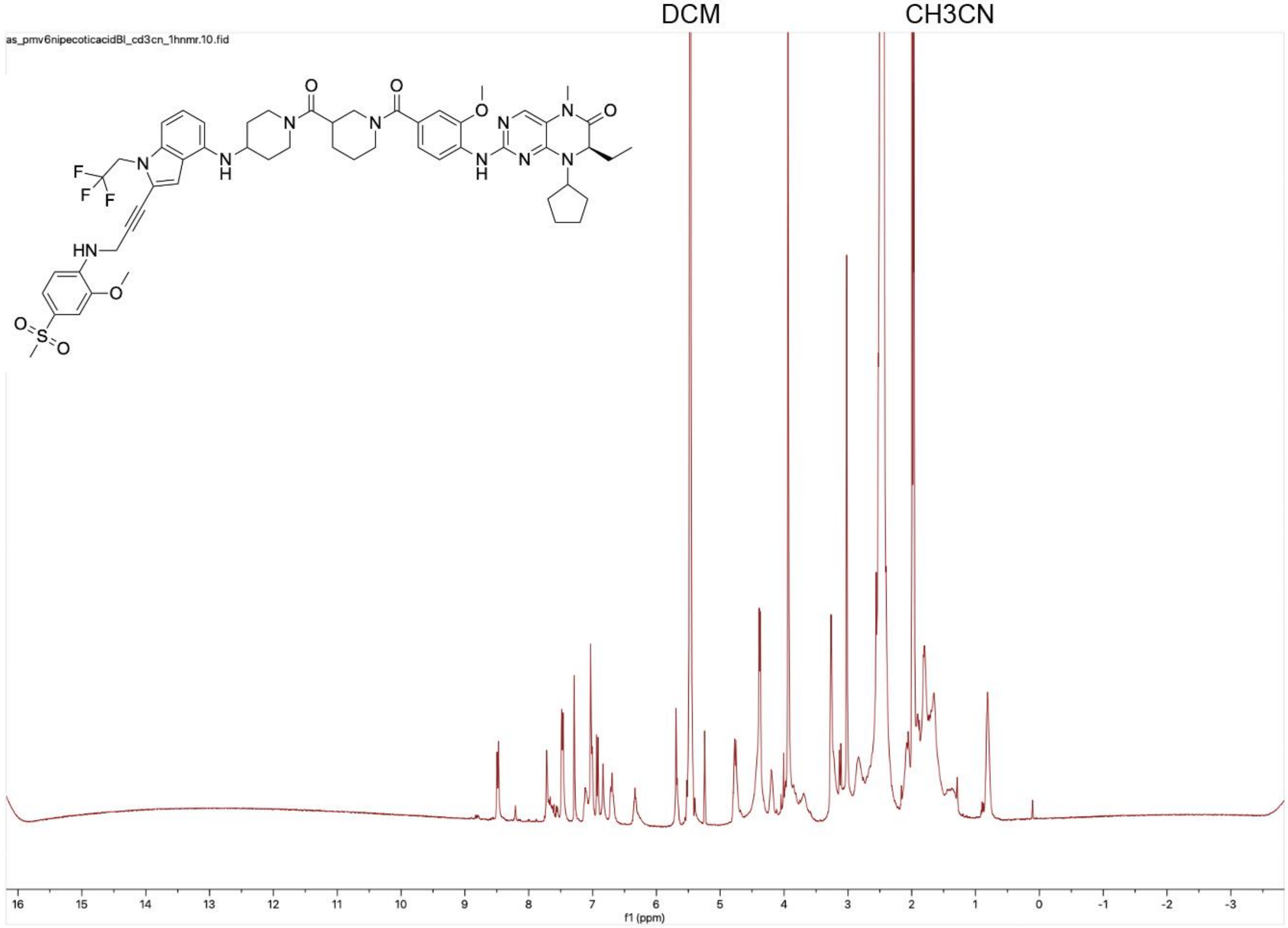
DCM
CH3CN
as_pmv6nipecoticacidBl_cd3cn_1hnmr.10.fid
f1 (ppm)

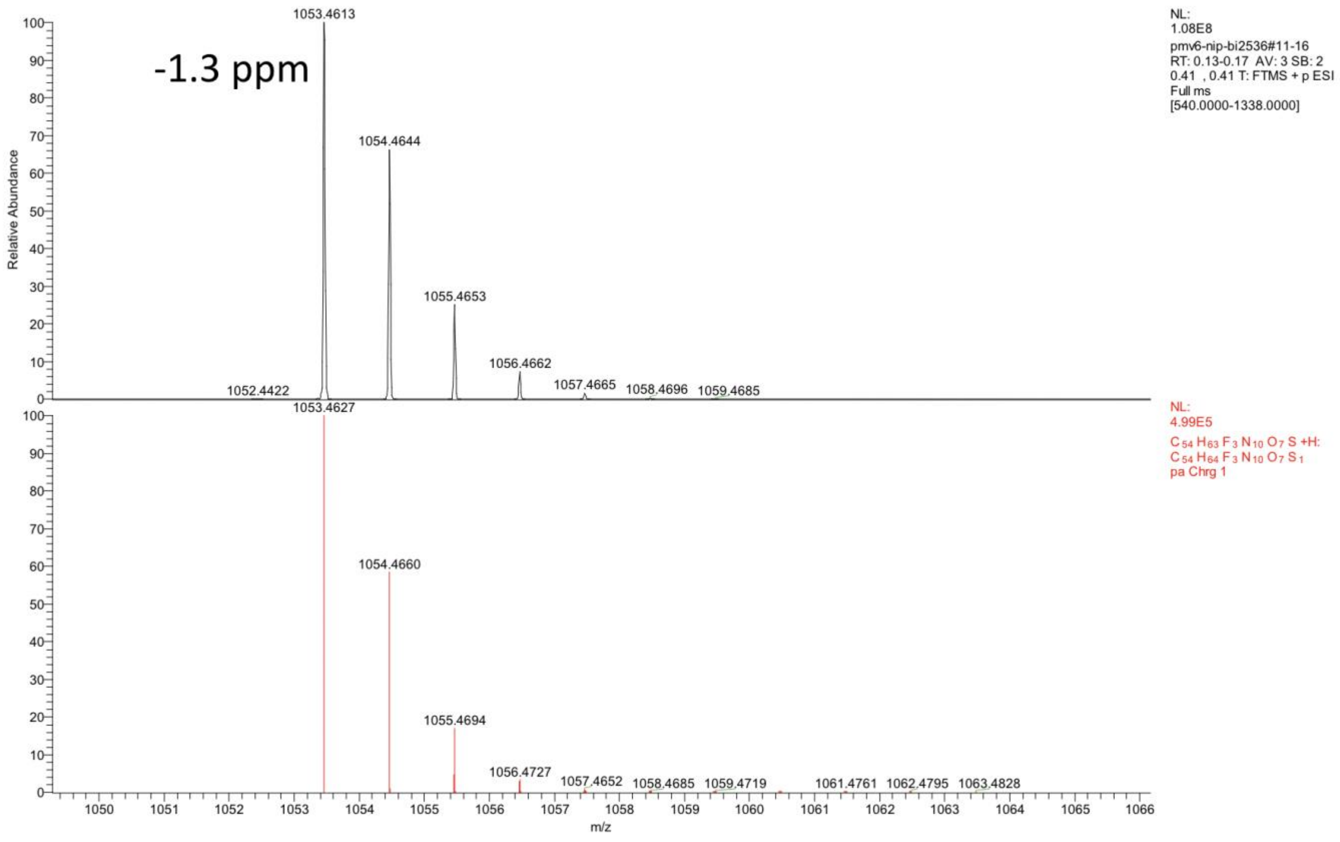

-1.3 ppm
1053.4613
1054.4644
1055.4653
1056.4662
1057.4665
1058.4696
1059.4685
1052.4422
NL:
1.08E8
pmv6-nip-bi2536#11-16
RT: 0.13-0.17 AV: 3 SB: 2
0.41 , 0.41 T: FTMS + p ESI
Full ms
[540.0000-1338.0000]
1053.4627
1054.4660
1055.4694
1056.4727
1057.4652
1058.4685
1059.4719
1061.4761
1062.4795
1063.4828
NL:
4.99E5
$C_{54}H_{63}F_3N_{10}O_7S$ +H:
$C_{54}H_{64}F_3N_{10}O_7S_1$
pa Chrg 1
Relative Abundance
m/z

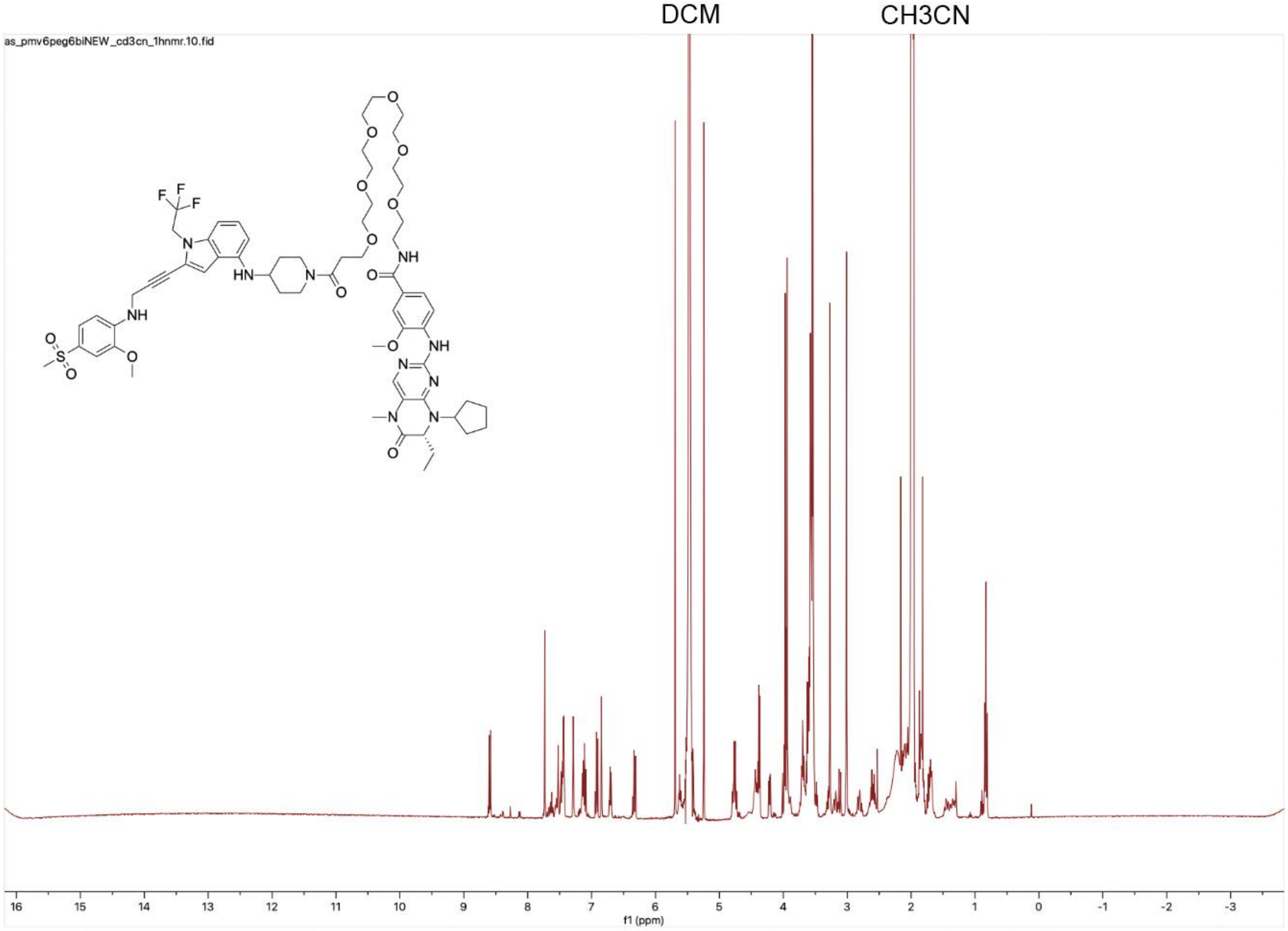
DCM
CH3CN
as_pmv6peg6biNEW_cd3cn_1hnmr.10.fid
f1 (ppm)

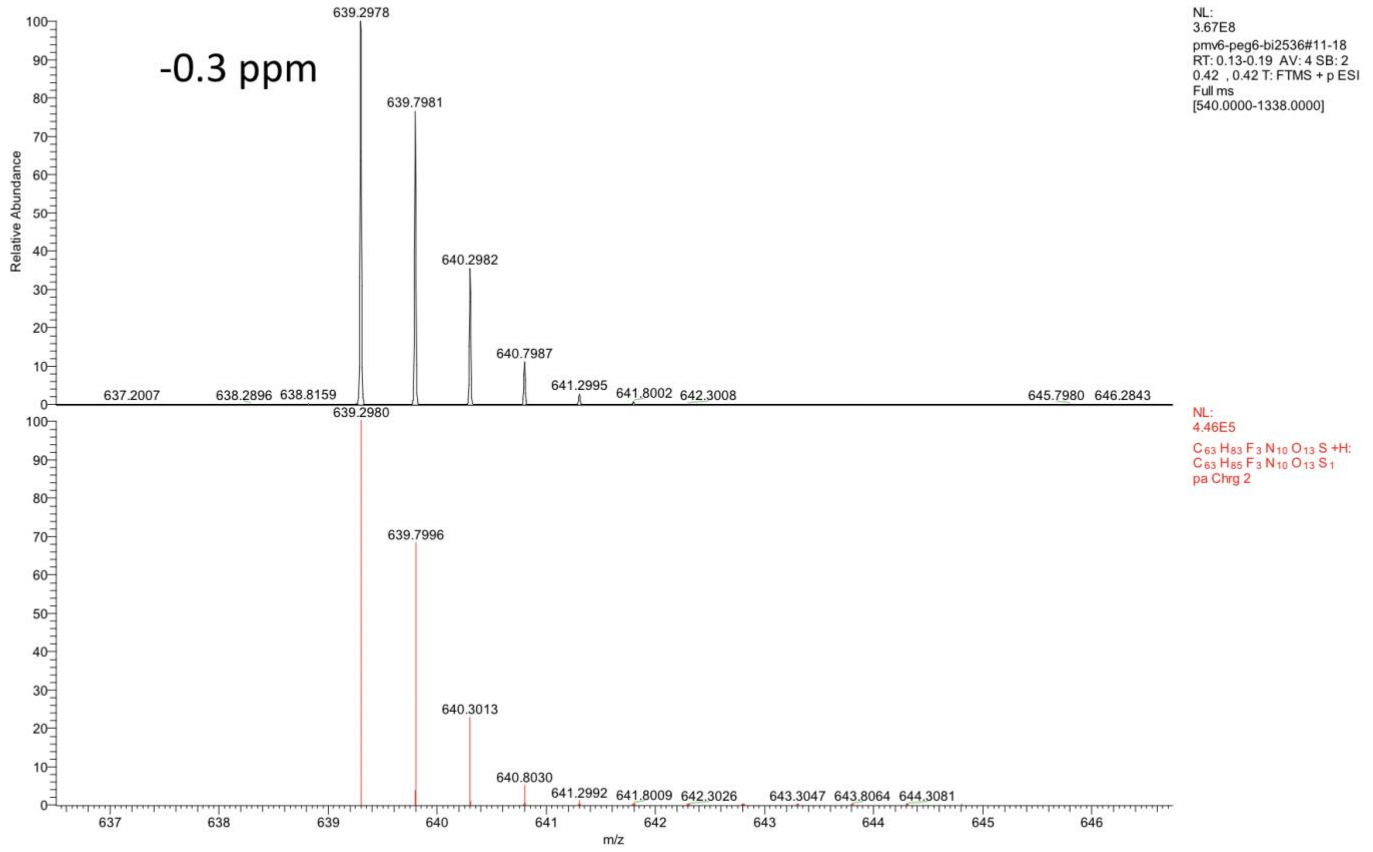
-0.3 ppm
NL:
3.67E8
pmv6-peg6-bi2536#11-18
RT: 0.13-0.19 AV: 4 SB: 2
0.42 , 0.42 T: FTMS + p ESI
Full ms
[540.0000-1338.0000]
639.2978
639.7981
640.2982
640.7987
641.2995
641.8002
642.3008
637.2007
638.2896
638.8159
645.7980
646.2843
NL:
4.46E5
C 63 H 83 F 3 N 10 O 13 S +H:
C 63 H 85 F 3 N 10 O 13 S 1
pa Chrg 2
639.2980
639.7996
640.3013
640.8030
641.2992
641.8009
642.3026
643.3047
643.8064
644.3081
Relative Abundance
m/z

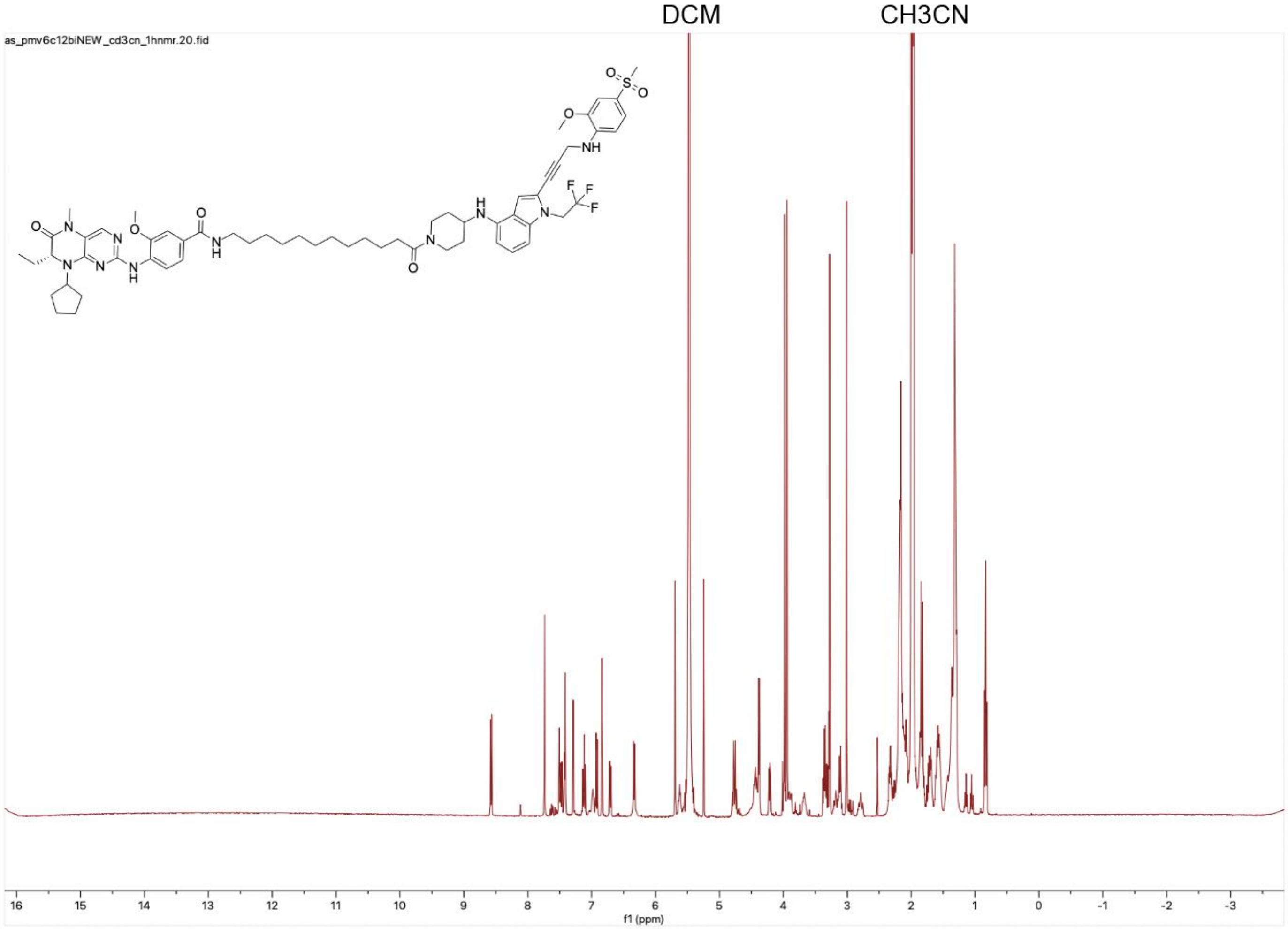

DCM
CH3CN
as_pmv6c12biNEW_cd3cn_1hnmr.20.fid
f1 (ppm)

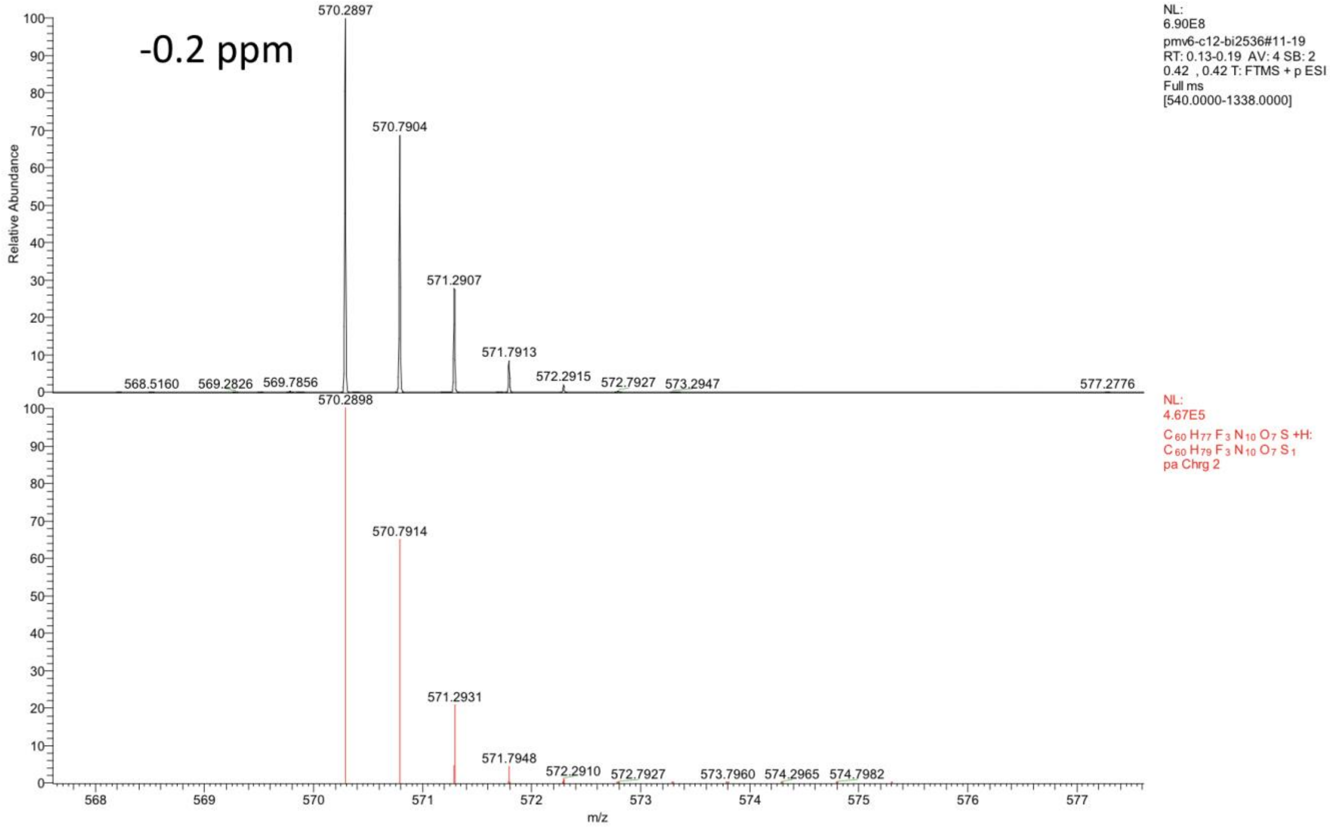
-0.2 ppm
570.2897
570.7904
571.2907
571.7913
572.2915
572.7927
573.2947
568.5160
569.2826
569.7856
577.2776
NL:
6.90E8
pmv6-c12-bi2536#11-19
RT: 0.13-0.19 AV: 4 SB: 2
0.42 , 0.42 T: FTMS + p ESI
Full ms
[540.0000-1338.0000]
570.2898
570.7914
571.2931
571.7948
572.2910
572.7927
573.7960
574.2965
574.7982
NL:
4.67E5
C 60 H 77 F 3 N 10 O 7 S +H:
C 60 H 79 F 3 N 10 O 7 S 1
pa Chrg 2
Relative Abundance
m/z

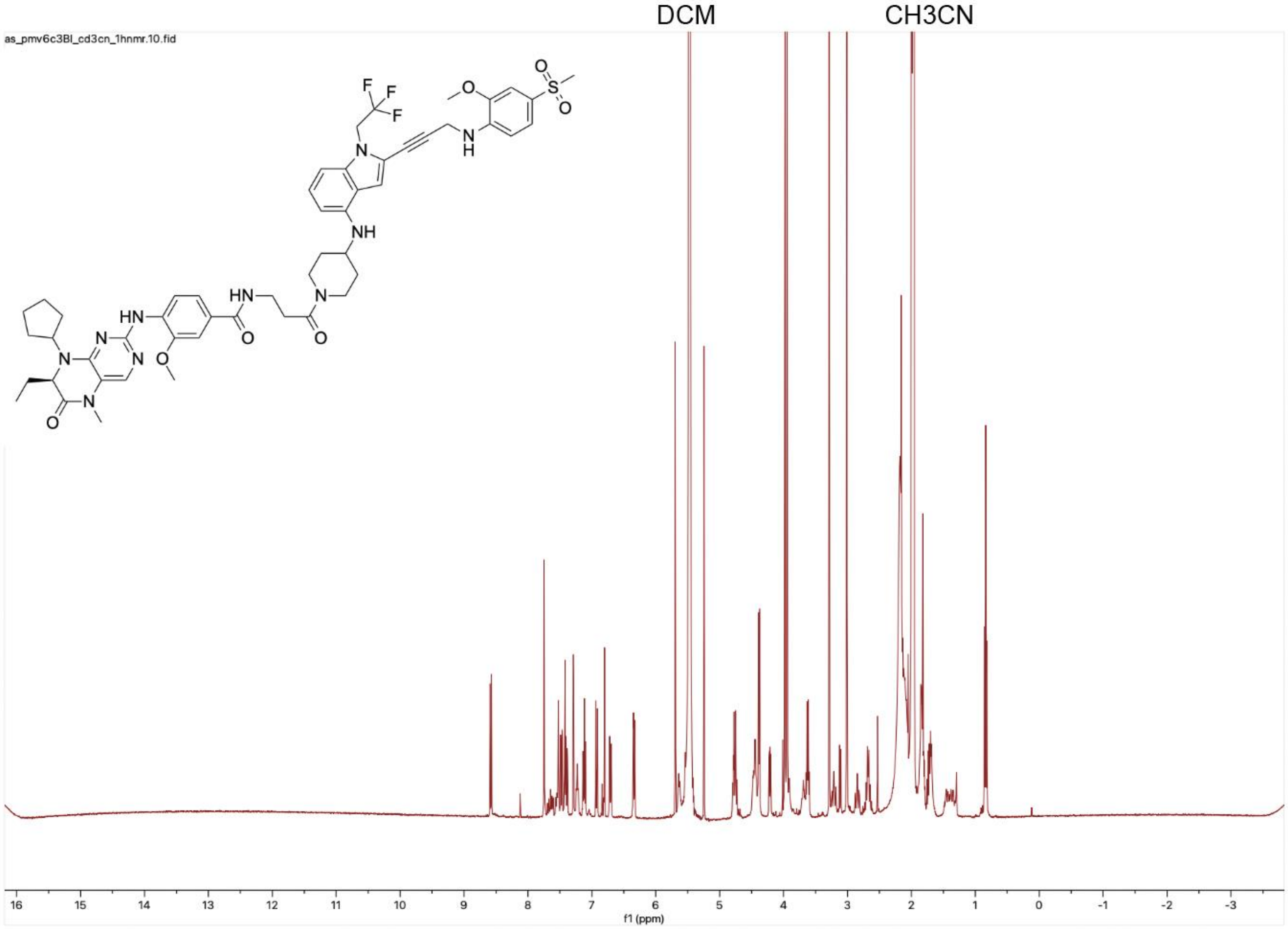

DCM
CH3CN
as_pmv6c3BI_cd3cn_1hnmr.10.fid
16 15 14 13 12 11 10 9 8 7 6 5 4 3 2 1 0 -1 -2 -3
f1 (ppm)

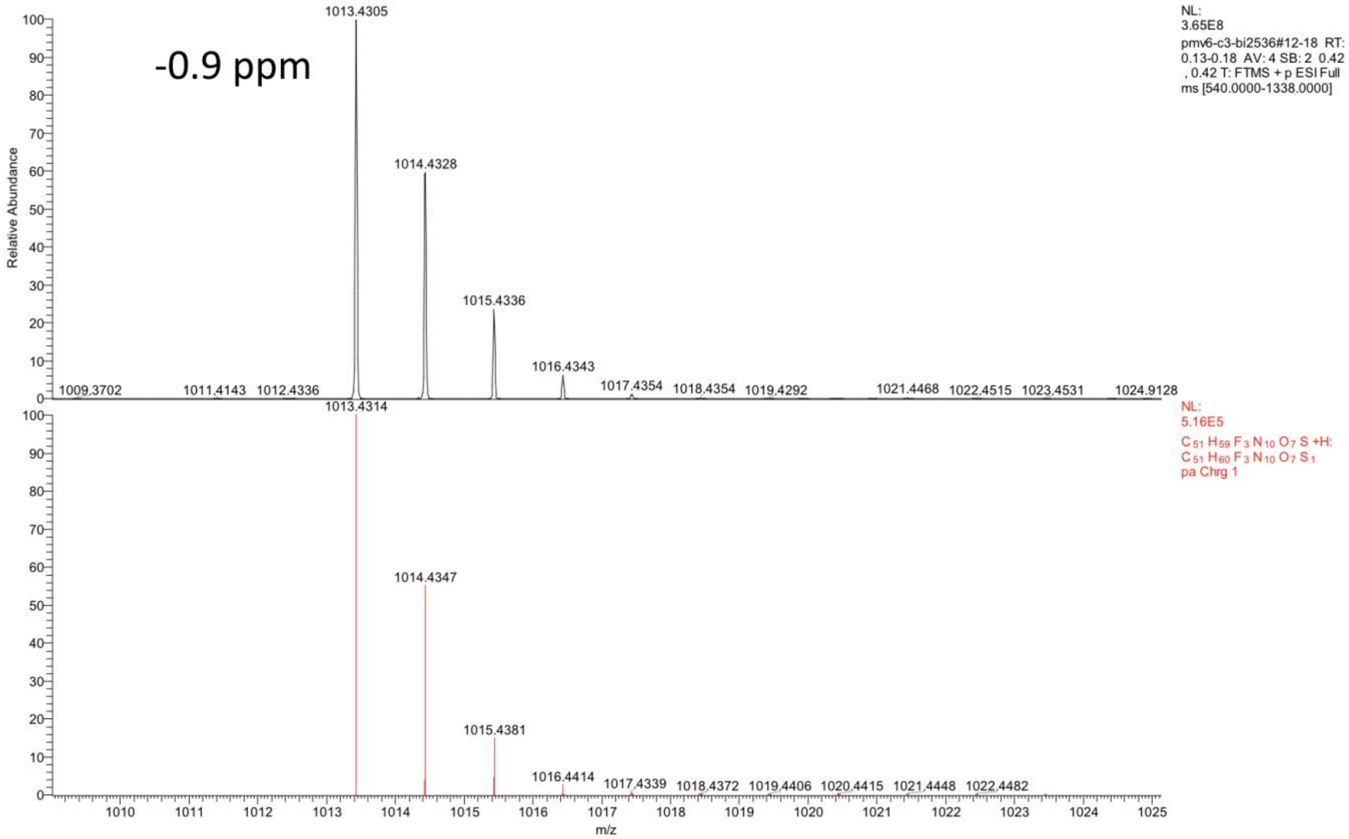


*PMV6-PEG4-Adavosertib.*

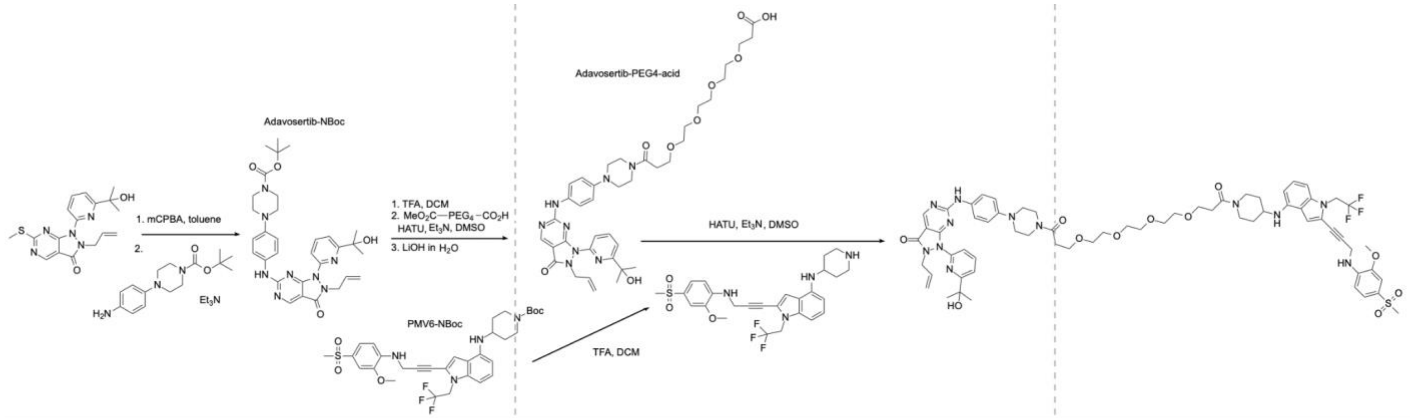


**1-(4-(4-((2-allyl-1-(6-(2-hydroxypropan-2-yl)pyridin-2-yl)-3-oxo-2,3-dihydro-1H-pyrazolo[3,4-d]pyrimidin-6-yl)amino)phenyl)piperazin-1-yl)-16-(4-((2-(3-((2-methoxy-4-(methylsulfonyl)phenyl)amino)prop-1-yn-1-yl)-1-(2,2,2-trifluoroethyl)-1H-indol-4-yl)amino)piperidin-1-yl)-4,7,10,13-tetraoxahexadecane-1,16-dione (PMV6-PEG4-Adavosertib, PMV6-PEG4-Ada).** 2-allyl-1-(6-(2-hydroxypropan-2-yl)pyridin-2-yl)-6-(methylthio)-1,2-dihydro-3H-pyrazolo[3,4-d]pyrimidin-3-one (50 mg, 0.140 mmol, AaronChem) was dissolved in 4 mL of toluene and 3-chloroperbenzoic acid (32 mg, 0.185 mmol) was added and stirred for 1 hour. 1-Boc-4-(4-aminophenyl)piperazine (46.6 mg, 0.168 mmol, Combi-Blocks) and triethylamine (100 μL, 0.718 mmol) were added to the solution and left to stir overnight. The reaction was purified by reverse-phase HPLC (acetonitrile:water gradient up to 90%) to yield adavosertib-NBoc as a green-brown solid (38 mg, 0.0648 mmol, 46.3% yield). Some of the resulting solid (30 mg, 0.0512 mmol) was dissolved in DCM (1 mL), and trifluoroacetic acid was added to the solution (1.5 mL). After stirring for 1 hour, solvent and reactants were removed under reduced pressure. The resulting residue was dissolved in DMSO (2 mL); HATU (40 mg, 0.105 mmol), 3-oxo-2,6,9,12,15-pentaoxaoctadecan-18-oic acid (acid-PEG4-mono-methyl ester, 50 µL, 0.162 mmol), and triethylamine (50 µL, 0.359 mmol) were added to the solution and the mixture was stirred for 30 minutes. LiOH (2M solution in $H_2O$, 2 mL) was added to the solution and left to stir for 1 hour. The reaction was purified by reverse-phase HPLC (acetonitrile:water gradient up to 90%) to yield adavosertib-PEG4-acid as a light yellow solid (25 mg, 0.0328 mmol, 64.0%). *tert*-butyl-4-((2-(3-((2-methoxy-4-(methylsulfonyl)phenyl)amino)prop-1-yn-1-yl)-1-(2,2,2-trifluoroethyl)-1H-indol-4-yl)amino)piperidine-1-carboxylate (PMV6-NBoc, 15.1 mg, 0.0238 mmol) was dissolved in DCM (1 mL), and trifluoroacetic acid (2 mL) was added to the solution. After stirring for 1 hour, solvents were removed under reduced pressure. The residue was redissolved in DMSO (1 mL) and adavosertib-PEG4-acid (25 mg, 0.0328 mmol), HATU (20 mg, 0.0526 mmol), and triethylamine (40 µL, 0.287 mmol) were added to the solution. After completion (30 minutes), the reaction was purified by reverse-phase HPLC (acetonitrile:water gradient up to 90%) to yield PMV6-PEG4-Adavosertib as a yellow solid (15.3 mg, 0.0120 mmol, 50.4% yield). $^{1}$H NMR (400 MHz, DMSO-d6) δ 8.83 (s, 1H), 8.04 (t, J = 7.9 Hz, 1H), 7.76 (d, J = 8.1 Hz, 1H), 7.61 (t, J = 6.8 Hz, 3H), 7.40 (dd, J = 8.4, 1.9 Hz, 1H), 7.26 (d, J = 2.0 Hz, 1H), 7.10 – 6.86 (m, 4H), 6.71 (d, J = 8.3 Hz, 1H), 6.48 (t, J = 6.3 Hz, 1H), 6.23 (d, J = 7.8 Hz, 1H), 5.67 (ddt, J = 16.5, 10.2, 6.0 Hz, 1H), 5.51 (d, J = 7.9 Hz, 1H), 5.32 (s, 1H), 5.03 - 4.79 (m, 3H), 4.70 (d, J = 6.0 Hz, 1H), 4.35 (q, J = 5.8 Hz, 6H), 3.90 (s, 2H), 3.69 – 3.57 (m, 6H), 3.52 – 3.38 (m, 13H), 3.20 – 3.02 (m, 4H), 2.76 (t, J = 11.8 Hz, 1H), 2.60 (dt, J = 17.2, 6.9 Hz, 4H), 2.04 – 1.90 (m, 2H), 1.48 (s, 4H), 1.41 (d, J = 6.4 Hz, 4H), 1.36 – 1.14 (m, 1H), 1.30 (d, J = 11.5 Hz, 1H), 1.07 (t, J = 7.0 Hz, 7H). $^{13}$C NMR (101 MHz, DMSO-d6) δ 169.30, 168.97, 168.06, 161.62, 161.46, 160.95, 156.45, 147.47 (d, *J* = 15.8 Hz), 146.58, 141.83, 140.98, 139.23, 138.29, 132.25 (d, *J* = 80.3 Hz), 129.05 – 124.63 (m), 122.57 (d, *J* = 164.8 Hz), 118.63 (d, *J* = 8.3 Hz), 118.46, 116.75, 116.58, 116.15, 109.09, 108.21, 107.60, 100.04, 99.08, 93.60, 73.75, 72.78, 71.51 – 68.75 (m), 67.32 (d, *J* = 8.0 Hz), 56.36 (d, *J* = 29.1 Hz), 51.20 – 48.17 (m), 47.05, 46.20 – 42.94 (m), 41.32, 39.16, 33.27 (d, *J* = 5.2 Hz), 32.93, 32.66, 31.88, 31.34, 30.87, 28.11 (d, *J* = 7.4 Hz), 18.97. HRMS (top: observed, bottom: theoretical isotope pattern; M+H, $C_{64}H_{77}F_3N_{12}O_{11}S$+2H] m/z theoretical 640.2827, found 640.2825.

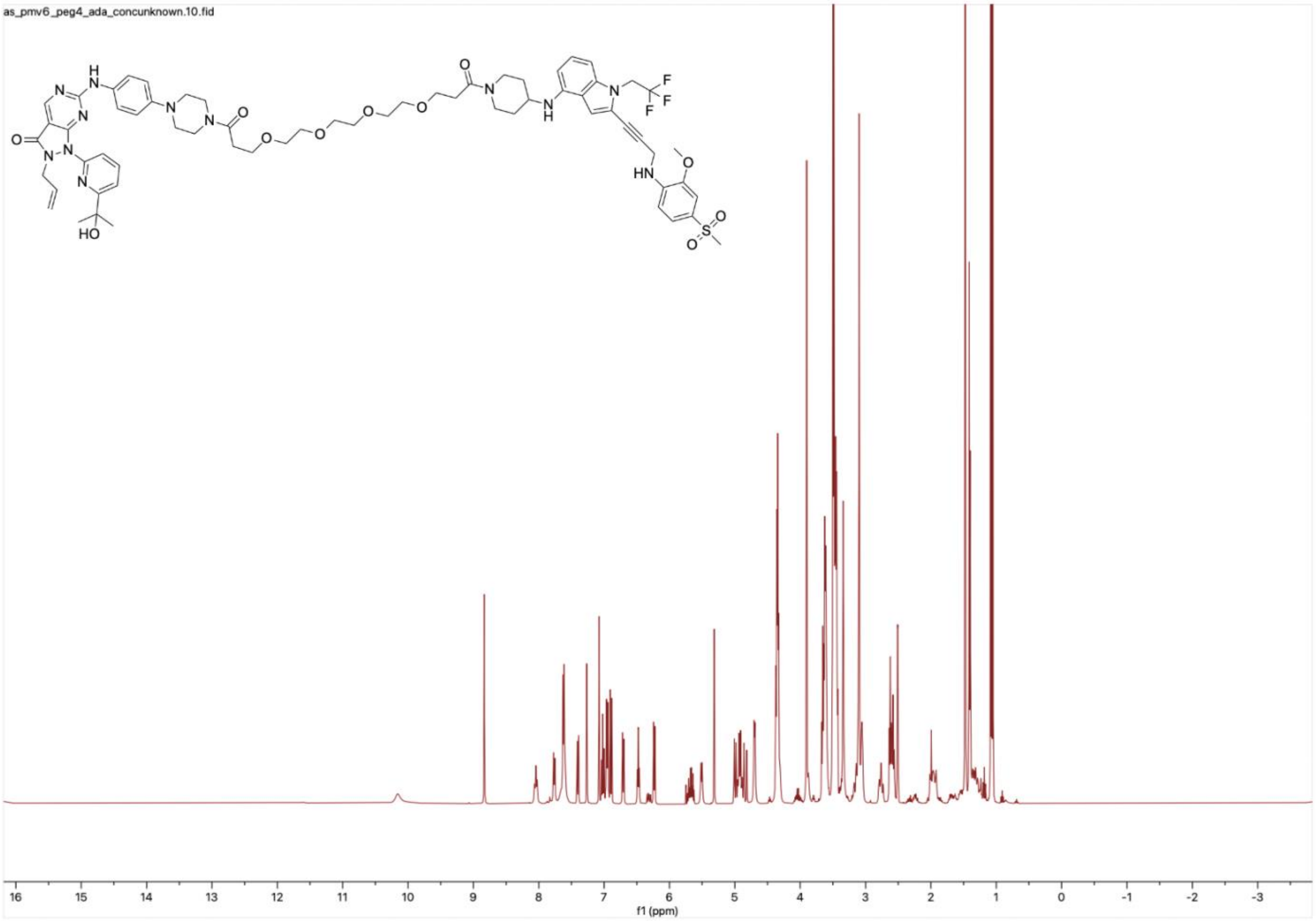
as_pmv6_peg4_ada_concunknown.10.fid
16 15 14 13 12 11 10 9 8 7 6 5 4 3 2 1 0 -1 -2 -3
f1 (ppm)

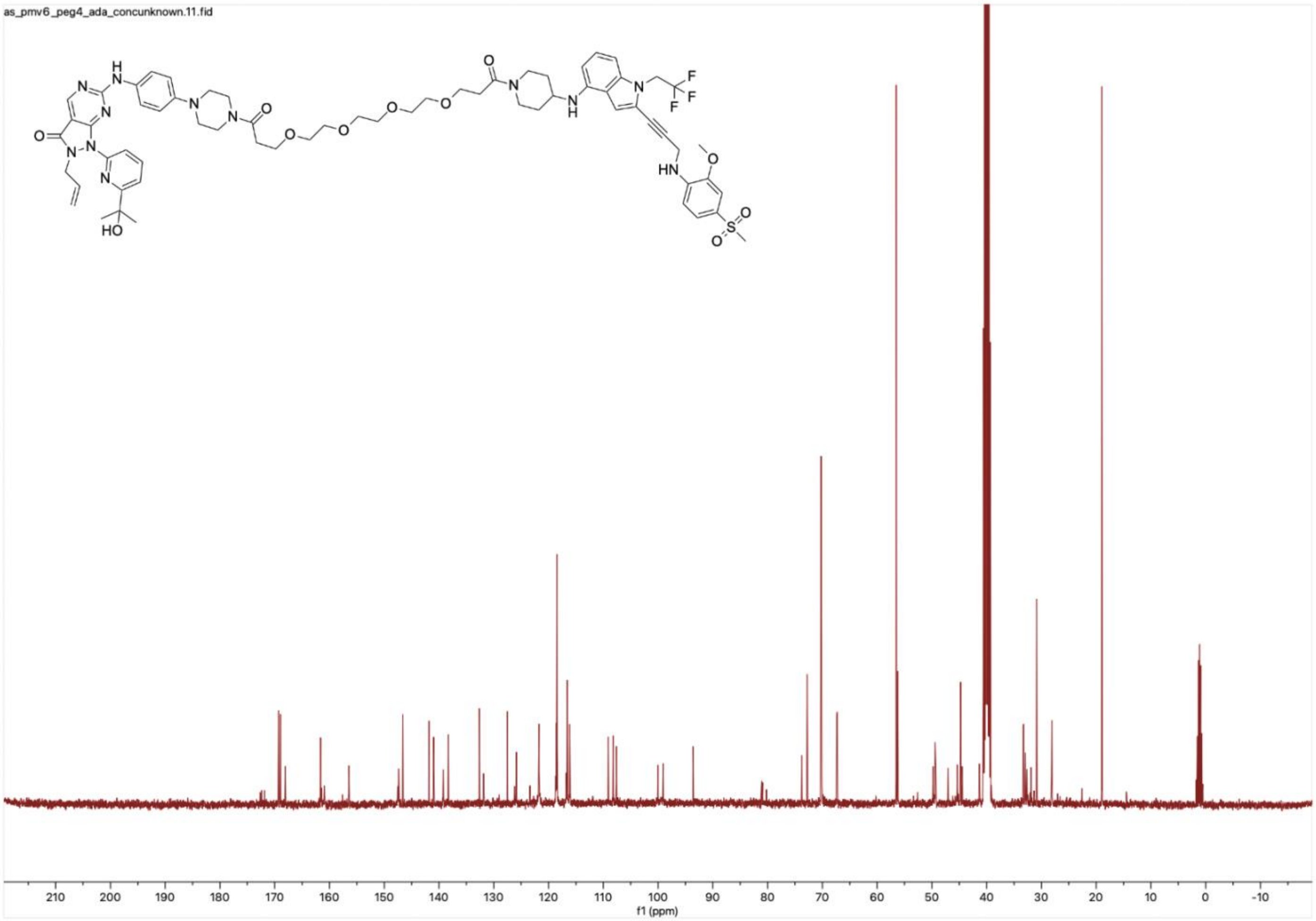
as_pmv6_peg4_ada_concunknown.11.fid
210
200
190
180
170
160
150
140
130
120
110
100
90
80
70
60
50
40
30
20
10
0
-10
f1 (ppm)

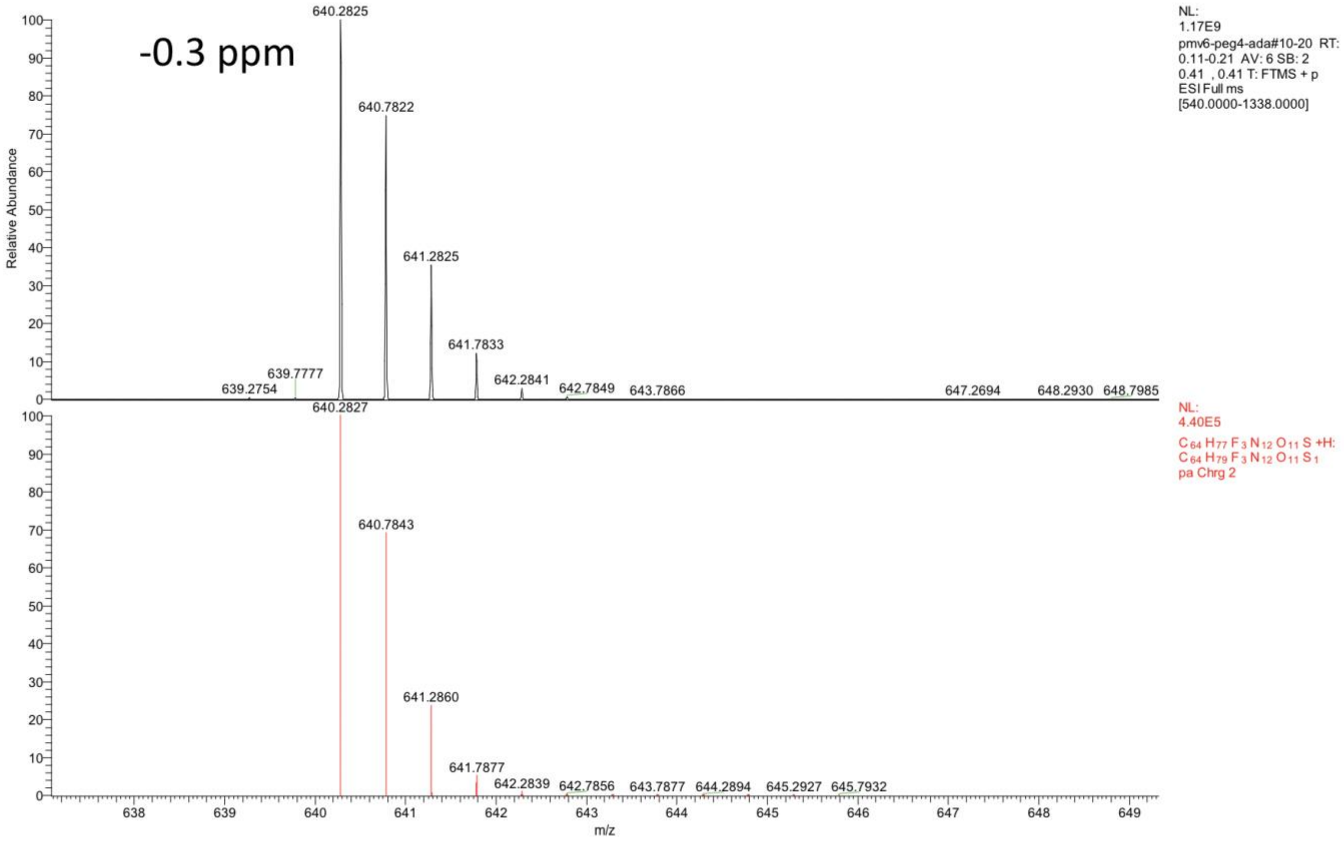


*BI2536-C8-AP1867.*

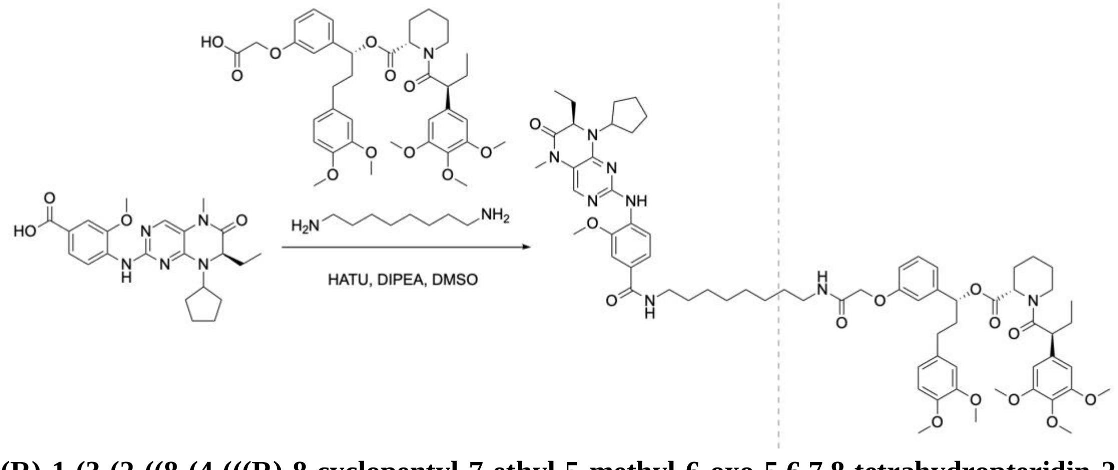


**(R)-1-(3-(2-((8-(4-(((R)-8-cyclopentyl-7-ethyl-5-methyl-6-oxo-5,6,7,8-tetrahydropteridin-2-yl)amino)-3-methoxybenzamido)octyl)amino)-2-oxoethoxy)phenyl)-3-(3,4-dimethoxyphenyl)propyl-(S)-1-((S)-2-(3,4,5-trimethoxyphenyl)butanoyl)piperidine-2-carboxylate (BI2536-C8-AP1867)**. To a solution of (R)-4-(8-cyclopentyl-7-ethyl-5-methyl-6-oxo-5,6,7,8-tetrahydropteridin-2-ylamino)-3-methoxybenzoic acid (4 mg, 0.00941 mmol, 1ClickChemistry) in DMSO (1 mL), meta-AP1867 (6.5 mg, 0.00937 mmol, MedChemExpress), 1,8-diaminooctane (1.4 μL, 0.00952 mmol), HATU (8 mg, 0.0205 mmol), and DIPEA (20 μL, 0.115 mmol) were added to the solution. After completion (30 minutes), the reaction was purified by reverse-phase HPLC (acetonitrile:water gradient up to 90%) to yield BI2536-C8-AP1867 as an off-white solid (3.2 mg, 0.00261 mmol, 27.7% yield). A stock solution was created in DMSO-d6, and a portion of the stock solution was set aside for $^{1}$H NMR characterization. Due to mixing with residual DCM in the NMR tube from cleaning, the portion of the stock that underwent NMR was discarded. [1]H NMR (400 MHz, DMSO-d6) spectrum is shown below. HRMS (top: observed, bottom: theoretical isotope pattern; M+H, $C_{68}H_{90}N_8O_{13}$+H] m/z theoretical 1227.6700, found 1227.6691.

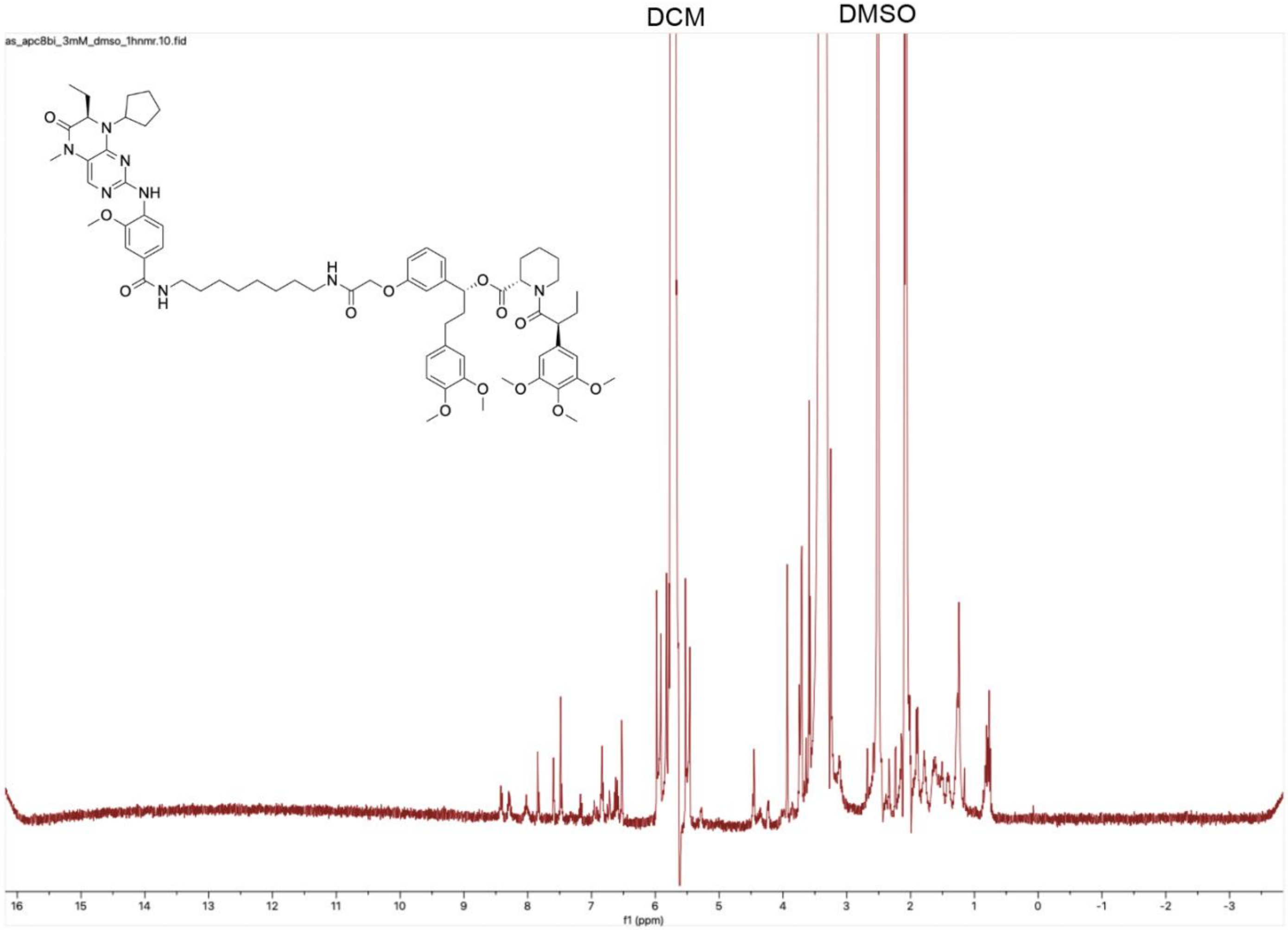

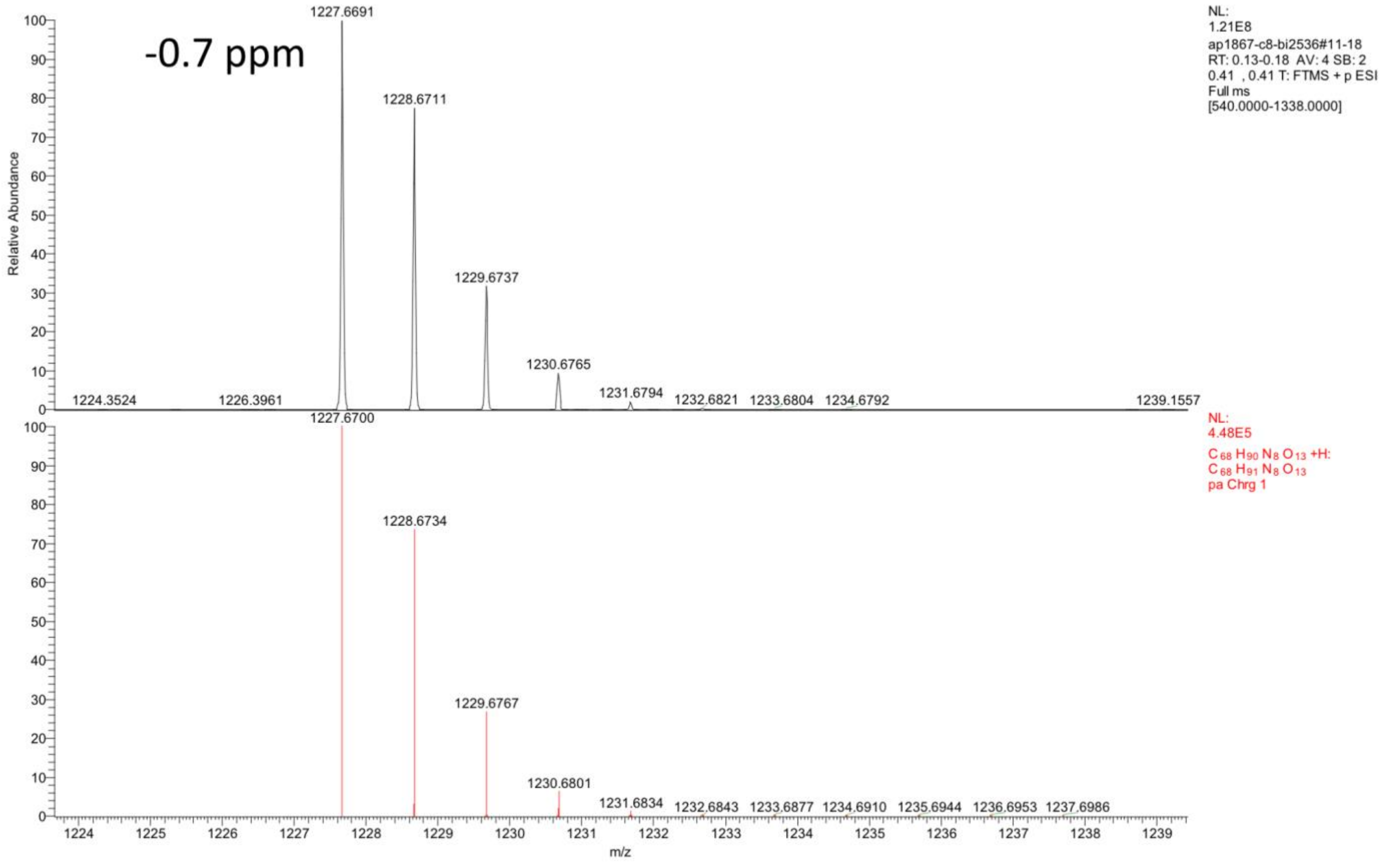


*KG5-PEG4-BI2536.*

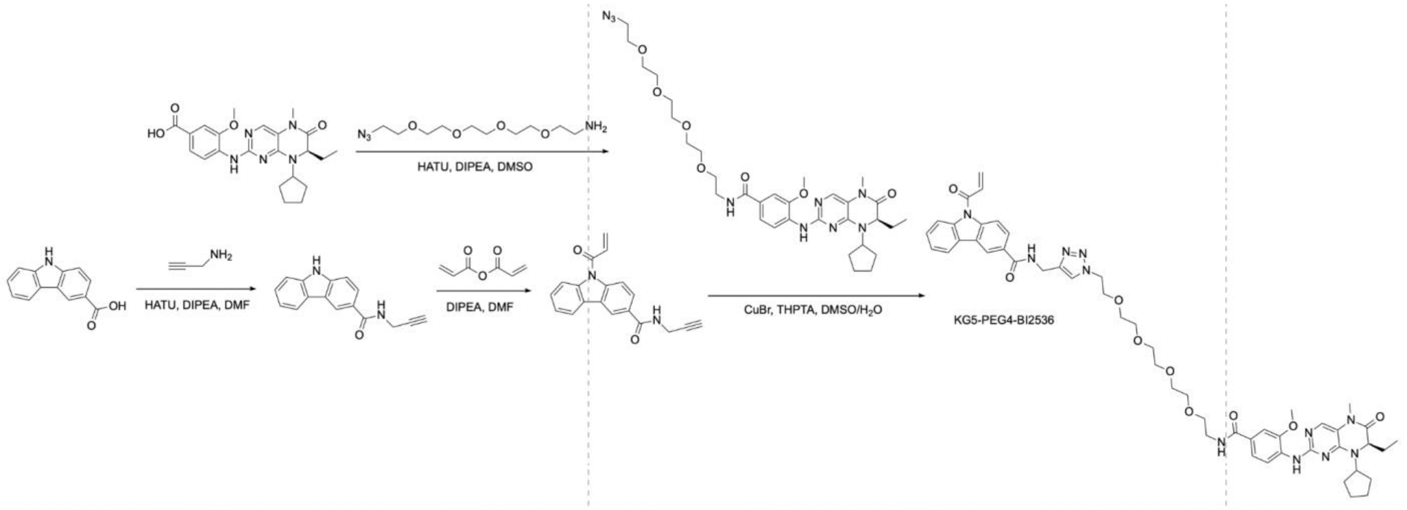


**(R)-9-acryloyl-N-((1-(1-(4-((8-cyclopentyl-7-ethyl-5-methyl-6-oxo-5,6,7,8-tetrahydropteridin-2-yl)amino)-3-methoxyphenyl)-1-oxo-5,8,11,14-tetraoxa-2-azahexadecan-16-yl)-1H-1,2,3-triazol-4-yl)methyl)-9H-carbazole-3-carboxamide (KG5-PEG4-BI2536, KG5-BI2536).** To a solution of 9H-carbazole-3-carboxylic acid (25 mg, 0.118 mmol, 1ClickChemistry) in DMF (1 mL), propargylamine (20 µL, 0.313 mmol), HATU (50 mg, 0.128 mmol), and DIPEA (50 µL, 0.288 mmol) were added to the solution. After completion (30 minutes), the reaction was purified by reverse-phase HPLC (acetonitrile:water gradient up to 90%) to yield N-(prop-2-yn-1-yl)-9H-carbazole-3-carboxamide as a white solid (20 mg, 0.0806 mmol, 68.3%). The resulting solid was dissolved in DMF (3 mL), and acrylic anhydride (60 µL, 0.521 mmol) and DIPEA (120 µL, 0.691 mmol) were added to the solution to react overnight. After completion, the reaction was purified by reverse-phase HPLC (acetonitrile:water gradient up to 90%) to yield 9-acryloyl-N-(prop-2-yn-1-yl)-9H-carbazole-3-carboxamide as a off-white solid (8.1 mg, 0.0268 mmol, 33.2% yield). To a solution of (R)-4-(8-cyclopentyl-7-ethyl-5-methyl-6-oxo-5,6,7,8-tetrahydropteridin-2-ylamino)-3-methoxybenzoic acid (7 mg, 0.0165 mmol, 1ClickChemistry) in DMSO (1 mL), $N_3$-PEG4-$NH_2$ (10 µL, 0.0382 mmol), HATU (8 mg, 0.0205 mmol), and DIPEA (20 µL, 0.115 mmol) were added to the solution. After completion (30 minutes), the reaction was purified by reverse-phase HPLC (acetonitrile:water gradient up to 90%) to yield (R)-N-(14-azido-3,6,9,12-tetraoxatetradecyl)-4-((8-cyclopentyl-7-ethyl-5-methyl-6-oxo-5,6,7,8-tetrahydropteridin-2-yl)amino)-3-methoxybenzamide as a off-white solid (5.5 mg, 0.00822 mmol, 49.8% yield). The resulting solid was dissolved in DMSO (1 mL) and combined with 9-acryloyl-N-(prop-2-yn-1-yl)-9H-carbazole-3-carboxamide (3.5 mg, 0.0116 mmol), CuBr (2 mg, 0.0140 mmol), THPTA (6 mg, 0.0138 mmol), and water (0.3 mL). After completion (10 minutes), the reaction was purified by reverse-phase HPLC (acetonitrile:water gradient up to 90%) to yield KG5-PEG4-BI2536 as an off-white solid (3.6 mg, 0.00370 mmol, 45.1% yield). A stock solution was created in DMSO, and a portion of the stock solution was set aside for $^1$H NMR characterization. Due to mixing with residual DCM in the NMR tube from cleaning, the portion of the stock that underwent NMR was discarded. $^1$H NMR (400 MHz, DMSO) spectrum is shown below. HRMS (top: observed, bottom: theoretical isotope pattern; M+H, $C_{51}H_{61}N_{11}O_9$+H] m/z theoretical 972.4726, found 972.4709.

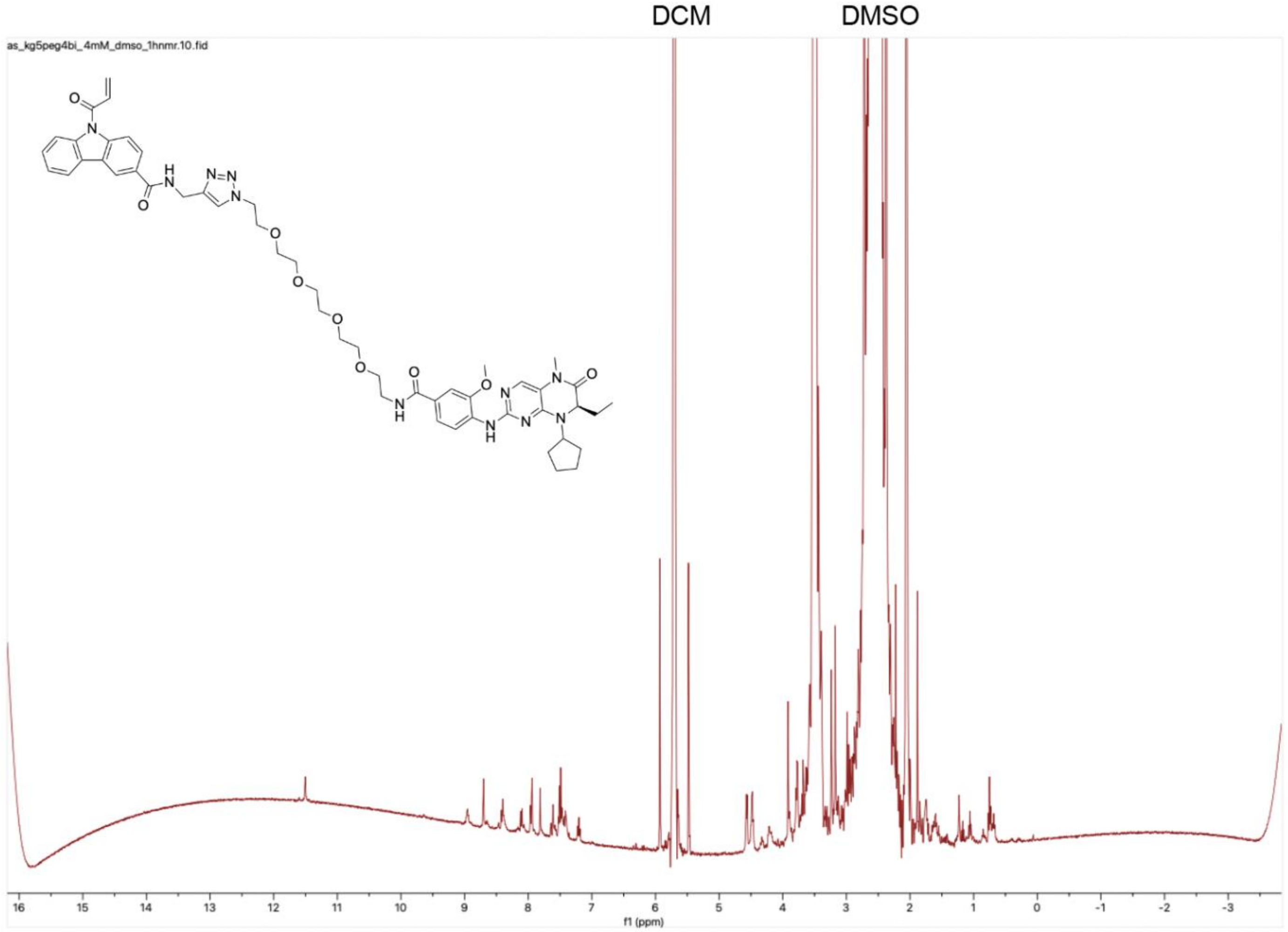

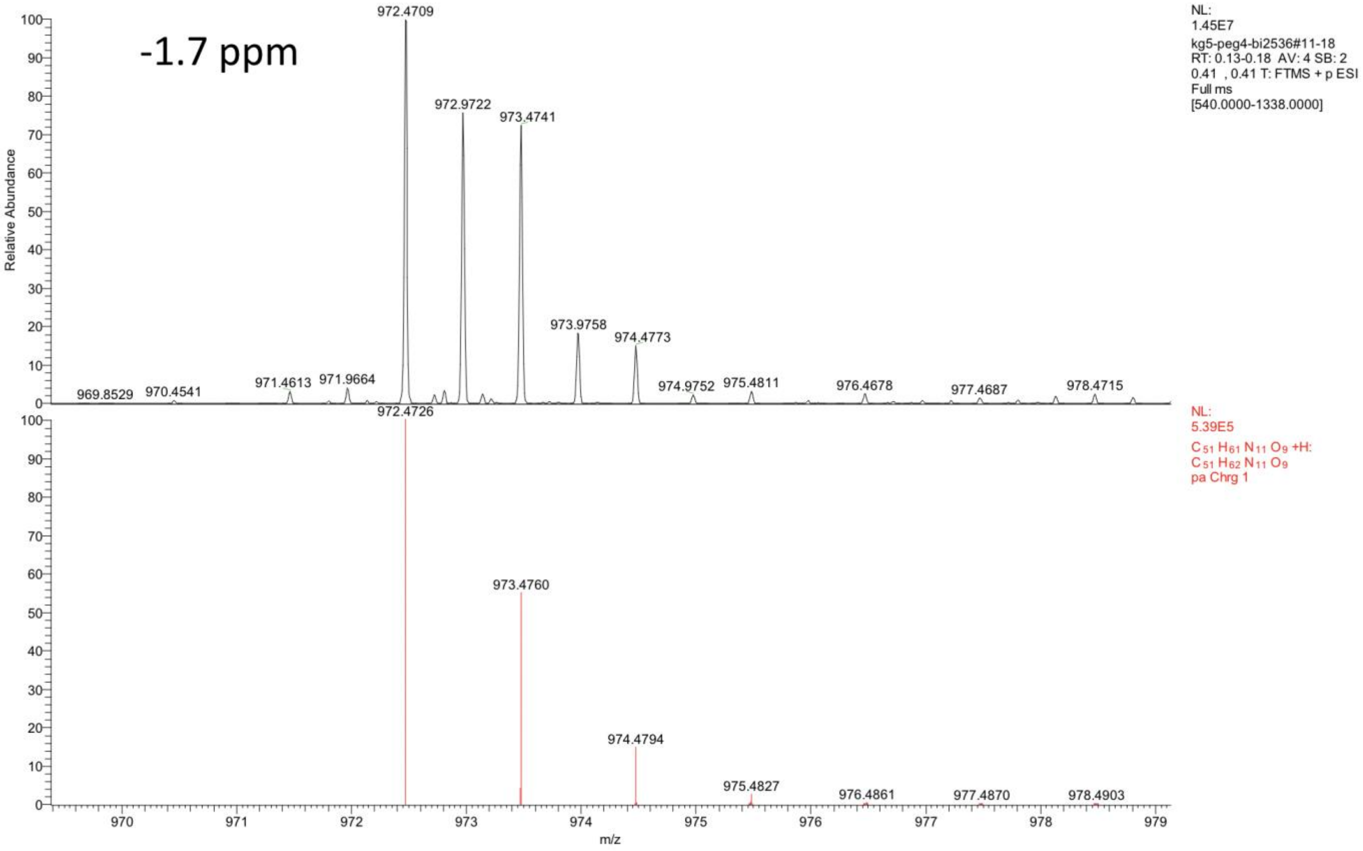


*BI2536-PEG2-Halo (HLDA-131).*

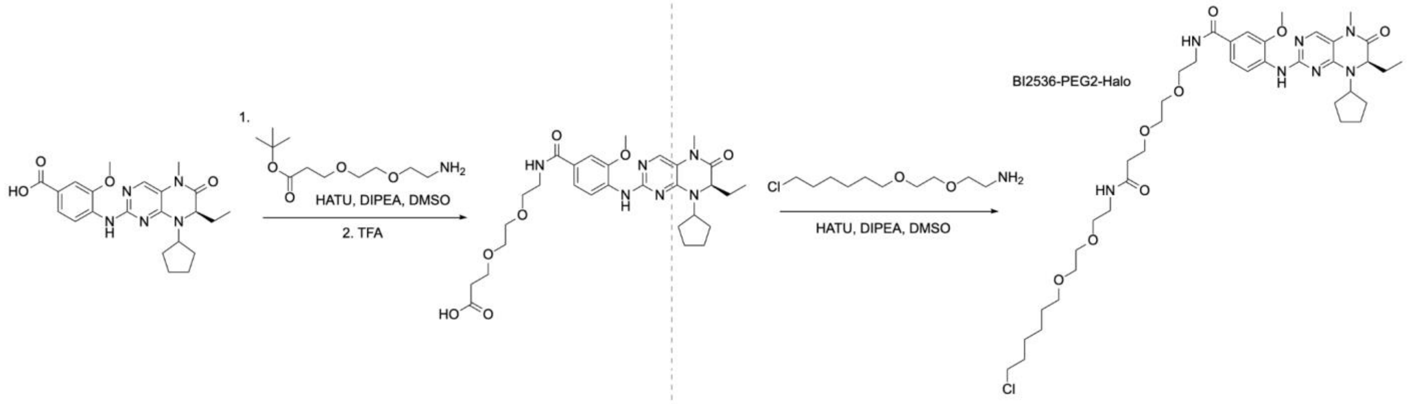


**(R)-N-(22-chloro-9-oxo-3,6,13,16-tetraoxa-10-azadocosyl)-4-((8-cyclopentyl-7-ethyl-5-methyl-6-oxo-5,6,7,8-tetrahydropteridin-2-yl)amino)-3-methoxybenzamide (HLDA-131, BI2536-PEG2-Halo).** To a solution of (R)-4-(8-cyclopentyl-7-ethyl-5-methyl-6-oxo-5,6,7,8-tetrahydropteridin-2-ylamino)-3-methoxybenzoic acid (10 mg, 0.0235 mmol, 1ClickChemistry) in DMSO (1 mL), $NH_2$-PEG2-$CH_2CH_2CO_2$tBu (20 µL, 0.0858 mmol), HATU (12 mg, 0.0315 mmol), and DIPEA (20 µL, 0.115 mmol) were added. After stirring for 30 minutes, trifluoroacetic acid (2 mL) was added and the reaction was stirred for 16 hours. After completion, the reaction was purified by reverse-phase HPLC (acetonitrile:water gradient up to 90%) to yield (R)-3-(2-(2-(4-((8-cyclopentyl-7-ethyl-5-methyl-6-oxo-5,6,7,8-tetrahydropteridin-2-yl)amino)-3-methoxybenzamido)ethoxy)ethoxy)propanoic acid (BI2536-PEG2-acid) as a yellow solid (7 mg, 0.0120 mmol, 51.1% yield). To a solution of the product in DMSO (1 mL), 2-(2-((6-chlorohexyl)oxy)ethoxy)ethan-1-amine hydrochloride (10 mg, 0.0385 mmol, MedChemExpress), HATU (8 mg, 0.0205 mmol), and DIPEA (20 µL, 0.115 mmol) were added to the solution. After completion (30 minutes), the reaction was purified by reverse-phase HPLC (acetonitrile:water gradient up to 90%) to yield HLDA-131 as a yellow solid (5.2 mg, 0.00658 mmol, 54.8% yield). $^{1}$H NMR spectra matched what was previously reported[33]. HRMS (top: observed, bottom: theoretical isotope pattern; M+H, $C_{39}H_{60}ClN_7O_8$+H] m/z theoretical 790.4265, found 790.4254.

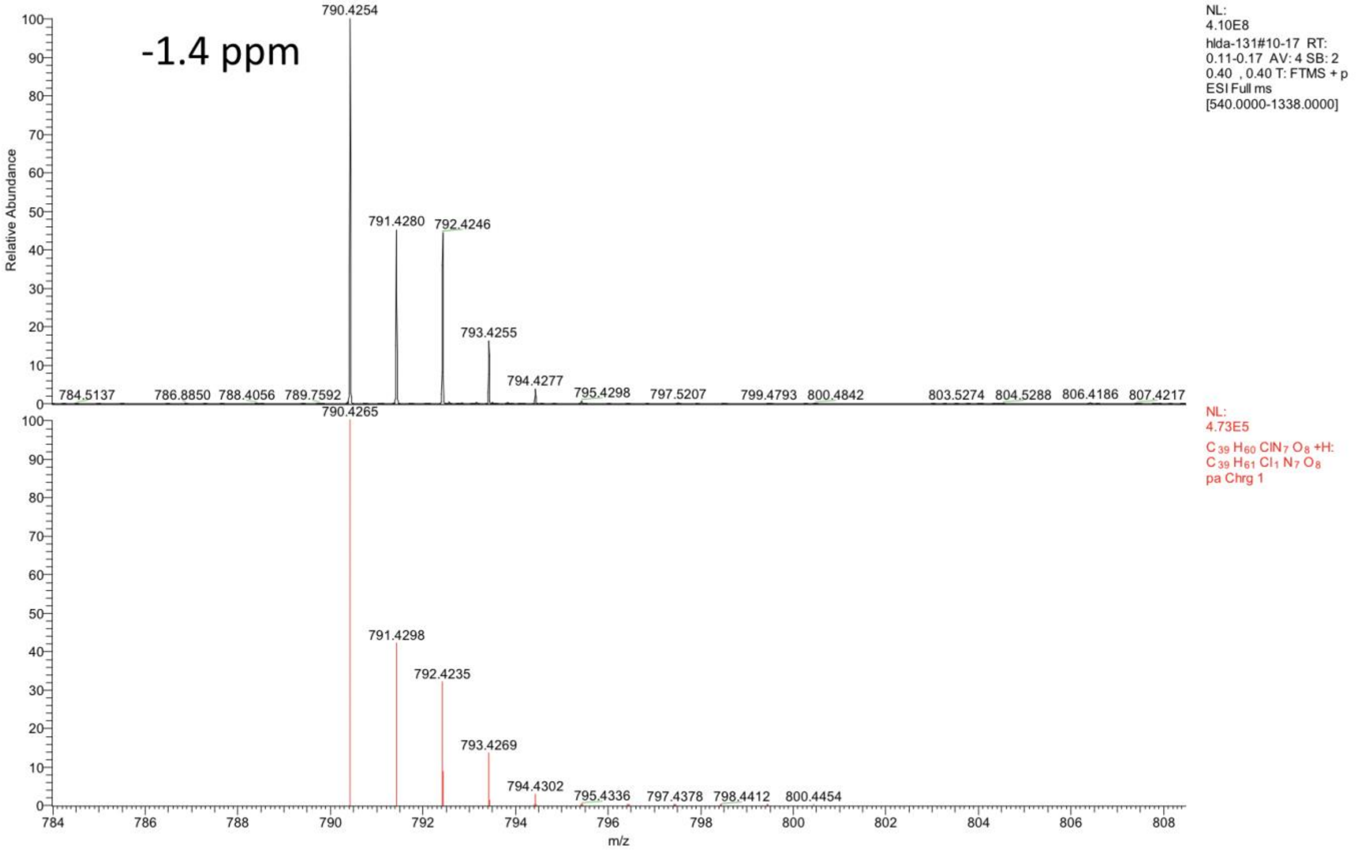


*Adavosertib-PEG2-Halo.*

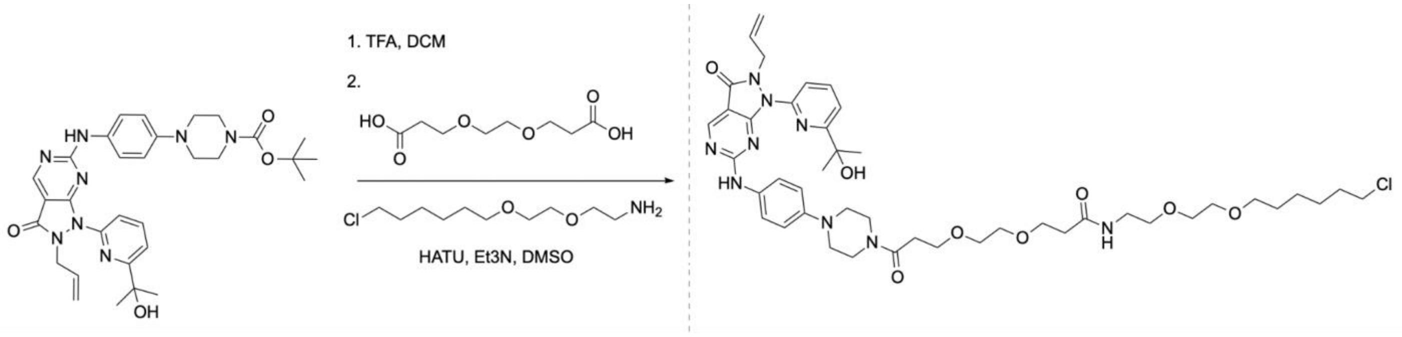


**3-(2-(3-(4-(4-((2-allyl-1-(6-(2-hydroxypropan-2-yl)pyridin-2-yl)-3-oxo-2,3-dihydro-1H-pyrazolo[3,4-d]pyrimidin-6-yl)amino)phenyl)piperazin-1-yl)-3-oxopropoxy)ethoxy)-N-(2-(2-((6-chlorohexyl)oxy)ethoxy)ethyl)propanamide (Adavosertib-PEG2-Halo, Ada-PEG2-Halo).** Adavosertib-NBoc (synthesis described in the PMV6-PEG4-Adavosertib section, 22.2 mg, 0.0379 mmol) was dissolved in DCM (1 mL) and TFA was added (3 mL); the mixture was left to stir for 1 hour and solvents were evaporated under reduced pressure. The residue was dissolved in DMSO (2 mL). PEG2 diacid (7.8 mg, 0.0379 mmol) and 2-(2-((6-chlorohexyl)oxy)ethoxy)ethan-1-amine hydrochloride (9.8 mg, 0.0378 mmol) were added, followed by triethylamine (75 µL, 0.539 mmol) and HATU (31.6 mg, 0.0832 mmol). After completion (30 minutes), the reaction was purified by reverse-phase HPLC (acetonitrile:water gradient up to 90%) to yield adavosertib-PEG2-Halo as a yellow solid (12.2 mg, 0.0139 mmol, 36.7% yield). $^{1}$H NMR (400 MHz, DMSO-d6) δ 8.84 (s, 1H), 8.05 (t, *J* = 7.8 Hz, 1H), 7.88 (t, *J* = 5.6 Hz, 1H), 7.76 (d, *J* = 8.1 Hz, 1H), 7.65 – 7.58 (m, 1H), 7.60 (s, 1H), 6.96 (d, *J* = 8.7 Hz, 2H), 5.76 (s, 2H), 5.67 (ddt, *J* = 16.4, 10.2, 6.0 Hz, 1H), 5.32 (s, 1H), 5.04 – 4.96 (m, 1H), 4.83 (dq, *J* = 17.1, 1.5 Hz, 1H), 4.69 (d, *J* = 6.0 Hz, 2H), 3.69 – 3.55 (m, 9H), 3.48 (tp, *J* = 5.2, 2.7 Hz, 8H), 3.47 – 3.31 (m, 4H), 3.19 (q, *J* = 5.9 Hz, 2H), 3.12 (t, *J* = 5.1 Hz, 2H), 3.06 (t, *J* = 5.2 Hz, 2H), 2.63 (t, *J* = 6.6 Hz, 2H), 2.31 (t, *J* = 6.5 Hz, 2H), 1.75 – 1.63 (m, 2H), 1.50 (d, *J* = 6.8 Hz, 1H), 1.47 (s, 7H), 1.46 (s, 1H), 1.43 – 1.21 (m, 4H). $^{13}$C NMR (101 MHz, DMSO-d6) δ 170.57, 169.26, 168.06, 161.60, 160.95, 156.49, 147.38, 139.27, 132.67, 131.85, 118.73, 116.75, 116.60, 72.79, 70.66, 70.08, 70.05, 69.98, 69.89, 69.58, 67.28, 55.38, 49.80, 49.32, 47.06, 45.81, 45.34, 41.32, 39.00, 36.51, 33.28, 32.48, 30.92, 29.53, 26.59, 25.39. HRMS (top: observed, bottom: theoretical isotope pattern; M+H, $C_{44}H_{62}ClN_9O_8$+H] m/z theoretical 880.4483, found 880.4473.

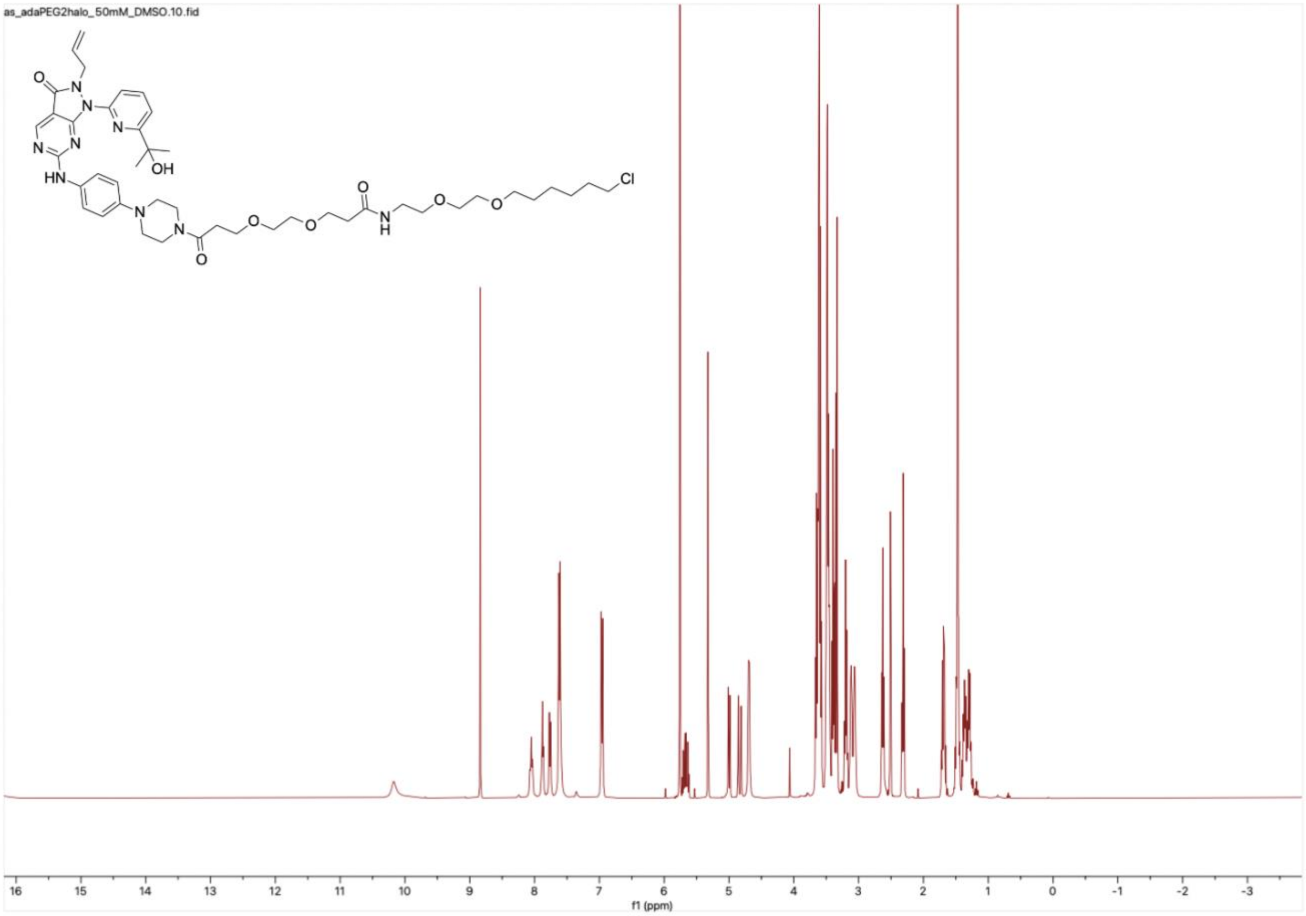

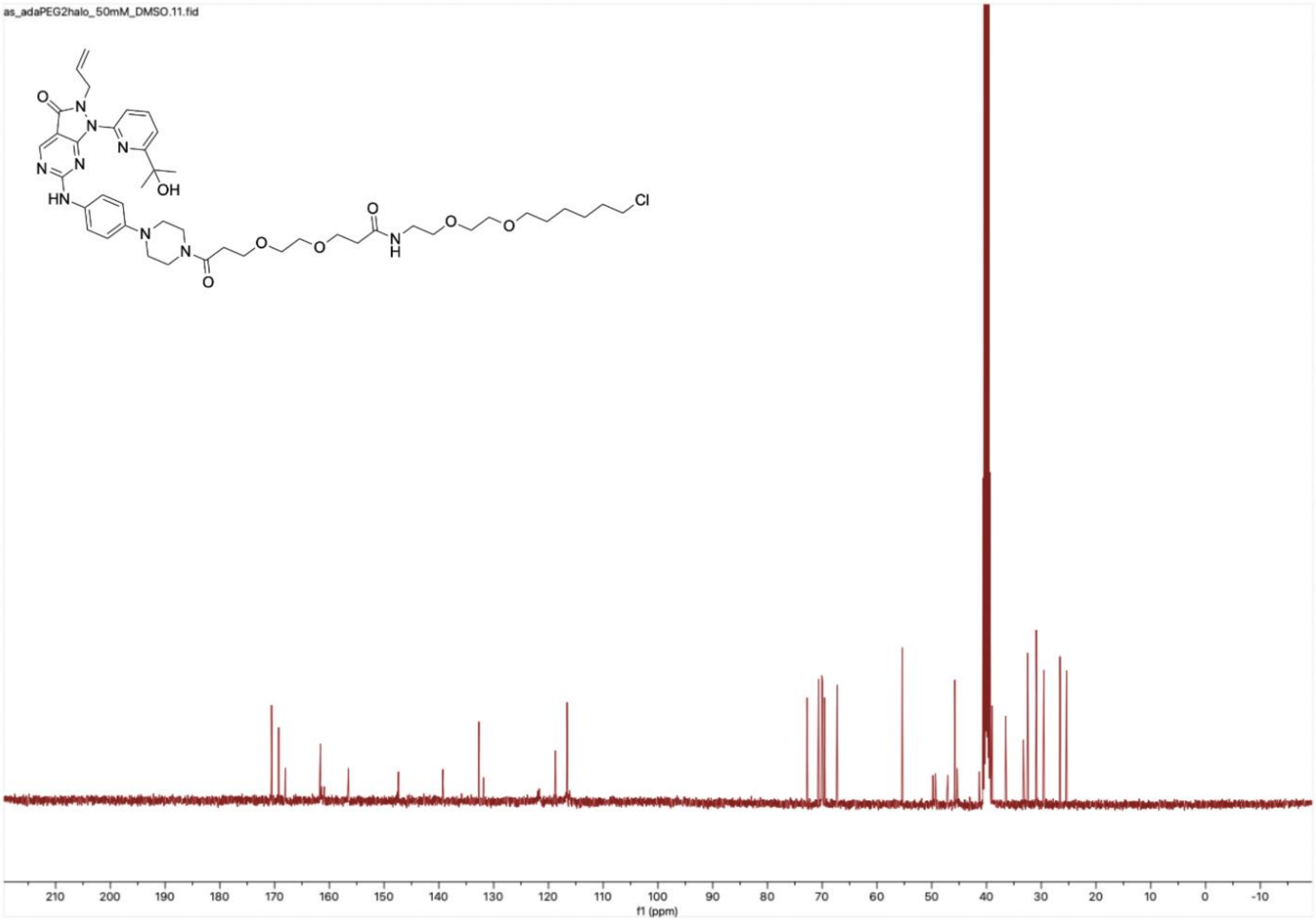
as_adaPEG2halo_50mM_DMSO.11.fid
210 200 190 180 170 160 150 140 130 120 110 100 90 80 70 60 50 40 30 20 10 0 -10
f1 (ppm)

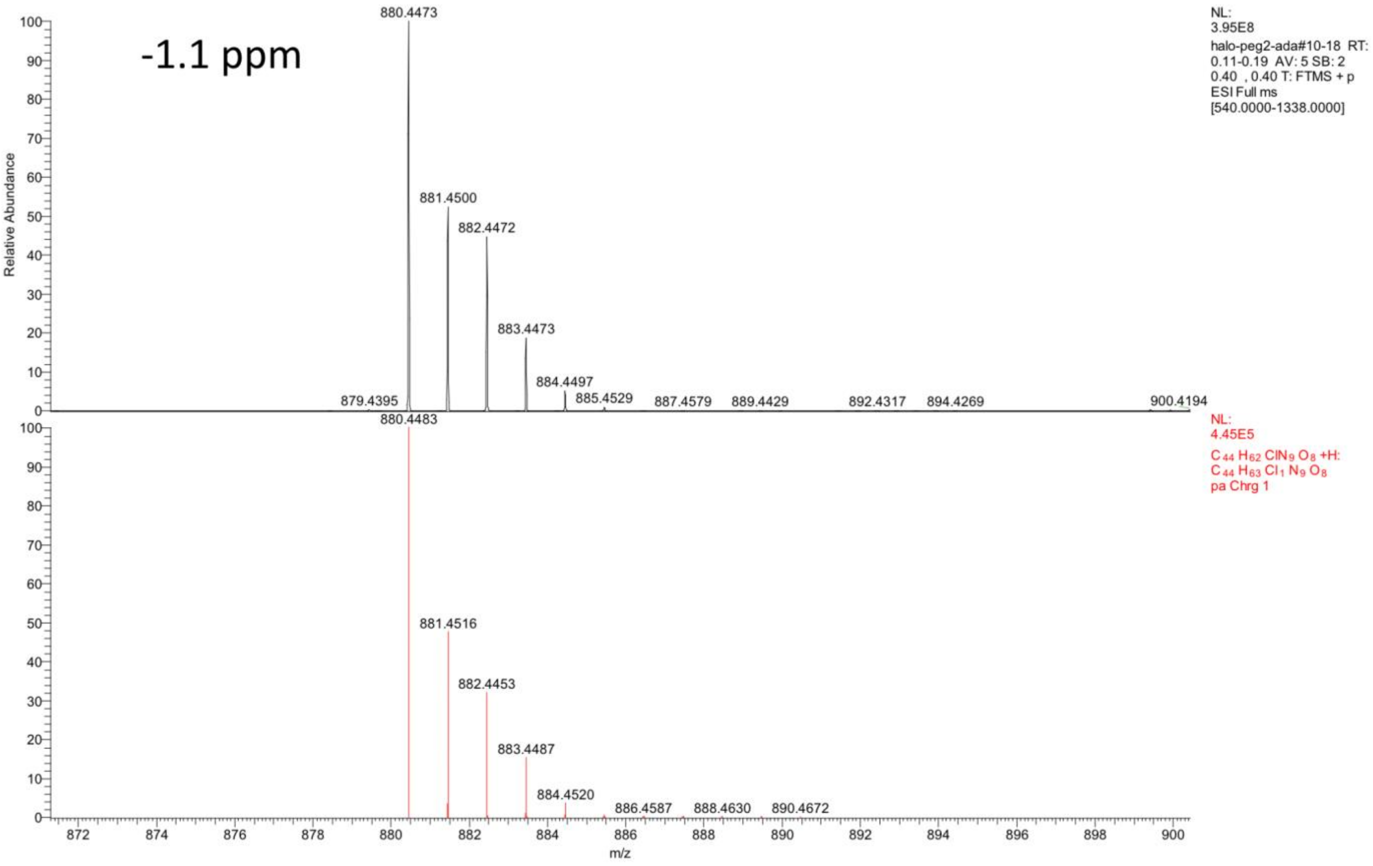

-1.1 ppm
880.4473
881.4500
882.4472
883.4473
884.4497
879.4395
885.4529
887.4579
889.4429
892.4317
894.4269
900.4194
NL:
3.95E8
halo-peg2-ada#10-18 RT:
0.11-0.19 AV: 5 SB: 2
0.40 , 0.40 T: FTMS + p
ESI Full ms
[540.0000-1338.0000]
880.4483
881.4516
882.4453
883.4487
884.4520
886.4587
888.4630
890.4672
NL:
4.45E5
$C_{44}H_{62}ClN_9O_8$ +H:
$C_{44}H_{63}Cl_1N_9O_8$
pa Chrg 1
Relative Abundance
m/z
